# Generating Ensembles of Gene Regulatory Networks to Assess Robustness of Disease Modules

**DOI:** 10.1101/2020.07.12.198747

**Authors:** James T. Lim, Chen Chen, Adam D. Grant, Megha Padi

## Abstract

The use of biological networks such as protein-protein interaction and transcriptional regulatory networks is becoming an integral part of biological research in the genomics era. However, these networks are not static, and during phenotypic transitions like disease onset, they can acquire new “communities” of genes that carry out key cellular processes. Changes in community structure can be detected by maximizing a modularity-based score, but because biological systems and network inference algorithms are inherently noisy, it remains a challenge to determine whether these changes represent real cellular responses or whether they appeared by random chance. Here, we introduce Constrained Random Alteration of Network Edges (CRANE), a computational method that samples networks with fixed node strengths to identify a null distribution and assess the robustness of observed changes in network structure. In contrast with other approaches, such as consensus clustering or established network generative models, CRANE produces more biologically realistic results and performs better in simulations. When applied to breast and ovarian cancer networks, CRANE improves the recovery of cancer-relevant GO terms while reducing the signal from non-specific housekeeping processes. CRANE is a general tool that can be applied in tandem with a variety of stochastic community detection methods to evaluate the veracity of their results.

## Introduction

Elucidating the genetic variants and mechanisms that drive complex disease remains a difficult problem. Instead of being characterized by a single gene or mutation, complex diseases appear to be caused by many changes and perturbations scattered around the gene regulatory network that create a considerable amount of variability in disease susceptibility [1–3]. Network-based methods have therefore become a popular approach to prioritize candidate disease genes according to their role in the network of molecular interactions in the cell [4, 5]. These methods capitalize on the idea that biological networks are composed of “communities” of genes or nodes that work in concert to carry out cellular functions [6–8]. Identifying the differential networks and communities associated with disease tissue can reveal more mechanistic insights than standard approaches like differential expression analysis. Thus, a wide variety of tools have been developed to identify the changes in network edges and network structure that accompany disease onset [9–12].

However, determining the robustness and significance of changes in network structure remains a challenge. Gene regulatory networks are often inferred from transcriptomic data using imperfect inference tools, with no easy way of assessing their underlying variance [13–16]. Moreover, community detection algorithms can lead to a wide range of possible solutions corresponding to local optima of the fitness function [17–19]. Two types of approaches have been used to judge the quality of network communities in the past: consensus clustering and statistical significance [15, 16, 20–22]. The consensus approach combines multiple solutions from the optimization algorithm to find the most likely assignment of genes to communities [14, 20]. However, this method leads to a significant reduction of resolution that impairs biological interpretation of the discovered communities [23]. Alternatively, the statistical significance of individual communities can be estimated by comparing them with structures formed in random networks with the same degree characteristics as the original network [13, 24, 25]. These random networks are typically created using generative models.

In the present study, we set out to evaluate the significance of disease-specific modules, and the genes within them. We first reconstructed transcriptional networks using state-of-the-art methods that integrate transcription factor (TF) motifs with gene expression data, and then optimized a modularity-based score to identified differential modules more active in disease conditions than in matched controls [26]. Other approaches for differential network analysis could be used, including DiffCoEx, DINA, DNA, and Diffany [9–12], but these methods are limited to either identifying individual correlation-based edges or examining pre-defined gene sets and network features, making them less generalizable to multiple types of questions and networks [9–12, 27]. We then applied several popular network generative models – configuration model [16] and stochastic block model (SBM) [35] - to simulate the null distribution of the network and estimate the significance of the differential modules.

We found that the available network generative models cannot realistically simulate the characteristics of real transcriptional networks. The configuration model assumes every edge is independent and exponentially distributed. The SBM assumes latent structures (i.e., blocks) with pre-specified inter and intra-block edge densities that can be independently distributed between all genes and regulators in the same block. Although these models are mathematically elegant, the assumptions of edge independence and exponentially distributed edge weights are not compatible with gene regulatory networks, where each TF is not likely to independently regulate all genes, but only those that have a matching, accessible binding motif in their promoter. Therefore, we need a new, computationally efficient generative model that properly accounts for the known characteristics of gene regulation [28–31]

Here we present a new algorithm for network generation called <u>C</u>onstrained <u>R</u>andom <u>A</u>lteration of <u>N</u>etwork <u>E</u>dges (CRANE). CRANE produces ensembles of unipartite and bipartite weighted networks with fixed node strengths that resemble gene regulatory networks. CRANE can be used to generate realistic null distributions that help evaluate the importance of each individual gene or regulator in a disease module. To demonstrate the utility of CRANE, we apply it to simulated networks, as well as transcriptional networks derived from angiogenic ovarian tumors and hormone receptor-positive breast cancers. In simulations, CRANE performs better than all popular network generative models in finding the “true” disease module. When applied to breast and ovarian cancer networks, several generative models are able to improve recovery of some cancer-associated gene modules, but CRANE is the only method that consistently reveals biological insights across multiple networks and conditions while also reducing background noise from non-specific housekeeping processes. Although we have focused its application on disease modules, the core engine of CRANE is a general tool that can be used to generate ensembles of graphs consistent with gene regulatory networks for application in a number of contexts.

## Materials and Methods

### Data preprocessing and network inference

Batch-normalized ovarian PanCancer TCGA RNA-seq values were downloaded from cBioPortal. Low-expressing genes were removed by keeping only genes with at least 1 CPM in at least half of the total samples using the R package edgeR (v3.26.5) and processed with the voom function within the R package limma (v3.38.3) using TMM normalization. Angiogenic (n=124) and non-angiogenic (n=166) tumors were grouped as described in [32]. Preprocessed METABRIC breast cancer expression data was downloaded from cBioPortal [33, 34], along with estrogen receptor negative (ER-; n=445) and estrogen receptor positive (ER+; n=1449) status as measured by immunohistochemistry. Expression data from each subtype was integrated with transcription factor binding sites using the network inference algorithm PANDA (Passing Attributes between Networks for Data Assimilation) to create subtype-specific regulatory networks [35]. Subsampled networks were inferred by taking a random subset of 50 subjects without replacement from the gene expression of each respective subtype. The prior network of binding sites for 730 TFs was defined as the occurrence of the corresponding motif in a [-750,+250] bp window around the transcription start site [36].

The following formula was applied for analyses requiring exponentially transformed PANDA edge weights [36]:

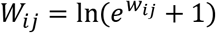

### CRANE algorithm

CRANE takes a weighted bipartite network as input and provides a perturbed version of that network as output. We first compute the strength of all the nodes in the network as a reference. As an optional step, noise can be added to the reference strength by adding normally distributed random numbers with mean of 0 and standard deviation determined from subsampled networks. Given m is the total number of TFs and n is the total number of genes, *A_ij_* is the m x n adjacency matrix of the reference network where rows (TFs) and columns (genes) are ordered randomly. We create an empty m x n adjacency matrix *B_ij_* that will become the CRANE perturbed network. The first row (first TF) of *B_ij_* is initialized with edge weights from the first row of *A_ij_*. Then for each *TF_l_*, where *l* = [1, …, *m* – 1], we perturb the *l*th row *B_lj_* by adding normally distributed random numbers with mean of 0 and standard deviation matching the original edge weights from the same *TF_l_*, *A_lj_*. The *α* parameter determines the magnitude of the perturbation. The *B_lj_* edges are then normalized by 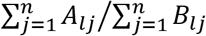 to ensure the TF strength in *B_lj_* is equal to *A_lj_*. Then we determine *B*_*l*+1,*j*_ (next TF) by computing 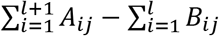. This allows *B_ij_* to keep the gene strength distribution equal to *A_ij_*. After the initial *B*_*l*+1,*j*_ have been determined, we check if any edge weights within *B*_*l*+1,*j*_ fall outside of the global maximum or minimum of the original edge weights in *A_ij_*. For any values in *B*_*l*+1,*j*_ greater than *max*(*A_ij_*) edge weight, we add the difference in value between *B*_*l*+1,*j*_ and *max*(*A_ij_*) to the corresponding *B*_1*j*_. For any values *B*_*l*+1,*j*_ less than *min*(*A_ij_*) we subtract the difference in value between *B*_*l*+1,*j*_ and *min*(*A_ij_*) and add to the corresponding *B_lj_*. Then the modified edge weights in *B_lj_* are normalized to maintain the correct TF strength and a new set of *B*_*l*+1,*j*_ are computed. We repeat the correction process until all values are within the range of *A_ij_*. We then iteratively move down each row in the matrix and repeat the procedure until the matrix is filled.

A more detailed description of CRANE can be found in the supplementary methods. Additionally, a unipartite version of CRANE is also available for use. CRANE is freely available as an R package at https://github.com/PadiLab/CRANE.

### Configuration Model

To fit the configuration model to the PANDA network, we exponentially transformed z-score edge weights to positive weights using the formula given in the “Data preprocessing and network inference” section. Based on the fact that PANDA is a fully connected graph, the configuration model could be written as described in [37],

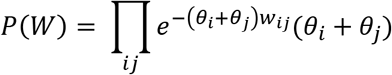

where the *θ* parameters are the Lagrange multipliers that need to be estimated. The Maximal Likelihood (ML) function then constrains the *θ* parameters,

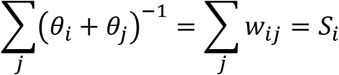

where *S_i_* represents the strength of node *i*. We used Barzilai-Borwein spectral methods for solving the ML system of equations directly using the R package *BB* (v 2019.10.1) [38].

### Stochastic Block Model

Fitting a Stochastic Block Model (SBM) to the PANDA network is very time consuming [39]. To efficiently test the performance of SBM, we introduced a strong assumption of equivalence between modularity optimization and SBM maximum likelihood [40]. Thus, the community structure detected by COmplex Network Description Of Regulators (CONDOR) algorithm [8] – a modularity maximization method for weighted bipartite networks – was used to approximate the block structure in the SBM. We assumed a normal distribution for edge weights as the PANDA network has z-scored edge weights. The parameters for every block could be estimated directly using the sample mean and sample variance of the corresponding edge bundles.

### Permutation of network edges

The permutation method takes a weighted bipartite network as input and provides a network with randomly shuffled edge weights as output. To retain as much of the regulatory information as possible, the edges in the network were first divided into motif positive and motif negative based on the prior network of binding sites for 730 TFs. The network was clustered using CONDOR [8]. Finally, the inter- and intra-community edge weights were grouped together by motif status and randomly shuffled.

### Using CRANE and other methods to estimate node significance

We performed ALPACA as described in [26] to identify modules that are different between the phenotype-positive network (network representing our biological process of interest) and its matching baseline network to obtain the module assignment *K_node_* and its differential score *S_node_* for each nodes. In order to test the significance of *S_node_*, we used CRANE (α = 0.1), configuration model, SBM, and permutation method to construct the null distribution using the following steps. (i) Use CRANE or other methods to generate a perturbed network starting from the baseline network. (ii) Compute a null differential modularity matrix *D_ij_* by comparing the newly generated network to the original baseline network. (iii) Score each node according to its contribution to the differential modularity of the module that it belongs to in *K_node_*. We repeated (i) through (iii) for the desired number of times (e.g. 1000) to generate a null distribution T for each node (*T_node_*). As a final step we computed the p-value for the differential modularity score (*S_node_*) by assuming *T_node_* follows a normal distribution:

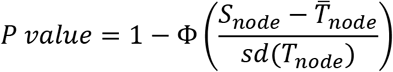

For GO term enrichment analysis, we ranked the target genes in each module by their p-value. We evaluated the top 25, 50, 75, etc., up to 500 core genes from each module to identify functional enrichment at different cutoffs. All enriched GO terms across all cutoffs were included in the final result for each module. GO term enrichment was calculated using the R package GOstats (v2.54.0), with the following parameters: the gene universe is defined to be the set of all possible target genes in the initial networks, and the p-value calculation is conditioned on the GO hierarchy structure. In each module, the p-values were adjusted for multiple testing using the Benjamini-Hochberg method.

### Evaluating CRANE on simulated networks

To simulate disease modules, we first took a random subset of 50 subjects out of 445 subjects from the estrogen receptor negative METABRIC breast cancer expression data and constructed a baseline PANDA network. We then inserted high edge weights (edge weight = 5) between randomly selected TFs and genes to create a disease phenotype network. The new module consisted of between 3 and 20 transcription factors, and five times as many genes as transcription factors. We then compared these disease networks to a second baseline network inferred from a different random subset of 50 subjects from the ER-breast tumors. We applied a panel of methods, including ALPACA, consensus ALPACA, CRANE (α = 0.1), configuration model, SBM, and permutation method and evaluated the results of each method by comparing the ranks of true positives (the known genes in the disease module) against a background consisting of target genes not in the disease module. We used Kolmogorov-Smirnov and Wilcoxon tests to look for significant differences in the distribution of the ranks. Both tests gave similar results, and so in the figures, we present the Wilcoxon p-values.

### Consensus clustering of ALPACA

ALPACA was downloaded from https://github.com/PadiLab/ALPACA. In order to generate consensus clusters, we first repeated ALPACA 1,000 times on the same pair of transcriptional networks, as described in [26] but with the nodes ordered randomly in each iteration of the Louvain algorithm. We combined the 1,000 resulting partitions to create an n x n consensus matrix *C* whose entry *C_ij_* indicates the number of partitions in which vertices *i* and *j* of the network were assigned to the same cluster, divided by the total number of partitions (1,000). For the final step, we applied the Louvain algorithm (igraph R package v1.2.4.1) on C to find the consensus cluster membership for each node [18, 41].

### Computing NMI

For a given weighted bipartite network, the CONDOR algorithm was used to detect the community structure of the network [4, 8]. Using CONDOR community assignments as input, the normalized mutual info (NMI) score between two networks was computed using the “compare” function in the R package igraph (v1.2.4.1) [41, 42].

## Results

### Consensus clustering of differential modules

Most community detection methods are stochastic and can result in multiple near-optimal solutions [17–19]. This ambiguity in the network clustering result can muddle the biological conclusions derived from regulatory networks. We have previously shown that comparing condition-specific networks by maximizing the change in modularity (ALPACA) can uncover biological processes and communities that could not be detected by differential expression alone. ALPACA computes the differential modularity score to measures the difference in edge density of modules in the “unhealthy” network to the expected density in a matched baseline “healthy” network, and optimizes this score to determine the network communities [26]. However, like other community detection methods based on score maximization, ALPACA results in multiple solutions and does not provide an estimate of the biological significance of the detected alterations. Since consensus clustering is an effective way of finding stable communities among multiple local optima [20], we integrated consensus clustering with ALPACA to see whether the most frequently found modules are enriched with genes responsible for driving the biological phenotype of interest.

We first constructed transcriptional regulatory networks by integrating known human transcription factor (TF) binding sites with gene expression data from angiogenic and non-angiogenic ovarian cancer using the Passing Attributes Between Networks for Data Assimilation (PANDA) algorithm [35]. We then set out to directly compare the modular structure of a baseline (non-angiogenic) network to an altered network (angiogenic) using ALPACA. We merged one thousand partitions from individual ALPACA solutions to generate a consensus co-membership matrix (see Materials and Methods for details). For the final step, we applied the Louvain method to the consensus matrix to determine the consensus community assignment [18]. Consistent with previous observations in the literature, we found that consensus clustering led to a significant loss of resolution (Figure 1) [14, 20], and the inability to detect more specific disease pathways with richer biological interpretations [23].

**Figure 1.**
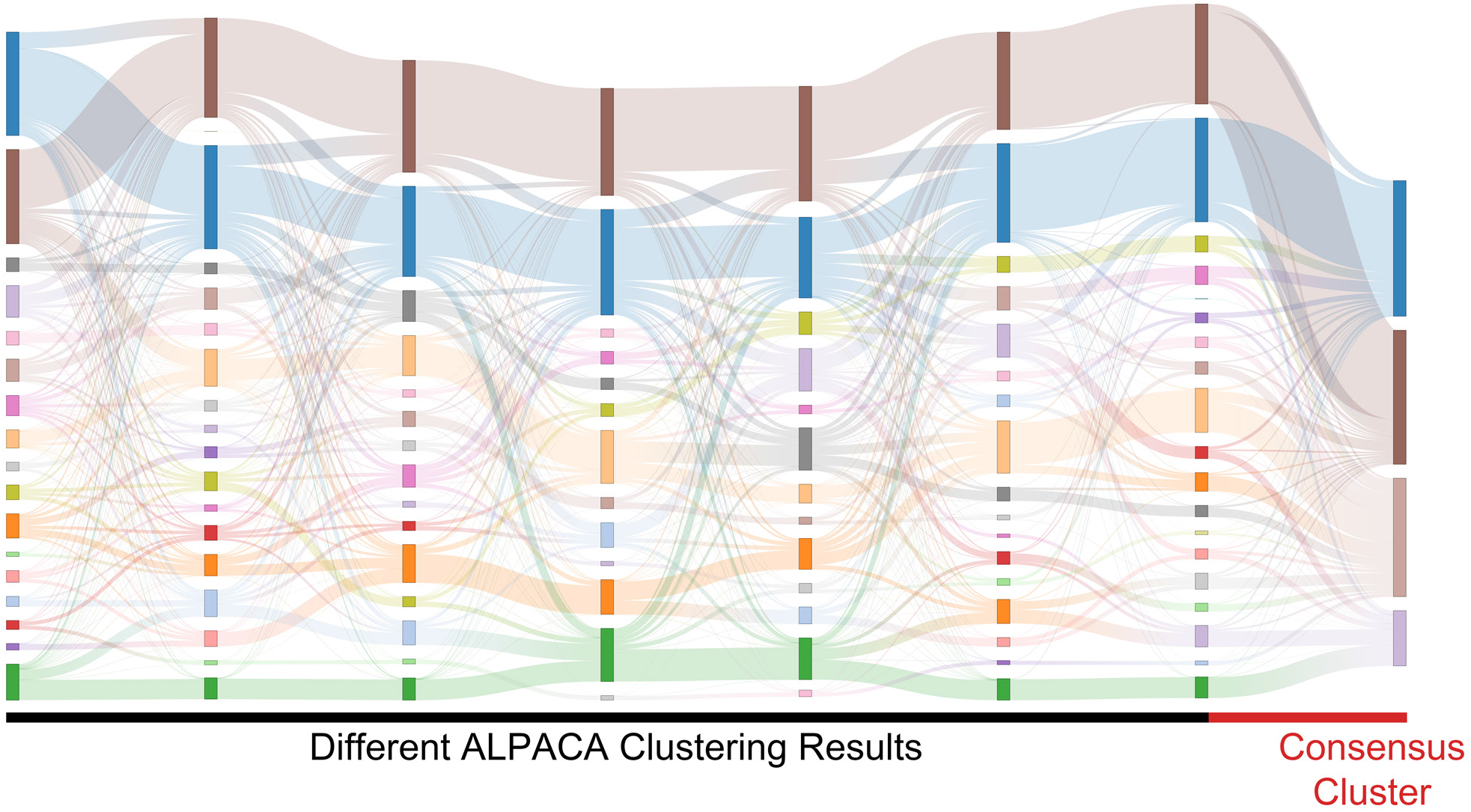
Consensus approach to network clustering leads to decrease in resolution. Sankey plots showing community assignments from seven separate runs of ALPACA on angiogenic vs. non-angiogenic ovarian cancer networks. Each column represents an ALPACA solution with the far-right column showing the result from consensus clustering of 1,000 different ALPACA solutions. The height of each box and ribbons indicates the size of each module and the number of shared nodes between corresponding modules respectively.

### Constrained Random Alteration of Network Edges (CRANE): new method for sampling weighted networks

Because the consensus approach leads to an unavoidable loss of community resolution, we searched for alternative methods for measuring module robustness. We looked at leading generative models to see whether we can simulate the natural variance of gene regulatory networks to test the significance of each module. We chose the configuration model [37] and the stochastic block model (SBM) [39] as they both have rigorous mathematical descriptions, mimic features of biological networks, and are two of the most commonly used generative models. To evaluate each model, we started with two breast cancer subtype-specific networks and computed the null distribution of the differential modularity score (DMS) of the genes using each generative model (Figure 2). We chose to use breast cancer expression data as this cancer has been extremely well-studied and provides enough data to create a “gold standard” distribution of networks by subsetting the data (n=50 for each subsample) and constructing multiple networks corresponding to the same biological condition. We found that both generative models failed to accurately recapitulate the true DMS variance computed from the subsampled networks (Figure 3A). We hypothesized that this is because general-purpose network models ignore the boundaries set by the TF motif prior, leading to an overestimation of the edge weight variance [37, 39]. In order to accurately model transcriptional regulation, the regulatory network must integrate TF binding information either from motif or epigenetics data [43].

**Figure 2.**
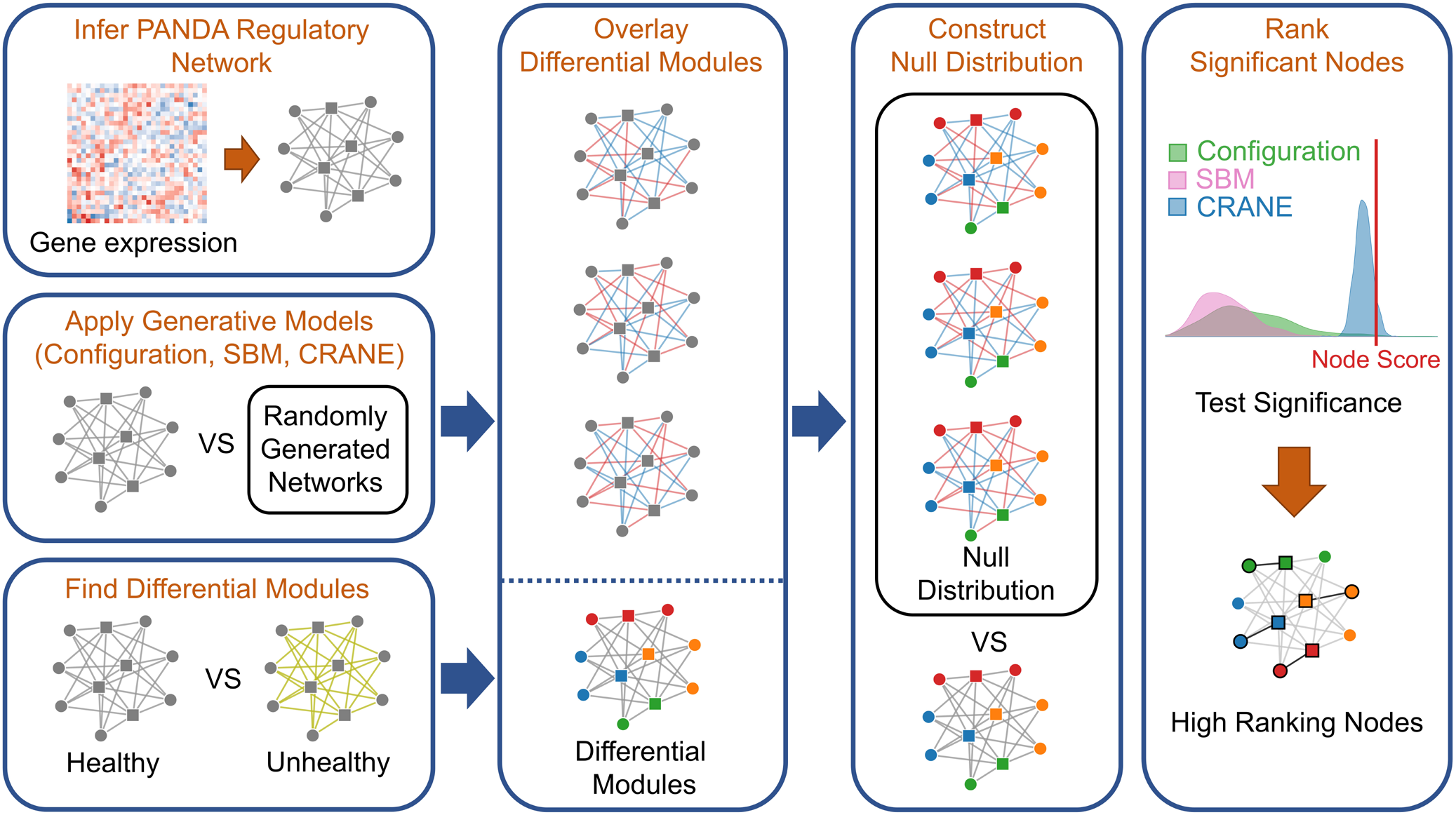
Workflow for applying network generative models to rank genes in disease modules. We first construct transcriptional networks representing the “healthy” and “disease” networks by integrating known TF binding sites with gene expression data. We then use ALPACA to identify putative disease modules and compute the differential modularity scores (DMS) for each node. We construct the null distribution of the DMS by comparing healthy networks to randomly generated networks using the configuration model, SBM, permutation, or CRANE. The p-value is calculated by comparing the true node DMS to each null distribution.

**Figure 3.**
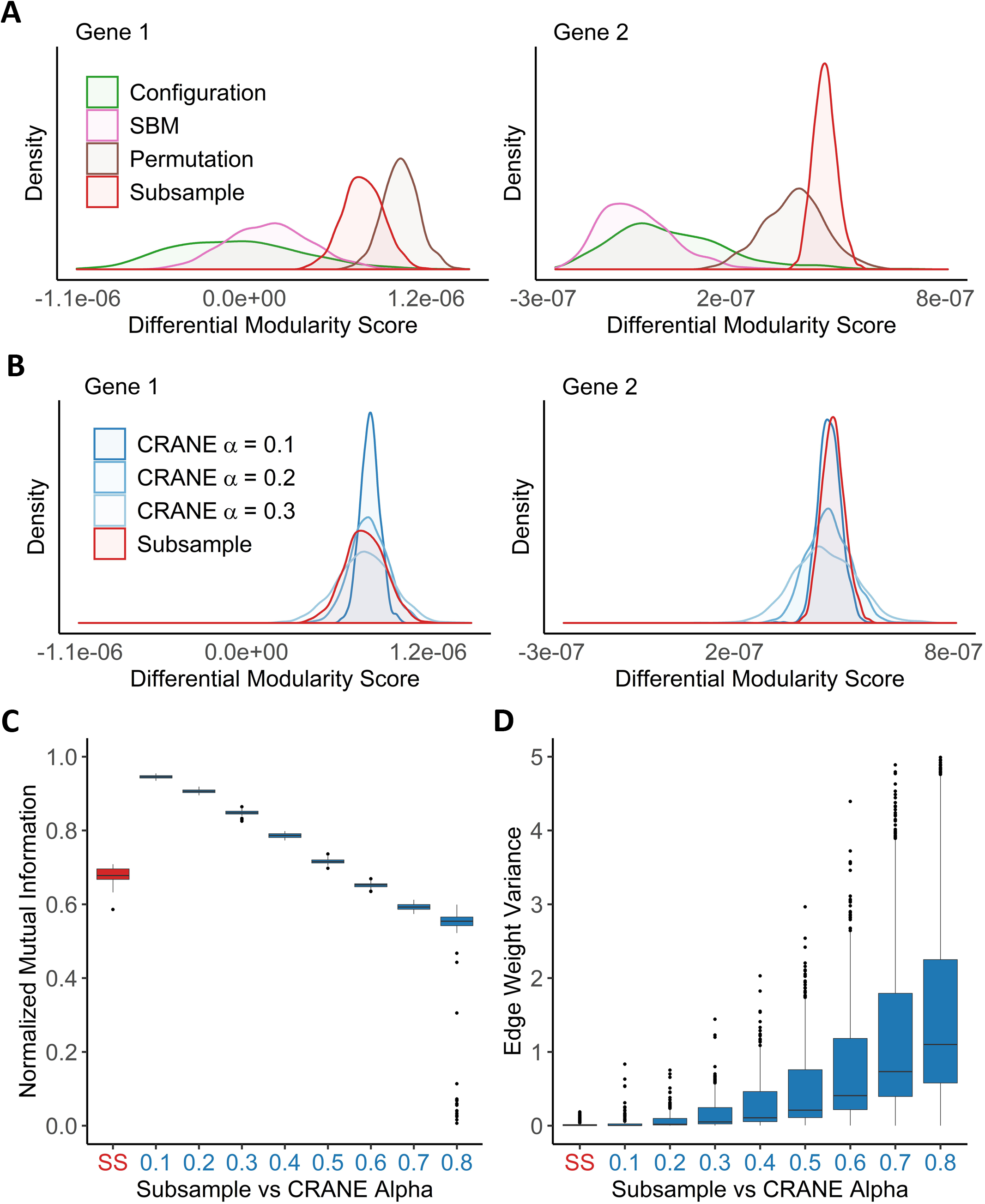
CRANE can generate networks that resemble subsampled data while maintaining control of key network properties. Using the breast cancer transcriptional network as a reference, network ensembles were generated using configuration model, SBM, permutation, and CRANE. As a “gold standard” comparison, PANDA networks were inferred by subsampling (n=50) the gene expression data without replacement from the estrogen receptor positive subtypes. **(A, B)** Density plots showing the null distribution of the differential modularity score (x-axis) computed using different methods for two example genes. **(C, D)** The boxplots show the impact of dialing the alpha parameter in CRANE on **(C)** community structure and **(D)** edge weight variance, as compared to subsampled (SS) networks. **(C)** Plot showing the normalized mutual information (y-axis) between the reference network and CRANE-generated networks for different values of alpha (x-axis). **(D)** The edge weight variance (y-axis) among subsampled or CRANE-generated networks at different values of alpha (x-axis).

Therefore, we developed a new algorithm, Constrained Random Alteration of Network Edges (CRANE), that samples networks with fixed node strengths while retaining the underlying structure defined by prior information from TF binding patterns. We reasoned that fixing the node strengths will help the newly generated networks incorporate the motif information from prior networks and create more proportional variance in edge weights. Moreover, sampling networks with matched node strength would help reduce bias from promiscuous hub TFs and genes that seed modules with disease-agnostic housekeeping processes. To accomplish this, CRANE perturbs each edge weight by adding normally distributed values centered at zero with standard deviation learned from the original network, to simulate the variance arising from normalized gene expression data and to maintain the constraining effect of the TF binding prior. The magnitude of each perturbation can be adjusted by a strength parameter, *α*. CRANE then iteratively redistributes any unrealistically large or small edge weights until they are confined within the range of the original network while the node strengths are kept the same (see Materials and Methods for details).

Compared to the configuration model and SBM, CRANE is better able to mimic the null distribution arising from the “gold standard” subsampled networks (Figure 3A, B). Because CRANE induces small perturbations to the existing edges, the mean of the DMS distribution remains in close proximity to the “gold standard” while other generative models have large deviations across multiple moments of the distribution. In addition, we examined the influence of CRANE on network community structure. We applied the algorithm, COmplex Network Description Of Regulators (CONDOR), for community detection in weighted bipartite networks [8] and used the normalized mutual information (NMI) to compare CRANE-generated networks against the original unperturbed network [42]. This demonstrated that increasing the parameter *α* leads to decreasing NMI score (or similarity) of CRANE-perturbed networks (Figure 3C) and increases edge weight variance (Figure 3D). The *α* parameter can therefore be used to mimic the distribution of the subsampled networks (Figure 3B).

### Evaluating CRANE using simulated networks

We set out to test CRANE in settings that resemble real weighted biological networks, but where we can readily control the modularity. To simulate the effect of measurement noise, we created two independent sets of randomly subsampled (n=50) gene expression data from the same baseline condition, estrogen receptor negative (ER-) breast cancer (BC), and used them to infer two gene regulatory networks, BCN1 and BCN2. Keeping BCN1 as the baseline network, we created an artificial disease module in BCN2 by increasing the edge weights between randomly selected subsets of transcription factors and genes, ranging from 3 to 20 TFs in size, and five times as many genes. We then applied a large panel of methods – namely, ALPACA, consensus clustering, random edge weight permutation, SBM, CRANE, and the subsampling “gold standard” – to identify the most significant differences in network structure. The Wilcoxon rank-sum test was used to evaluate how significantly each method recovered the true disease module. In order to include the configuration model in this panel, we also performed a second test after applying an exponential transformation on the network edge weights, since the configuration model requires positive edge weights [16].

We found that, although ALPACA by itself can successfully recover added modules of size greater than 48 nodes (Figure 4A), CRANE was able to dramatically improve performance. In general, ALPACA demonstrated good performance in the simulations. Consensus clustering improved performance in recovering a single added module, but embeds the artificial module within a much larger community, reducing resolution (Figure 4B). In contrast, CRANE exhibited superior performance over ALPACA in recovering module sizes greater than 36 while retaining high resolution. CRANE was also more successful than random edge weight permutation, the configuration model, and the SBM. As expected, the “gold standard” subsampled distribution performed best out of all the methods. We observed a similar trend in performance whether or not the exponential transformation was applied to the network edge weights (Supplementary Figure 1).

**Figure 4.**
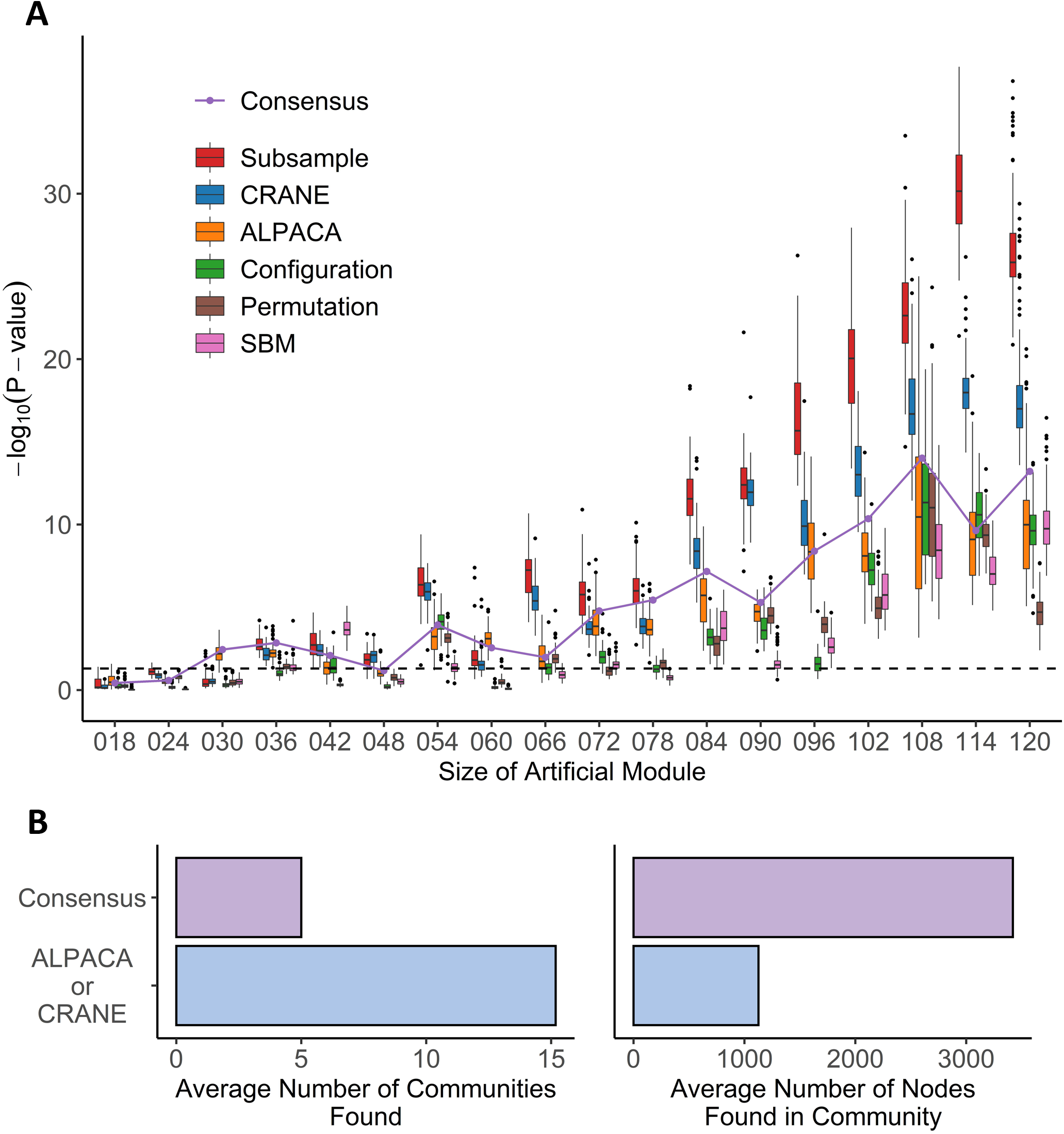
Performance of seven methods on identifying artificial modules in simulated disease networks. **(A)** Box plot shows performance of each method – subsample, CRANE, ALPACA, configuration model, permutation, or SBM – on network simulations with exponentially transformed edge weights. P-values (y-axis) for various sizes of artificial modules (x-axis) are computed using Wilcoxon rank-sum test. **(B)** The boxplots show the total number of modules and module size across all simulation trials for ALPACA versus consensus clustering.

### Evaluating CRANE on biological data

To determine if CRANE can be used to increase the detection of network alterations in complex diseases, we applied CRANE to real biological data. Since there is no “ground truth” dataset for disease modules in transcriptional networks, we employed two well-understood disease processes for which the resulting modules would be easy to interpret: angiogenesis in ovarian cancer, and estrogen response in breast cancer.

#### Ovarian cancer

Ovarian cancer is one of the leading causes of death among women in the developed world [44–46]. Ovarian cancer is divided up into many histologic subtypes based on cellular origin, pathogenesis, molecular alterations, and gene expression profile [46]. For instance, an angiogenesis gene signature can categorize ovarian cancer patients into a poor-prognosis subtype [47]. To test CRANE on angiogenic ovarian cancer, we first applied PANDA to infer human ovarian cancer gene regulatory networks from Pan-Cancer TCGA RNA-seq data. The normalized RNA-seq expression data was classified into 124 angiogenic and 166 non-angiogenic ovarian cancer tumors as described in [32]. We analyzed the set of core genes in differential modules uncovered by each method, and evaluated their functional enrichment for biological processes.

We first checked the performance of consensus clustering compared to ALPACA to see how the reduction in community resolution would impact the biological interpretation. Consistent with our previous work, ALPACA discovers finer community structure enriched for GO terms that are specific to the angiogenic ovarian cancer phenotype such as “blood vessel development” and “cardiovascular system development” (Figure 5A) [26]. In comparison, consensus-ALPACA results in loss of community resolution (Figure 5B), which in turn leads to the loss of the more specific GO terms. Instead, communities are enriched for general processes such as “RNA splicing,” “monoubiquitinated protein deubiquitination,” “translation initiation,” and “ribosome biogenesis” (Supplementary Table 1).

**Figure 5.**
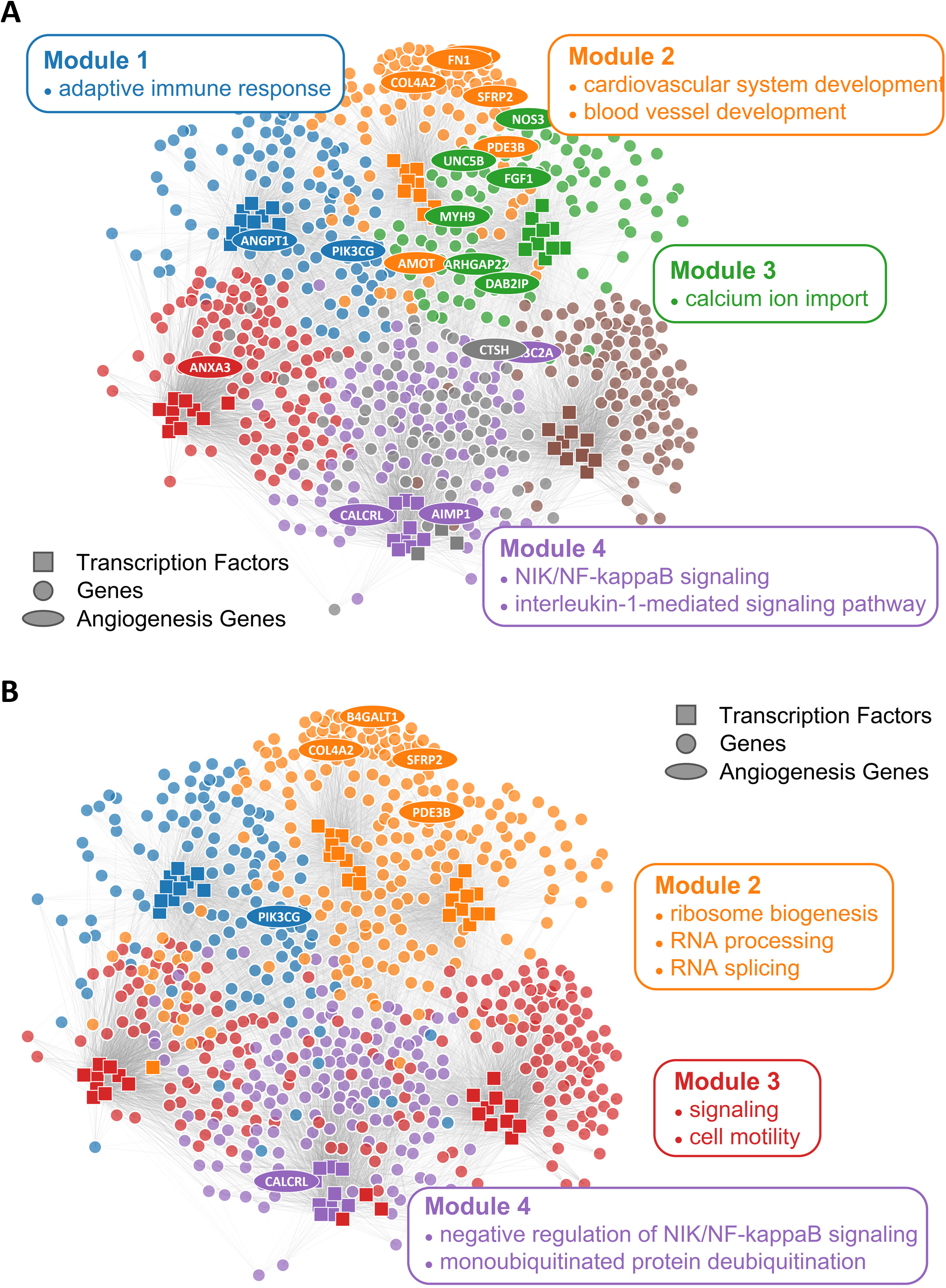
Comparison of ALPACA modules and consensus clustering in angiogenic ovarian tumors. Network with **(A)** ALPACA solution has seven modules while **(B)** the consensus method results in four modules. Top 10 core TFs and 100 core genes were extracted from each module based on DMS from ALPACA. The consensus community or ALPACA membership was then overlaid on top by coloring the nodes. The angiogenesis genes (ellipse) were labeled based on whether they were ranked within the top 100 genes in the respective methods. Network is annotated with representative enriched GO terms in each module with P_adj_ < 0.05.

We then applied CRANE to 100 different ALPACA solutions, and found that CRANE showed good performance in recovering true disease-specific processes (Figure 6A). CRANE performed near equivalent to the “gold standard” subsample method and exhibited mild improvement over ALPACA at uncovering GO terms such as “angiogenesis” and “positive regulation of angiogenesis” (Figure 6A). Although the improvement in GO term detection in the individual modules was modest, we noticed that both CRANE and subsampling increased the ranking of “blood vessel development” related genes across the whole network (Supplementary Figure 2). This is because the blood vessel development genes are split across two modules which allows them to be masked by other enriched processes present in the same modules, such as inflammation pathways (Supplementary Figure 3). Interestingly, compared to ALPACA and consensus clustering, CRANE reduces signals from non-specific housekeeping processes, like “RNA transport” and “RNA processing.” The permutation and SBM method performed poorly in uncovering the disease-specific GO terms as these methods had a tendency to overestimate the DMS distribution while underestimating the variance (Figure 6C).

**Figure 6.**
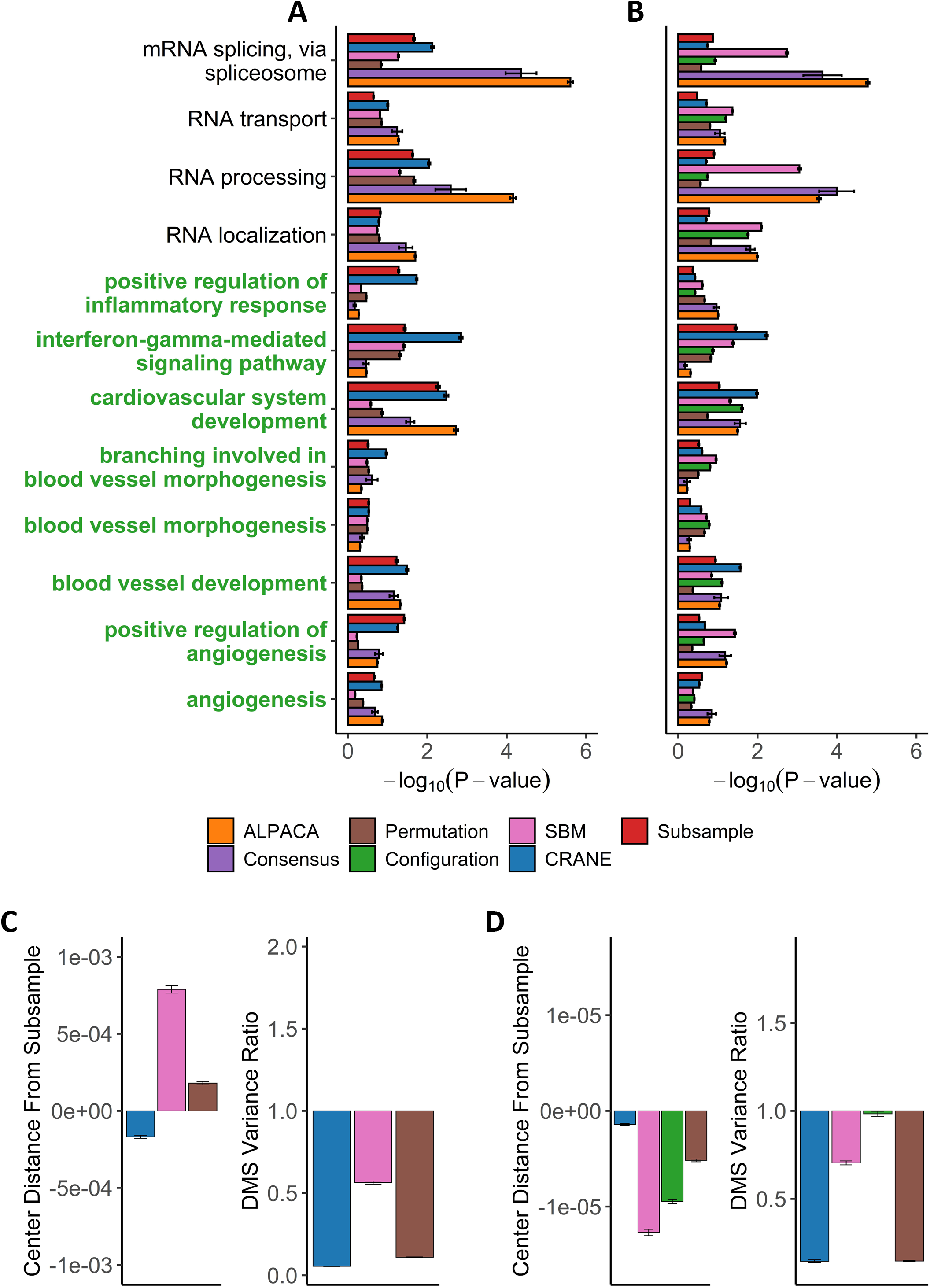
Performance of seven methods on discovering disease-relevant modules associated with angiogenic ovarian tumors. The top five-hundred genes in each module discovered by each method were extracted and subjected to GO term enrichment analysis. **(A, B)** Horizontal bar plots show a curated set of GO terms and their average −log_10_P-values over 100 different ALPACA runs. The GO terms (y-axis) colored in green are disease-relevant terms while black terms represent general biological processes. **(C, D)** The left vertical bar plot shows the average center distance and the right bar plot shows the average ratio between the mean of the null distribution created from subsampled networks and the mean of the null distribution generated from the indicated methods. Negative distance indicates that the specific method underestimates the center of the “gold standard” subsample distribution. For the variance ratio, values less than 1 represents greater variance in the subsample distribution compared to the indicated methods. The GO term enrichment analysis and the distribution analysis were performed on networks with either **(A, C)** PANDA edge weights or **(B, D)** exponentially transformed edge weights. The error bars represent mean+/-S.E.M.

CRANE and subsampling also consistently identified communities that represent inflammation and immune response. Genes in Module 1 deemed most significant by CRANE were enriched for interferon response, interleukins, cytokine signaling, and inflammation, consistent with the theory that chronic inflammation is associated with risk of cancer [48] (Supplementary table 2). Specifically, immunomodulators and interferon gamma been proposed as a therapeutic target in ovarian cancer [49, 50]. The enrichment in inflammation and immune response was not readily detectable using ALPACA, permutation, and SBM (Figure 6A, Supplementary table 3, 4, 5). Our results suggest that CRANE is able to uncover additional communities enriched with processes relevant to the disease phenotype.

We also tested our methods after exponentially transforming the edge weights and found that neither CRANE nor the “gold standard” subsampling method improve the recovery of angiogenesis related processes compared to ALPACA (Figure 6B). The exponentiation process leads to a change in community structure in the PANDA networks (NMI=0.69) that results in most of the blood vessel development genes being concentrated in a single giant differential module. (Supplementary Figure 4). The embedment of the angiogenesis genes in a large module along with overall increase in edge weight variance leads to reduction in CRANE performance, whereas other methods have inflated node p-values due to a tendency to underestimate the null distribution (Figure 6D).

#### Breast cancer

Breast cancer is the second most common cancer and a leading cause of death for women worldwide [51]. Although breast cancer is highly heterogeneous, one of its most important risk factors is overexpression of the estrogen receptor (ER+) leading to increased cell growth [52–54]. Cellular networks in ER+ breast tumors should therefore exhibit increased estrogen signaling.

We used PANDA to infer ER+ (1449 subjects) and ER- (445 subjects) gene regulatory networks from microarray data collected by the METABRIC consortium. After comparing the ER+ network to the ER-network using ALPACA, CRANE, and other methods, we analyzed the top-ranked genes from each method for enrichment in GO terms. Despite the expectations, consensus clustering and ALPACA both failed to detect estrogen specific pathways (Figure 7A, Supplementary Table 6, 7). Similar to the results from ovarian cancer, general biological processes such as RNA localization, mRNA splicing, protein catabolic process, and chromosome organization were highly enriched after consensus clustering (Supplemental Table 6).

**Figure 7.**
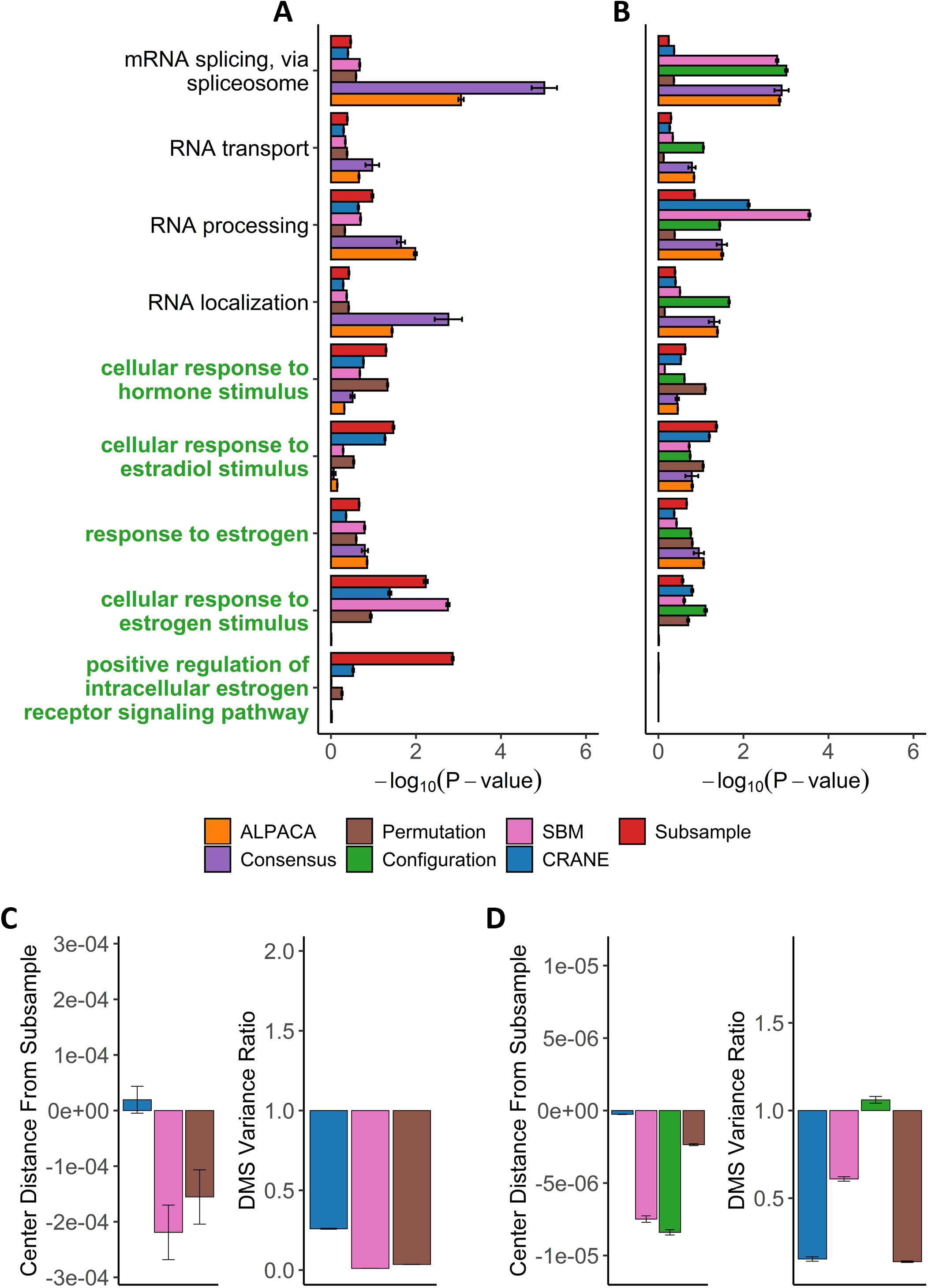
Performance of seven methods on discovering disease-relevant modules associated with ER-positive breast tumors. The top five-hundred genes in each module discovered by each method were extracted and subjected to GO term enrichment analysis. **(A, B)** Horizontal bar plots show a curated set of GO terms and their average −log_10_P-values over 100 different ALPACA runs. The GO terms (y-axis) colored in green are disease-relevant terms while black terms represent general biological processes. **(C, D)** The left vertical bar plot shows the average center distance and the right bar plot shows the average ratio between the mean of the null distribution created from subsampled networks and the mean of the null distribution generated from the indicated methods. Negative distance indicates that the specific method underestimates the center of the “gold standard” subsample distribution. For the variance ratio, values less than 1 represents greater variance in the subsample distribution compared to the indicated methods. The GO term enrichment analysis and the distribution analysis were performed on networks with either **(A, C)** PANDA edge weights or **(B, D)** exponentially transformed edge weights. The error bars represent mean+/-S.E.M.

On the contrary, reranking the nodes using CRANE effectively uncovered estrogen specific GO terms such as “cellular response to estrogen” and “positive regulation of intracellular estrogen receptor signaling pathway” with more significant p-values than ALPACA, consensus, and the permutation method (Figure 7A, Supplementary Table 8). Similar to what we observed in ovarian cancer, CRANE decreases the significance of non-specific housekeeping processes. The “gold standard” subsample distribution performed superior to CRANE, showing that real disease pathways are indeed robust relative to the empirical variance in regulatory networks.

We also applied the full panel of methods on exponentially transformed breast cancer PANDA networks. The exponential transformation decreased the discovery of estrogen related processes compared to the non-exponentiated network (Figure 7A, B). Nevertheless, all methods showed improvements in the significance level of “cellular response to estrogen stimulus” compared to ALPACA and consensus-ALPACA. Consistent with ovarian cancer networks, the configuration model had a tendency to underestimate the DMS distribution leading to a general inflation of GO term significance (Figure 7D). In comparison, the permutation methods perform well in discovering estrogen related GO terms. This is likely because for this specific condition, the permutation method produced a DMS distribution with small deviation from the subsample method. However, overall, the configuration model, SBM, and permutation methods show larger deviations in their DMS null distributions from the subsampled distribution, leading to unreliability in their performance, while CRANE exhibits near zero deviation from the subsampled distribution (Figure 6C, 6D, 7C, 7D). In summary, we found that different generative models may be useful in specific networks, context, and conditions, but only CRANE provides reliable and consistent performance across multiple settings in discovering disease relevant functional processes while minimizing noise from non-specific activities.

## Discussion

Phenotypic transitions like disease are often driven by the appearance of new groups of genes, or communities, that carry out relevant cellular processes. However, most community detection methods rely on maximizing a modularity-based score, and there is no easy way of determining whether the maxima represent true disease modules or whether they could have formed due to measurement noise. Consensus clustering offers an effective way of finding stable communities; however, the loss of community resolution leads to a reduction in interpretability. Additionally, biological experiments are resource-limited and do not generate enough data to empirically estimate network variance for statistical testing. To ensure that changes in network structure are truly associated with disease, it is important to devise a realistic null distribution for gene regulatory networks.

CRANE is a computational method for perturbing gene regulatory networks while maintaining fixed node strengths and the underlying “hard-wired” structure. The process of randomizing weighted bipartite networks with fixed node strength is challenging because each modification propagates to the rest of the network, leading to extreme edge weights if they are not properly controlled. For this reason, there is no accepted method for characterizing the full ensemble of strength-constrained weighted bipartite networks [28]. Existing methods rely on looser constraints, such as fixing the expectation value of the node strength over an ensemble of randomized networks. The well-established configuration model uses exponentially distributed edge weights to randomize networks [37], leading to higher variance in edge weights than is realistic, given that transcriptional regulation is dictated by physical binding sites. The stochastic block model, another leading generative model, overestimates the variance by treating every edge in a community the same regardless of prior information [39]. To model transcription, it is critical to understand that the TF regulation is specified by many factors such as the presence of activators or repressors, epigenetics, and physical binding affinity to gene promoters, which restricts their regulation to a limited number of genes [55–58]. The configuration model and SBM ignore this restriction and assume that each TF node can influence all genes in the network (configuration) or all genes in a community (SBM), which leads to improper estimation of edge weight variance. Alternatively, network variance could be empirically estimated by subsampling data and building multiple networks from the same sample condition, but the resource limitations in generating large disease-specific clinical datasets often make this approach impractical. For these reasons, we devised CRANE as a way to better control the edge variance while exactly fixing the node strengths, thus creating a more representative ensemble for biological networks.

We demonstrated that CRANE is able to control the variance in the community structure, edge weights, and differential modularity score (DMS) through the strength parameter *α*. However, the data shows that there is no single *α* value where CRANE is able to fully recapitulate the community structure, edge weight, and modularity variance of subsampled networks. This is likely because CRANE randomly perturbs all edges while subsampled networks retain the full correlation between the network edges. Creating an accurate perturbation method that perfectly simulates the subsampled network is challenging because each biological process is difficult to generalize. When applied to cancer networks, CRANE shows high accuracy in finding the center position of the subsample distribution but is less accurate in predicting the subsample variance (Figure 6C, 6D, 7C, 7D). It is tempting to hypothesize that better modeling of the variance of the null distribution would further improve the performance of CRANE. Although CRANE is unable to model all aspects of the subsampled distribution, we hypothesized that it could still help rank the most significant nodes and reduce noise in the network modules.

We evaluated the performance of CRANE on simulated networks and compared it with leading generative models and a naïve permutation method for randomizing network edges. CRANE was consistently more successful in recovering artificially added disease modules than all the other methods (Figure 4A). This is likely due to the strict constraints in CRANE that ensure the center of the null distribution of node score remains close to the true subsampled distribution while other conventional methods cause the distribution to deviate due to their looser constraints (Figure 3A, B). We note that the “gold standard” subsampled distribution performed best out of all the methods, again showing that there is room to further improve CRANE’s ability to capture all properties of gene regulatory networks.

CRANE also achieved more robust discovery of disease specific processes in cancer regulatory networks. Comparing angiogenic to non-angiogenic subtypes of ovarian cancer, we found that CRANE leads to a mild improvement in detecting GO terms like blood vessel development and cardiovascular development while also detecting disease-relevant inflammatory processes. Additionally, CRANE was able to minimize spurious signals from generic cellular processes that often dominate community structure in regulatory networks due to inherent biases from integrating prior information, like TF binding sites, during network inference. This indicates that CRANE is able to amplify the signal coming from disease-driving pathways while reducing noise from housekeeping processes that are present in all living cells.

In another test of the method, we compared ER+ to ER-subtypes of breast cancer and found that ALPACA and consensus clustering failed to identify modules related to estrogen signaling. In contrast, CRANE was able to discover that estrogen-related genes were highly significant in driving the modules specific to ER+ breast cancer. The superior performance of CRANE in breast cancer relative to ovarian cancer is likely rooted in differences in the performance of ALPACA in the two datasets. CRANE re-ranks the genes within ALPACA modules based on their significance. The angiogenesis genes were already highly clustered by ALPACA in the ovarian modules, making the improvements by CRANE modest (Supplemental Figure 3, 5). In breast cancer, the estrogen genes were less strongly clustered by ALPACA, providing more room for CRANE to improve the ranking of genes. We also found that CRANE performs poorly after exponentiating edge weights, because the exponential transformation leads to a reduction in ALPACA resolution (Supplemental Figure 4, 6). Therefore, exploring other edge weight transformation methods that retain the community structure while properly minimizing the noise from the negative PANDA edge weights may improve the performance of ALPACA and CRANE. Lastly, future improvements in the segregation of disease specific genes in ALPACA modules will in turn improve the performance of CRANE.

CRANE is a general computational tool that can be applied to both weighted unipartite and bipartite networks to generate a simulated network ensemble that retains the essential characteristics of gene regulatory networks. We have demonstrated that CRANE can be used to evaluate network modules and rank the nodes according to how significantly they drive the appearance of true disease modules. We expect that CRANE could be used in the future to evaluate the significance of other characteristics (e.g. information flow or betweenness centrality) in biological networks that are built around a “skeleton” comprised of prior information like TF binding sites. As gene regulatory networks become an increasingly common framing device for interpreting multi-omics data, CRANE provides a robust approach to identify what aspects of these networks are truly altered in disease.

## Author Contributions

MP and JTL conceived of the project. JTL developed the algorithm, performed analyses, and wrote the paper. CC performed comparison with network generative models. ADG and MP helped refine the analyses. All authors helped write the paper.

## Acknowledgements

This publication is supported by Institutional Research Grant number IRG-16-124-37 from the American Cancer Society.

