## Supplementary Method for "Generating Ensembles of Gene Regulatory Networks to Assess Robustness of Disease Modules"

### Constrained Random Alteration of Network Edges

Here we will provide a detailed implementation of CRANE (Constrained Random Alteration of Network Edges) and describe some of the challenges in randomizing weighted bipartite networks with fixed node strength.

#### 1 Edge Weight Perturbations Propagate

As an example, we will present a naive perturbation method to portray some of the challenges in altering network edge weights while fixing the node strength. We start with a bipartite network with $m$ TFs and $n$ genes. The edge weight from TF $i$ to gene $j$ is given by $a_{ij}$. We want to sample networks with matching strength sequence for both the TF nodes and gene nodes. The node degrees are given by

$$d_i = \sum_{j=1}^{n} a_{ij} \tag{1}$$

$$k_j = \sum_{i=1}^{m} a_{ij} \tag{2}$$

$$\sum_{i=1}^{m} d_i = \sum_{j=1}^{n} k_j \tag{3}$$

Fixing the strength sequences means we can only freely perturb $(n-1)$ edges for every TF and $(m-1)$ edges for every gene. The following steps will perturb the network while keeping the strength sequence the same.

**Step 1**: For each $\mathrm{TF}_i$ where $i \in [1, m-1]$, we can choose any perturbation for each edge to genes $\mathrm{G}_j$ for $j \in [1, n-1]$, and the final edge $b_{in}$ is determined by the degree of $\mathrm{TF}_i$:

$$b_{ij} = a_{ij} + \Delta_{ij} \quad \forall j \in [1, n-1] \tag{4}$$

$$b_{in} = d_i - \sum_{j=1}^{n-1} b_{ij} = a_{in} - \sum_{j=1}^{n-1} \Delta_{ij} \tag{5}$$

**Step 2**: The edge weights from $\text{TF}_m$ to all the $n$ genes, given by $b_{mj}$, are all completely determined by the constraints on the network at this point, and they can be computed as follows:

$$b_{mj} \;=\; k_j - \sum_{i=1}^{m-1} b_{ij} \tag{6}$$

However, due to the variability in the $\sum_{j=1}^{n-1} \Delta_{ij}$ the final edges could end up with either very large or very small values. This is undesirable since we do not want to have one TF or gene with edge weight that is significantly outside of its normal range.

#### 2 An Iterative Algorithm for Perturbing the network

We introduce Constrained Random Alteration of Network Edges (CRANE), a computational method that limits the propagation of edge weight perturbations. In this algorithm we will perturb and update the edge weights as we add one TF at a time while keeping the strength sequence the same.

**Optional Step:** If desired, we can introduce random perturbation to the degree sequence to both the TF nodes and gene nodes. The strength of perturbation will be determined by $\beta$. We then distribute the perturbation equally to all the edges.

$$\begin{aligned}
u &\sim \text{N}(0, \beta) && (7)\\
\tilde{d}_i &= d_i + u && (8)\\
\tilde{k}_j &= k_j + u && (9)\\
\tilde{a}_{ij} &= a_{ij} + \frac{\tilde{d}_i - d_i}{m} + \frac{\tilde{k}_j - k_j}{n} && (10)
\end{aligned}$$

**CRANE Algorithm:** To start, we will initialize some values. Given $a_{ij}$ is the adjacency matrix of the original network where rows and columns are randomly shuffled and $b_{ij}$ is an empty adjacency matrix that will become the perturbed network, we initialize the first row (first TF) of $b_{ij}$ with edges from the first row of $a_{ij}$. Additionally we define our perturbation parameters by sampling random numbers from a normal distribution centered at zero with standard deviation matching the original edge weights from the same TF. The parameter $\alpha$ determines the strength of the perturbation. Then for each $\text{TF}_l$, where $l = [1, \cdots, m-1]$, we will perturb each edge to genes $\text{G}_j$ for $j \in [1, n]$ and normalize the edge weights with $\frac{d_l}{\sum_{j=1}^{n} b_{lj}}$ to keep equation (1) true. Next, we determine the edges arising from $\text{TF}_{l+1}$ to gene $G_j$ (values in $b_{l+1,j}$) edges by subtracting the degree from the current perturbed state $\sum_{i=1}^{l} b_{ij}$ from the expected gene degree from the original network $\sum_{i=1}^{l+1} a_{ij}$. This allows us to keep equation (2) true. For some edges arising from $\text{TF}_{l+1}$ to gene $\text{G}_j$ (values in $b_{l+1,j}$), the individual edge values may exceed the maximum or the minimum edge weight from the original network. Therefore we include an inner loop to correct the extremely large or small values. For any $b_{l+1,j}$ bound outside of $a_{ij}$, we add $b_{l+1,j} - \min(a_{ij})$ or $b_{l+1,j} - \max(a_{ij})$ to $b_{lj}$. The adjusted $b_{lj}$ values are normalized and a new set of $b_{l+1,j}$ edge weights are computed. This inner loop will repeat until extreme values are corrected as describe above. The algorithm iteratively move down each row in $b_{ij}$ and repeat the procedure until the matrix is filled. The full algorithm is provided below in pseudocode format.

**Algorithm 1** CRANE

Given $a_{ij}$ is the adjacency matrix<br>
Randomize rows and columns of $a_{ij}$<br>
$d_i$ is the column sum of $a_{ij}$<br>
$k_j$ is the row sum of $a_{ij}$<br>
Set $b_{ij} = O_{m \times n}$<br>
Initialize $b_{1j} = a_{1j}$<br>
**for** $l = 1, \ldots, m-1$ **do**<br>
    $\sigma =$ standard deviation of row $i$<br>
    $u \sim N(0, \sigma)$<br>
    perturb $b_{lj}$ by adding $\alpha * u$, where $\alpha$ is the strength parameter<br>
    normalize $b_{lj}$ by multiplying $\frac{d_l}{\sum_{j=1}^{n} b_{lj}}$<br>
    compute the next row edges with $b_{l+1,j} = \sum_{i=1}^{l+1} a_{ij} - \sum_{i=1}^{l} b_{ij}$<br>
    **while** $b_{l+1,j} > \max(a_{ij})$ or $b_{l+1,j} < \min(a_{ij})$ **do**<br>
        **if** any value in $b_{l+1,j}$ is less than $\min(a_{ij})$ **then**<br>
            $b_{lj} = b_{lj} + b_{l+1,j} - \min(a_{ij})$<br>
        **else if** any value in $b_{l+1,j}$ is greater than $\max(a_{ij})$ **then**<br>
            $b_{lj} = b_{lj} + b_{l+1,j} - \max(a_{ij})$<br>
        **end if**<br>
        normalize $b_{lj}$ by multiplying $\frac{d_l}{\sum_{j=1}^{n} b_{lj}}$<br>
        recompute the next row edges with $b_{l+1,j} = \sum_{i=1}^{l+1} a_{ij} - \sum_{i=1}^{l} b_{ij}$<br>
    **end while**<br>
**end for**
