## Supplementary Figure for "Generating Ensembles of Gene Regulatory Networks to Assess Robustness of Disease Modules"

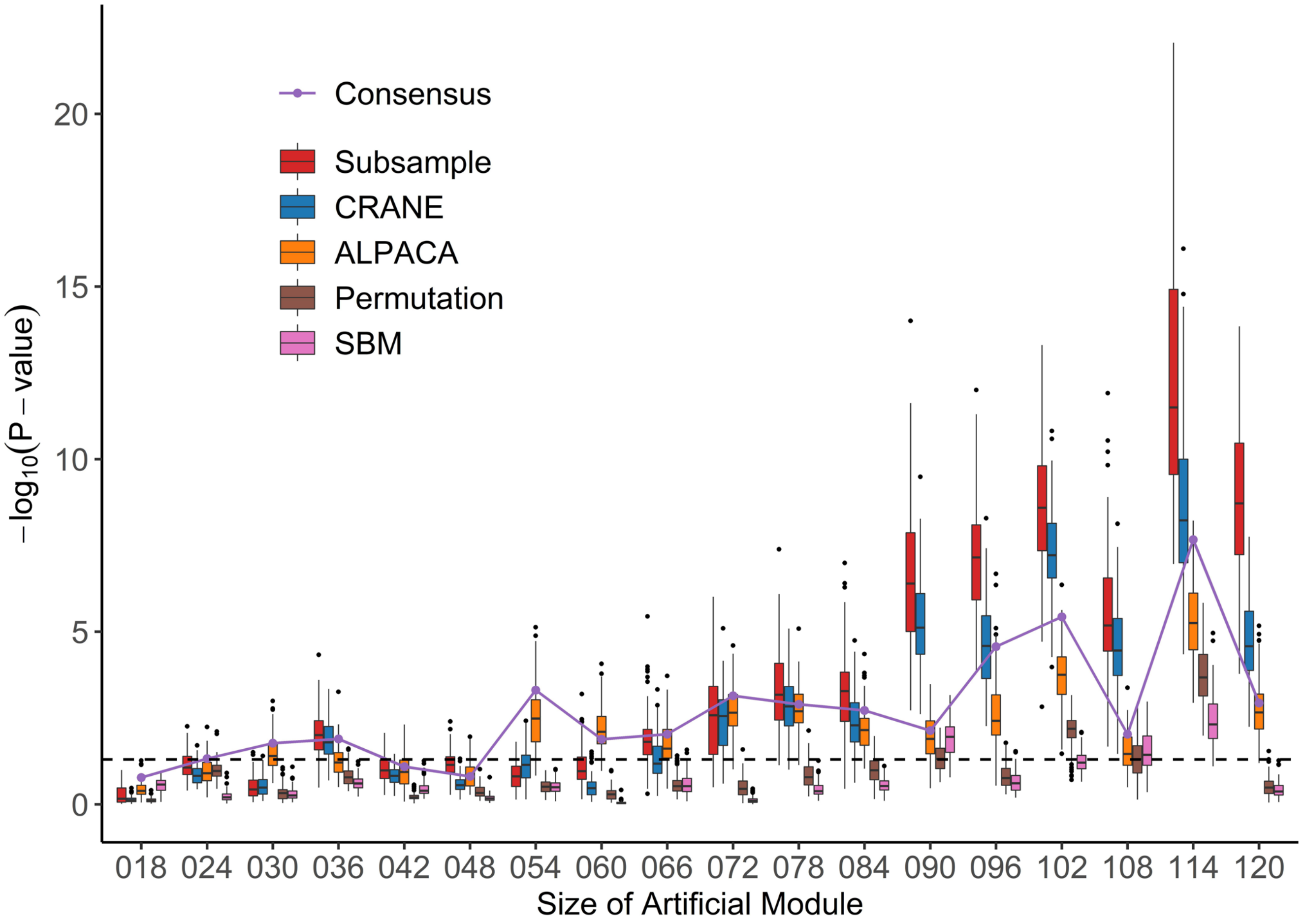


**Supplementary Figure 1. Performance of six methods on identifying artificial modules in simulated disease networks with non-exponentiated edge weights. (A)** Box plot shows performance of each method – subsample, CRANE, ALPACA,, permutation, or SBM – on network simulations without exponentially transformed edge weights. P-values (y-axis) for various sizes of artificial modules (x-axis) are computed using Wilcoxon test.

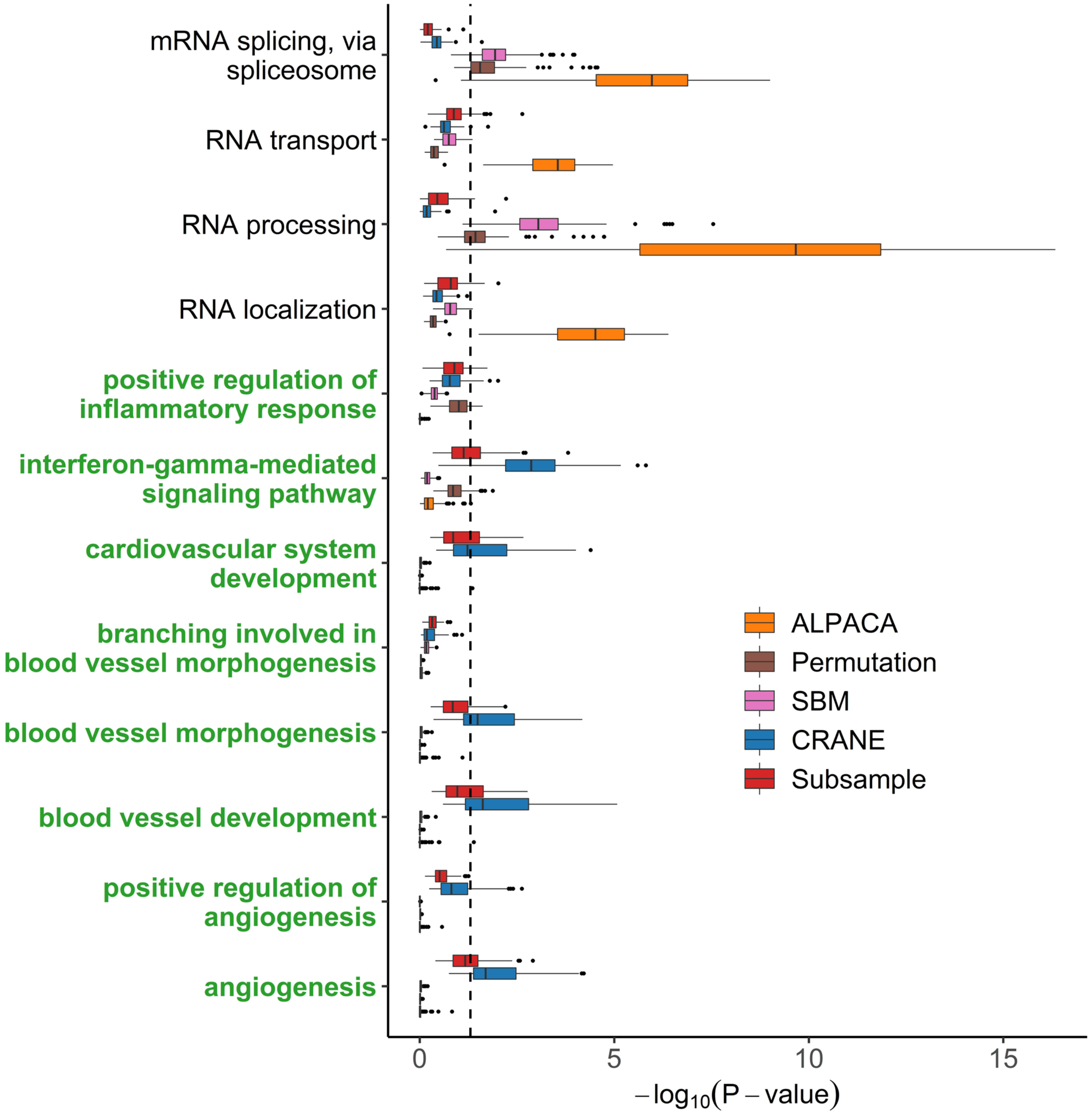


**Supplementary Figure 2. Core enrichment of disease relevant gene sets associated with angiogenic ovarian tumors.** Wilcoxon rank-sum test was used to evaluate how significantly each gene sets from the GO term process were highly ranked over the entire network. ALPACA ranked the genes by differential modularity scores while other methods ranked the genes by p-value. The GO terms (y-axis) colored in green are disease-relevant terms while black terms represent general biological processes.

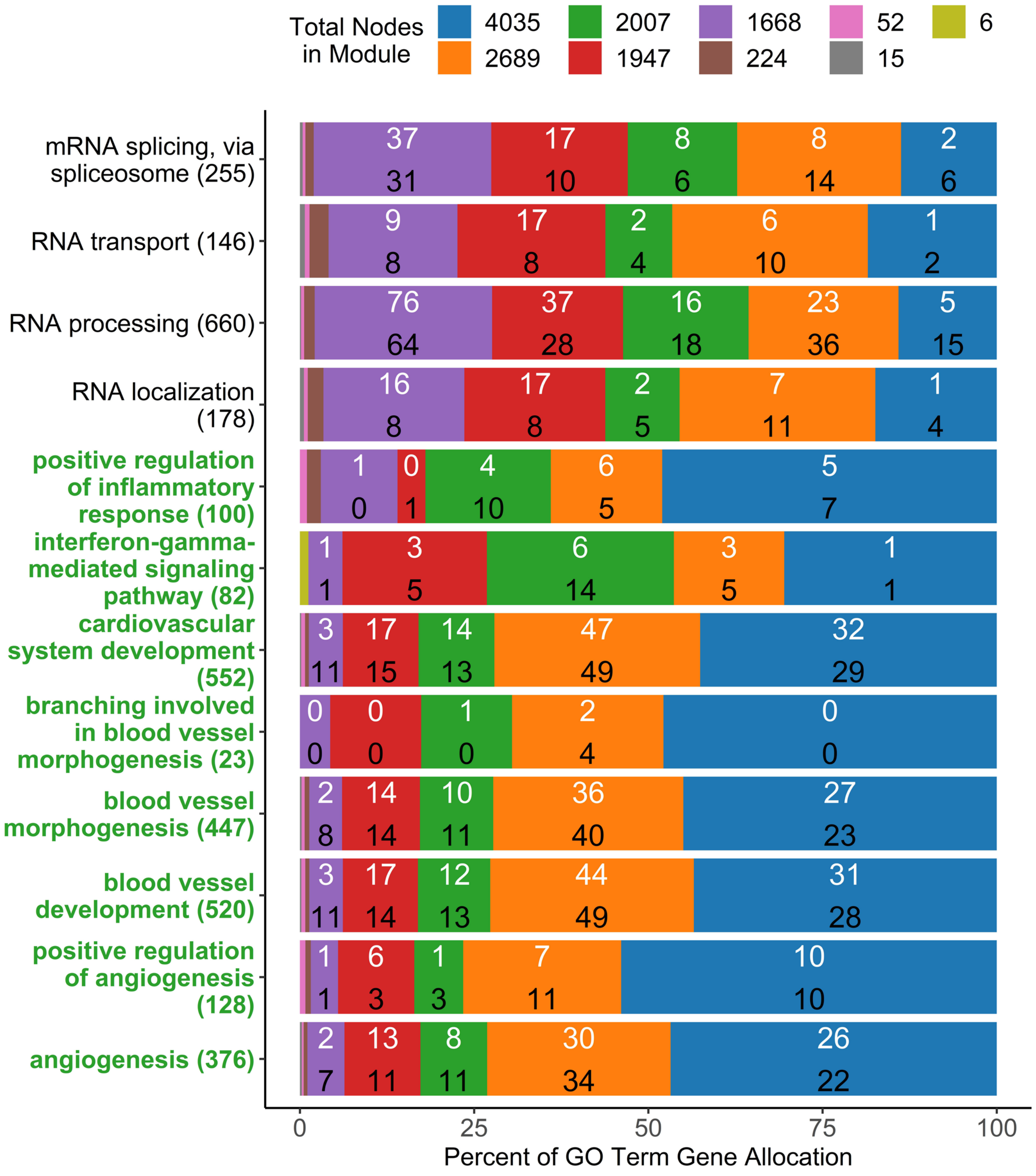


**Supplementary Figure 3. Module allocation of disease relevant gene sets associated with angiogenic ovarian tumors in non-exponentiated network.** The bar plot represents percent ratio of GO term genes allocated in different modules for a single run of ALPACA and CRANE with non-exponentiated edge weights. The white value inside the bars represents the number of GO term genes appeared in the top five-hundred ALPACA ranked genes while black values represents CRANE ranked genes within each module. The number in the parenthesis in the y-axis show the total number of gene for that GO term. The GO terms (y-axis) colored in green are disease-relevant terms while black terms represent general biological processes. The legends indicate the total number of genes found in each module by color.

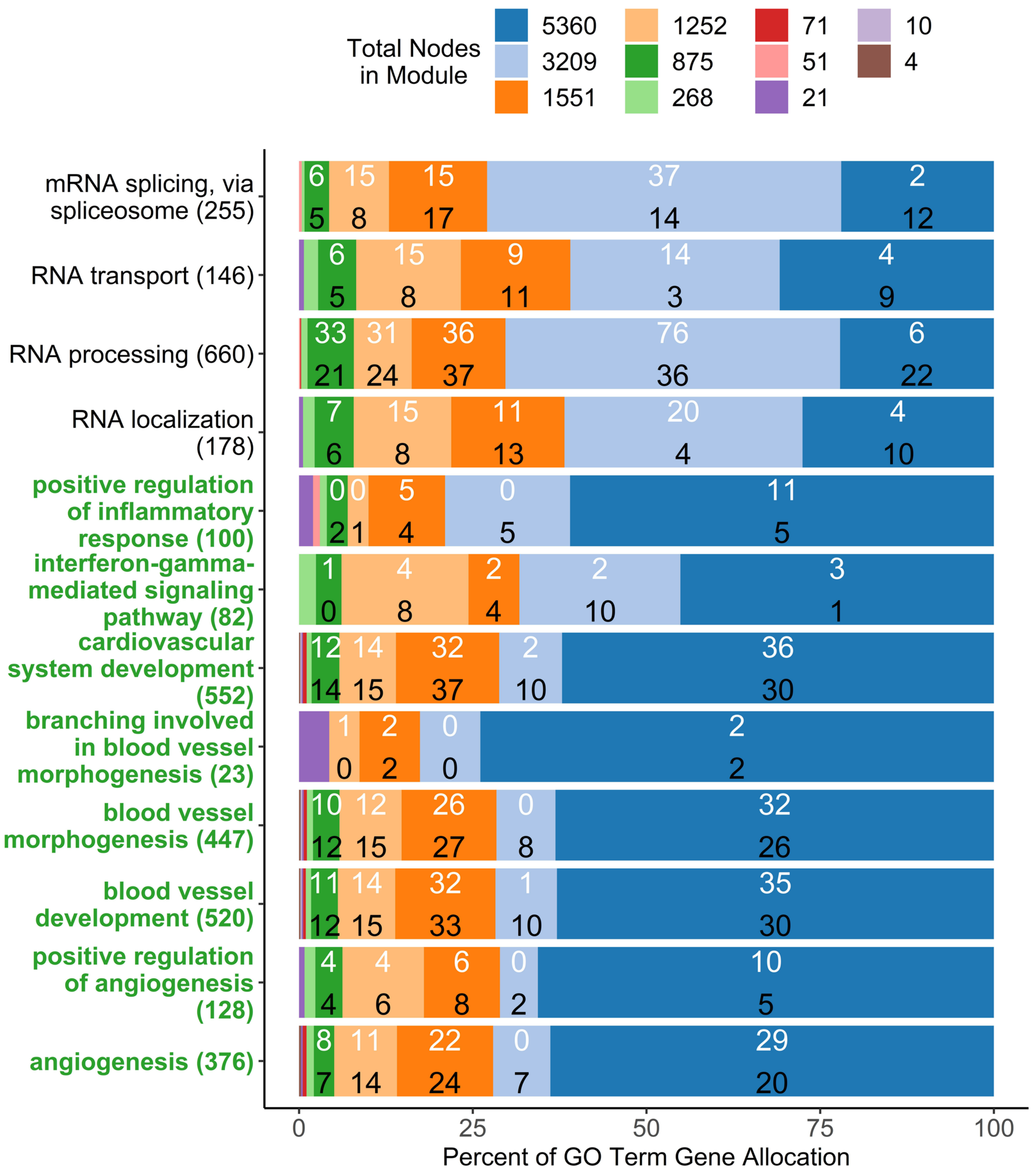


**Supplementary Figure 4. Module allocation of disease relevant gene sets associated with angiogenic ovarian tumors in exponentiated network.** The bar plot represents percent ratio of GO term genes allocated in different modules for a single run of ALPACA and CRANE with exponentiated edge weights. The white value inside the bars represents the number of GO term genes appeared in the top five-hundred ALPACA ranked genes while black values represents CRANE ranked genes within each module. The number in the parenthesis in the y-axis show the total number of gene for that GO term. The GO terms (y-axis) colored in green are disease-relevant terms while black terms represent general biological processes. The legends indicate the total number of genes found in each module by color.

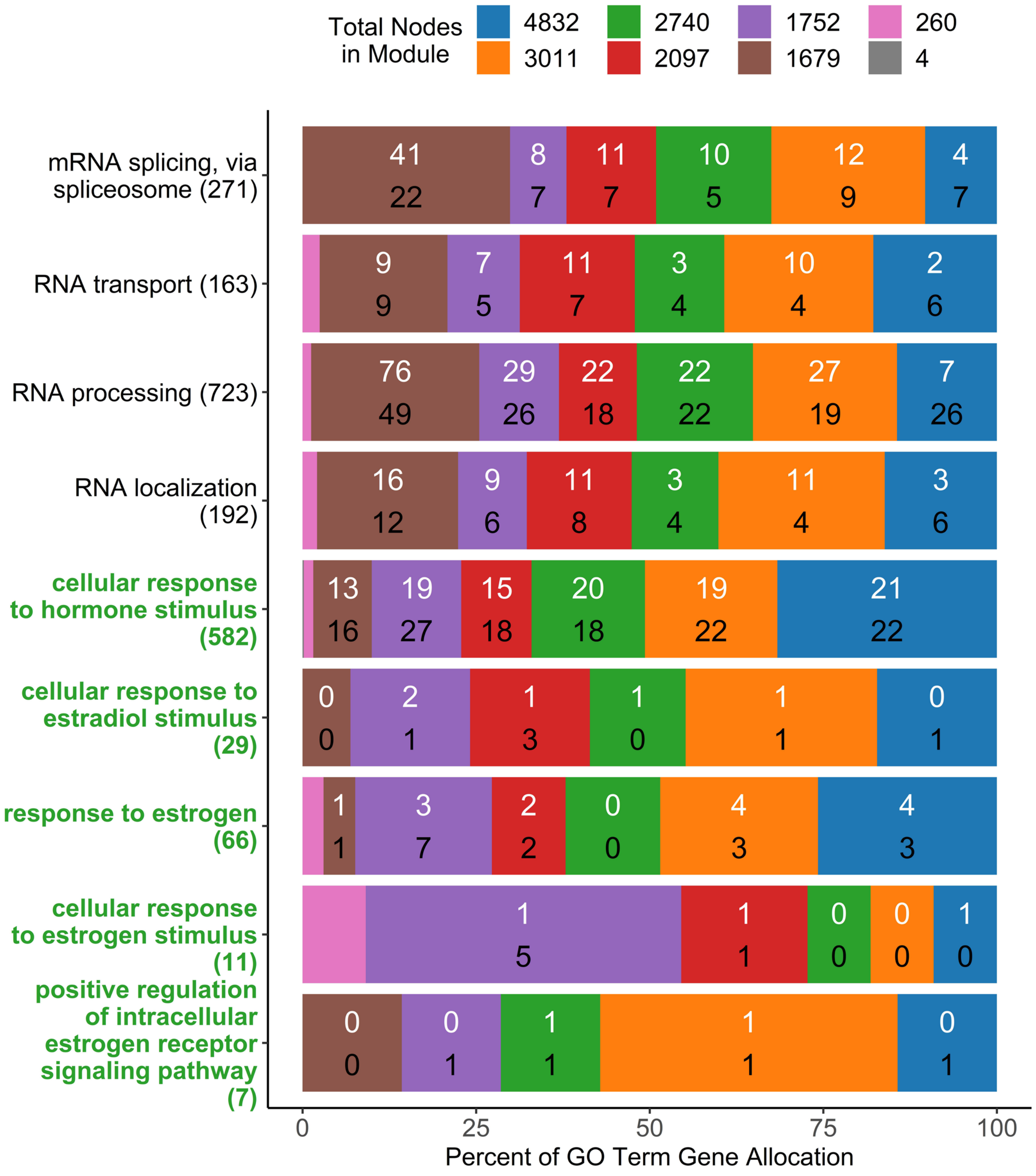


**Supplementary Figure 5. Module allocation of disease relevant gene sets associated with estrogen receptor positive breast tumors in non-exponentiated network.** The bar plot represents percent ratio of GO term genes allocated in different modules for a single run of ALPACA and CRANE with non-exponentiated edge weights. The white value inside the bars represents the number of GO term genes appeared in the top five-hundred ALPACA ranked genes while black values represents CRANE ranked genes within each module. The number in the parenthesis in the y-axis show the total number of gene for that GO term. The GO terms (y-axis) colored in green are disease-relevant terms while black terms represent general biological processes. The legends indicate the total number of genes found in each module by color.

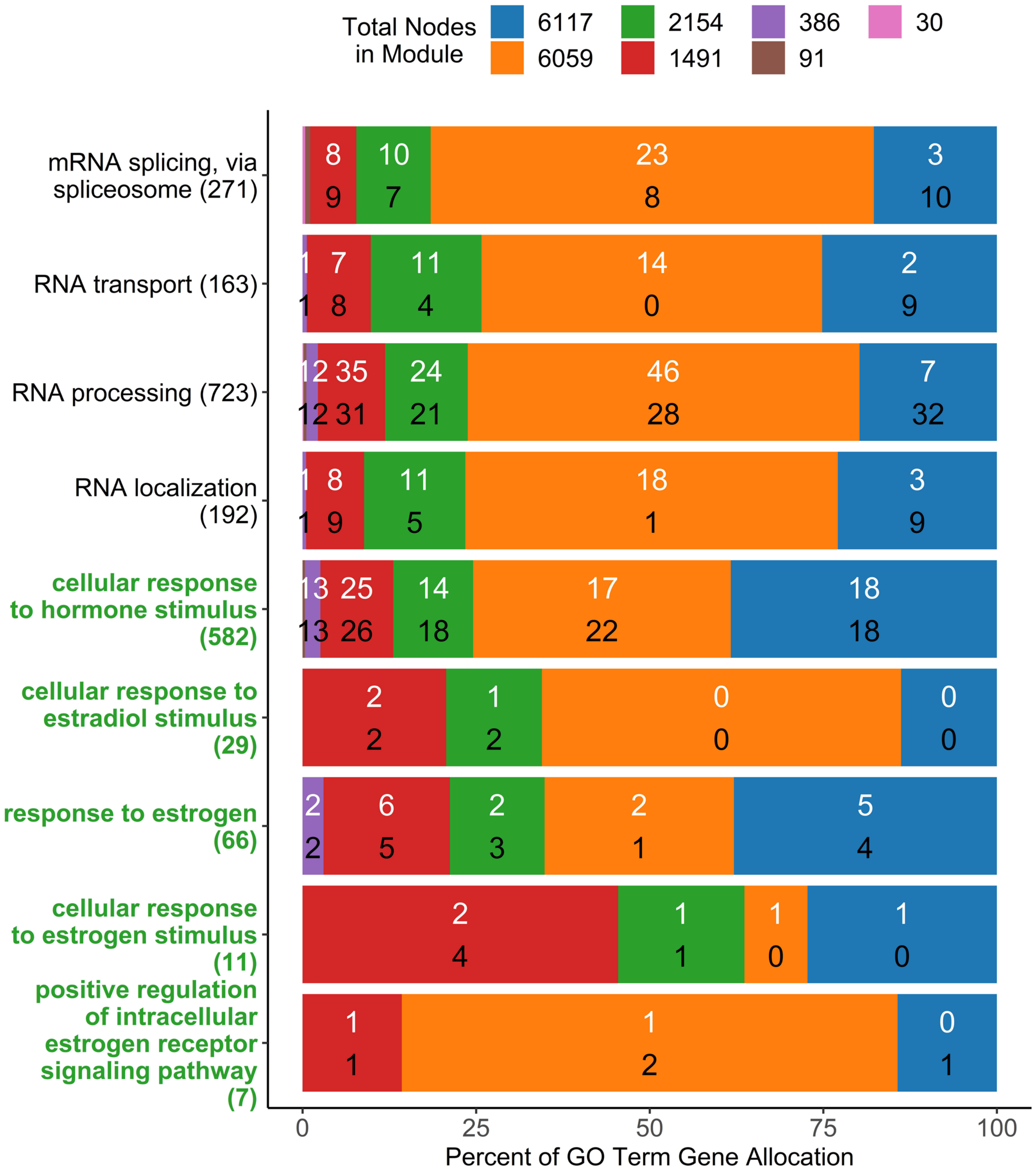


**Supplementary Figure 6. Module allocation of disease relevant gene sets associated with estrogen receptor positive breast tumors in exponentiated network.** The bar plot represents percent ratio of GO term genes allocated in different modules for a single run of ALPACA and CRANE with exponentiated edge weights. The white value inside the bars represents the number of GO term genes appeared in the top five-hundred ALPACA ranked genes while black values represents CRANE ranked genes within each module. The number in the parenthesis in the y-axis show the total number of gene for that GO term. The GO terms (y-axis) colored in green are disease-relevant terms while black terms represent general biological processes. The legends indicate the total number of genes found in each module by color.
