## Supplementary Table for "Generating Ensembles of Gene Regulatory Networks to Assess Robustness of Disease Modules"

**Supplementary Table 1: GO Terms Enriched in Non-Exponentiated Angiogenic Ovarian Cancer Networks Using Consensus Method**

| Modules | GOID | OddsRatio | pval | adjPval | GO Terms |
|---|---|---|---|---|---|
| Module 002 | GO:0000478 | 196.8828829 | 0.000122355 | 0.014437854 | endonucleolytic cleavage involved in rRNA processing |
| Module 002 | GO:0000027 | 73.7972973 | 0.000544439 | 0.021414583 | ribosomal large subunit assembly |
| Module 002 | GO:0000460 | 73.7972973 | 0.000544439 | 0.021414583 | maturation of 5.8S rRNA |
| Module 002 | GO:0042254 | 30.58522727 | 0.000205002 | 0.016126856 | ribosome biogenesis |
| Module 002 | GO:0006296 | 26.70588235 | 0.000296991 | 0.017522481 | nucleotide-excision repair, DNA incision, 5'-to lesion |
| Module 002 | GO:0090305 | 13.98761812 | 5.04E-07 | 0.000118959 | nucleic acid phosphodiester bond hydrolysis |
| Module 002 | GO:0035350 | Inf | 0.001140988 | 0.033190801 | FAD transmembrane transport |
| Module 002 | GO:0006777 | 79.65567766 | 0.000699445 | 0.012186485 | Mo-molybdopterin cofactor biosynthetic process |
| Module 002 | GO:0032324 | 59.73626374 | 0.001043375 | 0.014410393 | molybdopterin cofactor biosynthetic process |
| Module 002 | GO:0051189 | 59.73626374 | 0.001043375 | 0.014410393 | prosthetic group metabolic process |
| Module 002 | GO:0072396 | 40.66923077 | 0.000432525 | 0.016180869 | response to cell cycle checkpoint signaling |
| Module 002 | GO:0061635 | 38.06315789 | 0.002249079 | 0.040342847 | regulation of protein complex stability |
| Module 002 | GO:0006744 | 26.25730519 | 0.000471316 | 0.021224574 | ubiquinone biosynthetic process |
| Module 002 | GO:0042273 | 25.11 | 0.000369273 | 0.007159799 | ribosomal large subunit biogenesis |
| Module 002 | GO:1904874 | 20.50048309 | 4.03E-06 | 0.00033894 | positive regulation of telomerase RNA localization to Cajal body |
| Module 002 | GO:0006521 | 18.32826748 | 1.64E-05 | 0.00285458 | regulation of cellular amino acid metabolic process |
| Module 002 | GO:1904871 | 17.46721311 | 0.001623921 | 0.035608151 | positive regulation of protein localization to Cajal body |
| Module 002 | GO:0090670 | 17.07729469 | 5.08E-05 | 0.002973545 | RNA localization to Cajal body |
| Module 002 | GO:0090672 | 15.02083333 | 0.001628897 | 0.041446392 | telomerase RNA localization |
| Module 002 | GO:1903405 | 14.9704918 | 0.00227189 | 0.044916392 | protein localization to nuclear body |
| Module 002 | GO:1904851 | 14.9704918 | 0.00227189 | 0.044916392 | positive regulation of establishment of protein localization to telomere |
| Module 002 | GO:1902036 | 14.10356492 | 5.26E-05 | 0.0046553 | regulation of hematopoietic stem cell differentiation |
| Module 002 | GO:0051131 | 14.01447368 | 0.00048831 | 0.014722553 | chaperone-mediated protein complex assembly |
| Module 002 | GO:0031146 | 13.87480799 | 5.66E-05 | 0.0046553 | SCF-dependent proteasomal ubiquitin-dependent protein catabolic process |
| Module 002 | GO:0002479 | 12.83345179 | 8.00E-05 | 0.0046553 | antigen processing and presentation of exogenous peptide antigen via MHC class I, TAP-dependent |
| Module 002 | GO:0061418 | 12.83345179 | 8.00E-05 | 0.0046553 | regulation of transcription from RNA polymerase II promoter in response to hypoxia |
| Module 002 | GO:0006614 | 12.54320988 | 7.92E-07 | 0.000358946 | SRP-dependent cotranslational protein targeting to membrane |
| Module 002 | GO:0010972 | 12.28004535 | 9.73E-05 | 0.004814006 | negative regulation of G2/M transition of mitotic cell cycle |
| Module 002 | GO:0038061 | 12.10953347 | 0.000104919 | 0.003399135 | NIK/NF-kappaB signaling |
| Module 002 | GO:0031145 | 11.9367284 | 0.00011035 | 0.004814006 | anaphase-promoting complex-dependent catabolic process |
| Module 002 | GO:0072599 | 11.27947712 | 1.66E-06 | 0.000375784 | establishment of protein localization to endoplasmic reticulum |
| Module 002 | GO:0000387 | 10.35839501 | 0.000296892 | 0.010850049 | spliceosomal snRNP assembly |
| Module 002 | GO:0000184 | 9.941407151 | 3.98E-06 | 0.000450254 | nuclear-transcribed mRNA catabolic process, nonsense-mediated decay |
| Module 002 | GO:0002474 | 9.864988141 | 0.000255677 | 0.0064948 | antigen processing and presentation of peptide antigen via MHC class I |
| Module 002 | GO:0044106 | 9.864988141 | 0.000255677 | 0.0064948 | cellular amine metabolic process |
| Module 002 | GO:0002223 | 9.641519529 | 0.000282673 | 0.0064948 | stimulatory C-type lectin receptor signaling pathway |
| Module 002 | GO:0070498 | 9.5335097 | 0.000296957 | 0.0064948 | interleukin-1-mediated signaling pathway |
| Module 002 | GO:0006402 | 9.504117436 | 0.000303725 | 0.0064948 | mRNA catabolic process |
| Module 002 | GO:0006369 | 8.982315891 | 4.38E-05 | 0.002678864 | termination of RNA polymerase II transcription |

| | | | | | |
|---|---|---|---|---|---|
| Module 002 | GO:0060071 | 8.932705026 | 0.000394508 | 0.007246492 | Wnt signaling pathway, planar cell polarity pathway |
| Module 002 | GO:0038095 | 8.572222222 | 0.000471846 | 0.007485194 | Fc-epsilon receptor signaling pathway |
| Module 002 | GO:0043620 | 8.572222222 | 0.000471846 | 0.007485194 | regulation of DNA-templated transcription in response to stress |
| Module 002 | GO:0070125 | 8.561708861 | 0.001718227 | 0.018739413 | mitochondrial translational elongation |
| Module 002 | GO:0051290 | 8.376284585 | 0.000213306 | 0.010625816 | protein heterotetramerization |
| Module 002 | GO:0070126 | 8.348765432 | 0.001875774 | 0.019254271 | mitochondrial translational termination |
| Module 002 | GO:0090090 | 8.081020255 | 0.000182028 | 0.0064948 | negative regulation of canonical Wnt signaling pathway |
| Module 002 | GO:0016070 | 8.07311828 | 6.45E-08 | 8.83E-06 | RNA metabolic process |
| Module 002 | GO:0001736 | 7.931363904 | 0.000660237 | 0.010018382 | establishment of planar polarity |
| Module 002 | GO:0006612 | 7.902941176 | 1.91E-05 | 0.001080554 | protein targeting to membrane |
| Module 002 | GO:0000245 | 7.65620501 | 0.000963104 | 0.026158924 | spliceosomal complex assembly |
| Module 002 | GO:0090263 | 7.64526644 | 0.000773235 | 0.011244127 | positive regulation of canonical Wnt signaling pathway |
| Module 002 | GO:0034080 | 7.606413575 | 0.001076318 | 0.032312448 | CENP-A containing nucleosome assembly |
| Module 002 | GO:0010467 | 7.49306184 | 0.00017268 | 0.023139116 | gene expression |
| Module 002 | GO:0019083 | 7.42198384 | 3.21E-06 | 0.000244294 | viral transcription |
| Module 002 | GO:0006413 | 7.415678137 | 6.44E-12 | 2.39E-09 | translational initiation |
| Module 002 | GO:1903706 | 7.324256738 | 0.000933604 | 0.013033112 | regulation of hemopoiesis |
| Module 002 | GO:0000375 | 6.660287081 | 0.000543649 | 0.048565971 | RNA splicing, via transesterification reactions |
| Module 002 | GO:0033209 | 6.575702076 | 0.001468652 | 0.018223399 | tumor necrosis factor-mediated signaling pathway |
| Module 002 | GO:0042180 | 6.461242176 | 0.001584049 | 0.018223399 | cellular ketone metabolic process |
| Module 002 | GO:0002244 | 6.425587779 | 0.001618697 | 0.018223399 | hematopoietic progenitor cell differentiation |
| Module 002 | GO:0043488 | 6.282096172 | 0.001779741 | 0.018822104 | regulation of mRNA stability |
| Module 002 | GO:0071453 | 5.731598646 | 0.001032117 | 0.013854181 | cellular response to oxygen levels |
| Module 002 | GO:1905330 | 5.727069351 | 0.002616372 | 0.024139573 | regulation of morphogenesis of an epithelium |
| Module 002 | GO:0044839 | 5.676608986 | 0.00044188 | 0.007485194 | cell cycle G2/M phase transition |
| Module 002 | GO:0050852 | 5.650162935 | 0.002766713 | 0.024139573 | T cell receptor signaling pathway |
| Module 002 | GO:0070555 | 5.650162935 | 0.002766713 | 0.024139573 | response to interleukin-1 |
| Module 002 | GO:0002478 | 5.525600836 | 0.000214033 | 0.009212745 | antigen processing and presentation of exogenous peptide antigen |
| Module 002 | GO:0044786 | 5.352642276 | 0.000309191 | 0.013813123 | cell cycle DNA replication |
| Module 002 | GO:0000398 | 5.348393574 | 0.003485255 | 0.02764441 | mRNA splicing, via spliceosome |
| Module 002 | GO:1901988 | 5.313151365 | 0.001496143 | 0.018223399 | negative regulation of cell cycle phase transition |
| Module 002 | GO:1903311 | 4.983529412 | 0.001974075 | 0.045353889 | regulation of mRNA metabolic process |
| Module 002 | GO:0044265 | 4.863896154 | 0.000105051 | 0.003399135 | cellular macromolecule catabolic process |
| Module 002 | GO:0048863 | 4.72707231 | 0.00571398 | 0.041545394 | stem cell differentiation |
| Module 002 | GO:0000209 | 4.563315031 | 0.006588521 | 0.045244776 | protein polyubiquitination |
| Module 002 | GO:0034655 | 4.490644491 | 3.73E-06 | 0.000450254 | nucleobase-containing compound catabolic process |
| Module 002 | GO:0042773 | 4.408939014 | 0.000456301 | 0.018707892 | ATP synthesis coupled electron transport |
| Module 002 | GO:0140053 | 4.377573078 | 2.12E-06 | 0.000157119 | mitochondrial gene expression |
| Module 002 | GO:0045930 | 4.139914497 | 0.004886338 | 0.036283658 | negative regulation of mitotic cell cycle |
| Module 002 | GO:0019882 | 4.100199426 | 0.001308005 | 0.029430112 | antigen processing and presentation |
| Module 002 | GO:0006283 | 3.974546371 | 0.000925204 | 0.029345063 | transcription-coupled nucleotide-excision repair |

| | | | | | |
|---|---|---|---|---|---|
| Module 002 | GO:0030111 | 3.884114425 | 0.006541275 | 0.045244776 | regulation of Wnt signaling pathway |
| Module 002 | GO:0044772 | 3.849741479 | 0.003553099 | 0.034990297 | mitotic cell cycle phase transition |
| Module 002 | GO:0034641 | 3.674140127 | 2.89E-06 | 0.000497195 | cellular nitrogen compound metabolic process |
| Module 002 | GO:0006457 | 3.642839117 | 0.000491186 | 0.018644097 | protein folding |
| Module 002 | GO:0044271 | 3.591796875 | 0.000460222 | 0.008686689 | cellular nitrogen compound biosynthetic process |
| Module 002 | GO:0046483 | 3.50273569 | 3.01E-06 | 0.000406401 | heterocycle metabolic process |
| Module 002 | GO:0022613 | 3.487256875 | 0.001151125 | 0.028922019 | ribonucleoprotein complex biogenesis |
| Module 002 | GO:0010468 | 3.478060141 | 0.000862886 | 0.013960264 | regulation of gene expression |
| Module 002 | GO:0006725 | 3.457291667 | 3.70E-06 | 0.000406401 | cellular aromatic compound metabolic process |
| Module 002 | GO:0090150 | 3.378430376 | 3.80E-07 | 3.14E-05 | establishment of protein localization to membrane |
| Module 002 | GO:0016071 | 3.34521862 | 4.45E-07 | 3.61E-05 | mRNA metabolic process |
| Module 002 | GO:0043043 | 3.302892157 | 0.000443294 | 0.008686689 | peptide biosynthetic process |
| Module 002 | GO:0032200 | 3.273824963 | 0.000266389 | 0.012537484 | telomere organization |
| Module 002 | GO:1901360 | 3.234335242 | 1.03E-05 | 0.000736219 | organic cyclic compound metabolic process |
| Module 002 | GO:0043624 | 3.1256917 | 0.000947466 | 0.028024733 | cellular protein complex disassembly |
| Module 002 | GO:0009059 | 3.104964379 | 1.81E-11 | 5.14E-09 | macromolecule biosynthetic process |
| Module 002 | GO:0006396 | 3.091736527 | 1.38E-06 | 0.000131876 | RNA processing |
| Module 002 | GO:0006139 | 3.086556169 | 1.10E-05 | 0.000918717 | nucleobase-containing compound metabolic process |
| Module 002 | GO:0043933 | 3.047652316 | 1.20E-05 | 0.000799321 | protein-containing complex subunit organization |
| Module 002 | GO:0071826 | 3.024726855 | 0.001250055 | 0.030046619 | ribonucleoprotein complex subunit organization |
| Module 002 | GO:0034645 | 3.017942976 | 0.000432486 | 0.016193607 | cellular macromolecule biosynthetic process |
| Module 002 | GO:0006605 | 3.000876168 | 9.64E-06 | 0.000651765 | protein targeting |
| Module 002 | GO:0010608 | 2.914615385 | 4.84E-05 | 0.002461477 | posttranscriptional regulation of gene expression |
| Module 002 | GO:0043603 | 2.878484454 | 5.41E-08 | 6.07E-06 | cellular amide metabolic process |
| Module 002 | GO:0071704 | 2.851628198 | 1.61E-06 | 0.00017276 | organic substance metabolic process |
| Module 002 | GO:0072657 | 2.81754822 | 0.005149063 | 0.048594283 | protein localization to membrane |
| Module 002 | GO:0016579 | 2.782943038 | 0.000375681 | 0.012704846 | protein deubiquitination |
| Module 002 | GO:0006518 | 2.737316017 | 7.97E-10 | 1.01E-07 | peptide metabolic process |
| Module 002 | GO:0043604 | 2.734951323 | 6.04E-10 | 8.38E-08 | amide biosynthetic process |
| Module 002 | GO:0034660 | 2.590450887 | 4.33E-05 | 0.002387204 | ncRNA metabolic process |
| Module 002 | GO:0044260 | 2.548665463 | 0.001982819 | 0.049032762 | cellular macromolecule metabolic process |
| Module 002 | GO:0044419 | 2.534600341 | 0.002276011 | 0.024548404 | interspecies interaction between organisms |
| Module 002 | GO:0006807 | 2.27032935 | 0.002747901 | 0.024139573 | nitrogen compound metabolic process |
| Module 002 | GO:0043170 | 2.188412853 | 0.002052883 | 0.047208859 | macromolecule metabolic process |
| Module 002 | GO:0070647 | 2.155270823 | 0.001825863 | 0.044798865 | protein modification by small protein conjugation or removal |
| Module 002 | GO:0006886 | 2.123044397 | 0.001450488 | 0.031217026 | intracellular protein transport |
| Module 002 | GO:0044237 | 2.010223147 | 0.000380491 | 0.012549994 | cellular metabolic process |
| Module 002 | GO:0090304 | 1.910476295 | 8.70E-10 | 1.81E-07 | nucleic acid metabolic process |
| Module 002 | GO:0044238 | 1.907150125 | 2.92E-06 | 0.000205914 | primary metabolic process |
| Module 002 | GO:0044403 | 1.791175988 | 0.002469773 | 0.046539783 | symbiont process |
| Module 002 | GO:1901576 | 1.773579263 | 0.000704162 | 0.02063857 | organic substance biosynthetic process |

| | | | | | |
|---|---|---|---|---|---|
| Module 002 | GO:1901566 | 1.666947835 | 0.000116255 | 0.005068696 | organonitrogen compound biosynthetic process |
| Module 002 | GO:0009058 | 1.531416666 | 5.59E-05 | 0.003132616 | biosynthetic process |
| Module 002 | GO:0000003 | 3.702411874 | 8.72E-06 | 0.010179427 | reproduction |
| Module 003 | GO:0035520 | 64.73809524 | 3.91E-05 | 0.019987244 | monoubiquitinated protein deubiquitination |
| Module 003 | GO:0016339 | 21.73634085 | 7.06E-05 | 0.024336058 | calcium-dependent cell-cell adhesion via plasma membrane cell adhesion molecules |
| Module 004 | GO:0030042 | 339.625 | 0.000107222 | 0.026912636 | actin filament depolymerization |
| Module 004 | GO:2001046 | 111.7846154 | 0.000363219 | 0.045583953 | positive regulation of integrin-mediated signaling pathway |
| Module 004 | GO:0021814 | 73.59589041 | 7.94E-05 | 0.047148733 | cell motility involved in cerebral cortex radial glia guided migration |
| Module 004 | GO:0016048 | 63.46511628 | 0.000725267 | 0.04516108 | detection of temperature stimulus |
| Module 004 | GO:0050965 | 63.46511628 | 0.000725267 | 0.04516108 | detection of temperature stimulus involved in sensory perception of pain |
| Module 004 | GO:2001224 | 63.46511628 | 0.000725267 | 0.04516108 | positive regulation of neuron migration |
| Module 004 | GO:0050951 | 56.40826873 | 0.000884125 | 0.045721911 | sensory perception of temperature stimulus |
| Module 004 | GO:0071474 | 50.7627907 | 0.001058184 | 0.047882813 | cellular hyperosmotic response |
| Module 004 | GO:0032418 | 29.12566845 | 0.000240265 | 0.04781275 | lysosome localization |
| Module 004 | GO:0032012 | 15.5539316 | 6.46E-05 | 0.01970451 | regulation of ARF protein signal transduction |
| Module 004 | GO:0007015 | 15.51621129 | 0.000209684 | 0.035087094 | actin filament organization |
| Module 004 | GO:2000463 | 15.23566265 | 1.78E-05 | 0.008361731 | positive regulation of excitatory postsynaptic potential |
| Module 004 | GO:0035023 | 14.24868421 | 5.21E-05 | 0.009431286 | regulation of Rho protein signal transduction |
| Module 004 | GO:1901223 | 12.75138654 | 0.000158638 | 0.034404705 | negative regulation of NIK/NF-kappaB signaling |
| Module 004 | GO:0030036 | 10.98610747 | 0.000748526 | 0.04516108 | actin cytoskeleton organization |
| Module 004 | GO:0007186 | 9.214912281 | 3.02E-05 | 0.009431286 | G protein-coupled receptor signaling pathway |
| Module 004 | GO:0050906 | 6.743761468 | 0.000233752 | 0.045698464 | detection of stimulus involved in sensory perception |
| Module 004 | GO:0031644 | 6.013199426 | 0.000160823 | 0.038990045 | regulation of neurological system process |
| Module 004 | GO:0007265 | 4.74713788 | 5.90E-05 | 0.01970451 | Ras protein signal transduction |
| Module 004 | GO:0051056 | 3.952358849 | 2.06E-08 | 3.14E-05 | regulation of small GTPase mediated signal transduction |
| Module 004 | GO:0008360 | 3.836996337 | 0.000107428 | 0.046056877 | regulation of cell shape |
| Module 004 | GO:0043065 | 3.778431632 | 0.000397679 | 0.048119197 | positive regulation of apoptotic process |
| Module 004 | GO:0043269 | 3.621121408 | 0.000185854 | 0.045418167 | regulation of ion transport |
| Module 004 | GO:0009966 | 3.485152529 | 9.41E-05 | 0.018982958 | regulation of signal transduction |
| Module 004 | GO:0007155 | 3.28031203 | 5.99E-05 | 0.032972666 | cell adhesion |
| Module 004 | GO:0050896 | 2.042350424 | 2.23E-06 | 0.001930551 | response to stimulus |
| Module 004 | GO:0023052 | 1.982236325 | 1.11E-05 | 0.015053728 | signaling |
| Module 004 | GO:0048870 | 1.935434517 | 0.000104798 | 0.031615801 | cell motility |
| Module 004 | GO:0007154 | 1.914721881 | 3.35E-05 | 0.015150157 | cell communication |
| Module 004 | GO:0051674 | 1.909721548 | 3.42E-06 | 0.003341634 | localization of cell |
| Module 004 | GO:0035556 | 1.888988391 | 9.38E-05 | 0.023519375 | intracellular signal transduction |
| Module 004 | GO:0040011 | 1.875169532 | 2.25E-06 | 0.003341634 | locomotion |
| Module 004 | GO:0006928 | 1.771564964 | 8.33E-06 | 0.00542679 | movement of cell or subcellular component |

**Supplementary Table 2: GO Terms Enriched in Non-Exponentiated Angiogenic Ovarian Cancer Networks Using CRANE**

| Modules | GOID | OddsRatio | pval | adjPval | GO Terms |
|---|---|---|---|---|---|
| Module 001 | GO:1903217 | Inf | 1.05E-05 | 0.002555441 | negative regulation of protein processing involved in protein targeting to mitochondrion |
| Module 001 | GO:0042495 | Inf | 5.52E-05 | 0.015458325 | detection of triacyl bacterial lipopeptide |
| Module 001 | GO:2000473 | Inf | 0.000128836 | 0.007898638 | positive regulation of hematopoietic stem cell migration |
| Module 001 | GO:0061033 | Inf | 0.000460984 | 0.029602195 | secretion by lung epithelial cell involved in lung growth |
| Module 001 | GO:2000502 | Inf | 0.000551135 | 0.032867714 | negative regulation of natural killer cell chemotaxis |
| Module 001 | GO:0002728 | Inf | 0.000859145 | 0.047498441 | negative regulation of natural killer cell cytokine production |
| Module 001 | GO:0038123 | 272.15 | 0.000164819 | 0.017452351 | toll-like receptor TLR1:TLR2 signaling pathway |
| Module 001 | GO:0042494 | 272.15 | 0.000164819 | 0.017452351 | detection of bacterial lipoprotein |
| Module 001 | GO:0071727 | 272.15 | 0.000164819 | 0.017452351 | cellular response to triacyl bacterial lipopeptide |
| Module 001 | GO:0034982 | 128.564706 | 0.000217673 | 0.023556679 | mitochondrial protein processing |
| Module 001 | GO:1903748 | 107.127451 | 0.000289631 | 0.023556679 | negative regulation of establishment of protein localization to mitochondrion |
| Module 001 | GO:0070339 | 90.7 | 0.000544072 | 0.027698211 | response to bacterial lipopeptide |
| Module 001 | GO:0071220 | 90.7 | 0.000544072 | 0.027698211 | cellular response to bacterial lipoprotein |
| Module 001 | GO:0045356 | 88.1463415 | 0.000761505 | 0.025288297 | positive regulation of interferon-alpha biosynthetic process |
| Module 001 | GO:0018344 | 68.01875 | 0.000812146 | 0.034984737 | protein geranylgeranylation |
| Module 001 | GO:0002480 | 63.2380952 | 6.83E-08 | 0.000109909 | antigen processing and presentation of exogenous peptide antigen via MHC class I, TAP-independent |
| Module 001 | GO:0043615 | 54.41 | 0.001131487 | 0.042242168 | astrocyte cell migration |
| Module 001 | GO:0097354 | 45.3375 | 0.001501332 | 0.046708111 | prenylation |
| Module 001 | GO:0098543 | 45.3375 | 0.001501332 | 0.046708111 | detection of other organism |
| Module 001 | GO:0045359 | 38.214455 | 0.000241879 | 0.0249332 | positive regulation of interferon-beta biosynthetic process |
| Module 001 | GO:0046641 | 36.1133333 | 1.43E-05 | 0.003791646 | positive regulation of alpha-beta T cell proliferation |
| Module 001 | GO:0046634 | 33.678125 | 0.00021611 | 0.011482625 | regulation of alpha-beta T cell activation |
| Module 001 | GO:0045351 | 27.9155673 | 1.12E-05 | 0.002489987 | type I interferon biosynthetic process |
| Module 001 | GO:0002407 | 24.2814594 | 0.000418799 | 0.026385184 | dendritic cell chemotaxis |
| Module 001 | GO:0032729 | 23.4371968 | 4.72E-05 | 0.008076784 | positive regulation of interferon-gamma production |
| Module 001 | GO:0035520 | 23.0193133 | 0.000750483 | 0.042647055 | monoubiquitinated protein deubiquitination |
| Module 001 | GO:0045954 | 22.6846154 | 0.000599097 | 0.02231181 | positive regulation of natural killer cell mediated cytotoxicity |
| Module 001 | GO:0001774 | 18.0266667 | 0.000996701 | 0.030552721 | microglial cell activation |
| Module 001 | GO:0007252 | 17.752459 | 0.001040279 | 0.030707497 | I-kappaB phosphorylation |
| Module 001 | GO:0150076 | 16.8765625 | 0.001178241 | 0.032561965 | neuroinflammatory response |
| Module 001 | GO:0098581 | 15.9067669 | 0.000401254 | 0.032999104 | detection of external biotic stimulus |
| Module 001 | GO:0032760 | 14.4097222 | 5.20E-05 | 0.005914925 | positive regulation of tumor necrosis factor production |
| Module 001 | GO:0030890 | 14.3061157 | 0.000306142 | 0.014641312 | positive regulation of B cell proliferation |
| Module 001 | GO:0032728 | 14.2368421 | 0.001816661 | 0.042706923 | positive regulation of interferon-beta production |
| Module 001 | GO:0032722 | 14.2197368 | 0.001822803 | 0.042706923 | positive regulation of chemokine production |
| Module 001 | GO:0042035 | 13.8156553 | 5.80E-05 | 0.008076784 | regulation of cytokine biosynthetic process |
| Module 001 | GO:0031343 | 13.2271892 | 0.000416547 | 0.020225663 | positive regulation of cell killing |

| | | | | | |
|---|---|---|---|---|---|
| Module 001 | GO:0050830 | 13.2097959 | 1.54E-05 | 0.002685807 | defense response to Gram-positive bacterium |
| Module 001 | GO:0042107 | 12.7724057 | 8.24E-05 | 0.008606048 | cytokine metabolic process |
| Module 001 | GO:0032735 | 11.876584 | 0.000623812 | 0.031738525 | positive regulation of interleukin-12 production |
| Module 001 | GO:0032946 | 10.0590985 | 9.51E-07 | 0.000758087 | positive regulation of mononuclear cell proliferation |
| Module 001 | GO:0006576 | 9.32640693 | 0.001396443 | 0.046467831 | cellular biogenic amine metabolic process |
| Module 001 | GO:0031295 | 9.01656051 | 1.15E-05 | 0.004434293 | T cell costimulation |
| Module 001 | GO:0060337 | 8.65817556 | 1.61E-07 | 0.000254229 | type I interferon signaling pathway |
| Module 001 | GO:1903555 | 8.4208464 | 3.68E-06 | 0.001464708 | regulation of tumor necrosis factor superfamily cytokine production |
| Module 001 | GO:0060333 | 7.09325397 | 0.000360451 | 0.021739689 | interferon-gamma-mediated signaling pathway |
| Module 001 | GO:0002821 | 6.80429293 | 0.001266526 | 0.032561965 | positive regulation of adaptive immune response |
| Module 001 | GO:0050670 | 5.9320197 | 4.95E-05 | 0.005914925 | regulation of lymphocyte proliferation |
| Module 001 | GO:0034340 | 5.89413527 | 2.35E-05 | 0.006707156 | response to type I interferon |
| Module 001 | GO:0050729 | 5.76613624 | 0.000966386 | 0.030552721 | positive regulation of inflammatory response |
| Module 001 | GO:0070663 | 5.56902337 | 7.82E-05 | 0.00692483 | regulation of leukocyte proliferation |
| Module 001 | GO:0001817 | 5.26137513 | 0.000731766 | 0.045016018 | regulation of cytokine production |
| Module 001 | GO:0071706 | 5.01412921 | 4.18E-05 | 0.005736734 | tumor necrosis factor superfamily cytokine production |
| Module 001 | GO:0051251 | 4.66282676 | 3.13E-05 | 0.013661489 | positive regulation of lymphocyte activation |
| Module 001 | GO:1903037 | 3.91582987 | 8.28E-05 | 0.016015242 | regulation of leukocyte cell-cell adhesion |
| Module 001 | GO:0006959 | 3.90713646 | 0.000562728 | 0.0489489 | humoral immune response |
| Module 001 | GO:1903039 | 3.8634058 | 0.001803464 | 0.049108779 | positive regulation of leukocyte cell-cell adhesion |
| Module 001 | GO:0009617 | 3.86334373 | 0.000296956 | 0.032867714 | response to bacterium |
| Module 001 | GO:0050863 | 3.82752549 | 0.000107625 | 0.012800257 | regulation of T cell activation |
| Module 001 | GO:0009607 | 3.72776498 | 5.25E-06 | 0.002296172 | response to biotic stimulus |
| Module 001 | GO:0002250 | 3.63795045 | 0.000157195 | 0.031326678 | adaptive immune response |
| Module 001 | GO:0006955 | 3.54850991 | 0.000380118 | 0.014641312 | immune response |
| Module 001 | GO:0045321 | 3.52379605 | 9.50E-05 | 0.021396326 | leukocyte activation |
| Module 001 | GO:0009605 | 3.28640972 | 0.000267366 | 0.016691269 | response to external stimulus |
| Module 001 | GO:0050867 | 3.18842187 | 2.47E-06 | 0.002286333 | positive regulation of cell activation |
| Module 001 | GO:0002764 | 3.15375808 | 0.000385781 | 0.014641312 | immune response-regulating signaling pathway |
| Module 001 | GO:0002253 | 3.03256143 | 0.000555257 | 0.019240855 | activation of immune response |
| Module 001 | GO:0002263 | 3.03065247 | 0.000605203 | 0.0473646 | cell activation involved in immune response |
| Module 001 | GO:0022409 | 2.98111026 | 0.000904322 | 0.048520353 | positive regulation of cell-cell adhesion |
| Module 001 | GO:0006952 | 2.93891987 | 6.12E-06 | 0.002489987 | defense response |
| Module 001 | GO:0002694 | 2.88451616 | 5.42E-05 | 0.008456518 | regulation of leukocyte activation |
| Module 001 | GO:0001775 | 2.55062765 | 3.03E-05 | 0.008444677 | cell activation |
| Module 001 | GO:0002684 | 2.12248069 | 4.27E-05 | 0.008258079 | positive regulation of immune system process |
| Module 002 | GO:0010269 | 199.211009 | 0.000302451 | 0.047317989 | response to selenium ion |
| Module 002 | GO:0042420 | 84 | 0.000836058 | 0.047461193 | dopamine catabolic process |
| Module 002 | GO:0110024 | 84 | 0.000836058 | 0.047461193 | positive regulation of cardiac muscle myoblast proliferation |

| | | | | | |
|---|---|---|---|---|---|
| Module 002 | GO:1902389 | 84 | 0.000836058 | 0.047461193 | ceramide 1-phosphate transport |
| Module 002 | GO:0048484 | 44.8702929 | 0.000201949 | 0.040368894 | enteric nervous system development |
| Module 002 | GO:2001214 | 35.3169935 | 0.000222591 | 0.043360655 | positive regulation of vasculogenesis |
| Module 002 | GO:0043116 | 31.7285156 | 0.000256101 | 0.041795764 | negative regulation of vascular permeability |
| Module 002 | GO:1902667 | 27.3914141 | 0.000338469 | 0.047317989 | regulation of axon guidance |
| Module 002 | GO:0032271 | 16.488177 | 0.000196868 | 0.03996413 | regulation of protein polymerization |
| Module 002 | GO:0050982 | 16.3728037 | 0.000260462 | 0.044734319 | detection of mechanical stimulus |
| Module 002 | GO:0001569 | 14.9501385 | 0.000282318 | 0.044216223 | branching involved in blood vessel morphogenesis |
| Module 002 | GO:0035987 | 10.6358268 | 0.000851168 | 0.047461193 | endodermal cell differentiation |
| Module 002 | GO:0045665 | 10.5513673 | 0.00087175 | 0.047461193 | negative regulation of neuron differentiation |
| Module 002 | GO:0010721 | 7.30693069 | 3.50E-05 | 0.016681262 | negative regulation of cell development |
| Module 002 | GO:0051961 | 6.76208078 | 0.000161652 | 0.025702703 | negative regulation of nervous system development |
| Module 002 | GO:0007369 | 5.41795666 | 9.23E-06 | 0.005347128 | gastrulation |
| Module 002 | GO:0010769 | 4.68173026 | 0.00012695 | 0.041795764 | regulation of cell morphogenesis involved in differentiation |
| Module 002 | GO:0050767 | 4.357908 | 7.86E-05 | 0.018737197 | regulation of neurogenesis |
| Module 002 | GO:0070482 | 3.56395781 | 0.000121417 | 0.032366187 | response to oxygen levels |
| Module 002 | GO:0030030 | 3.2752753 | 7.11E-07 | 0.000976392 | cell projection organization |
| Module 002 | GO:0048858 | 3.23151823 | 0.000193423 | 0.041795764 | cell projection morphogenesis |
| Module 002 | GO:0030182 | 3.22721503 | 0.000321751 | 0.043758158 | neuron differentiation |
| Module 002 | GO:0051674 | 3.07723466 | 0.000351988 | 0.041974605 | localization of cell |
| Module 002 | GO:0001568 | 2.94459724 | 1.96E-07 | 0.000164474 | blood vessel development |
| Module 002 | GO:0072358 | 2.77234627 | 9.70E-08 | 0.000162928 | cardiovascular system development |
| Module 002 | GO:0032990 | 2.67711393 | 7.21E-05 | 0.023380911 | cell part morphogenesis |
| Module 002 | GO:0035295 | 2.53748588 | 0.000205664 | 0.040368894 | tube development |
| Module 002 | GO:0009792 | 2.46198803 | 5.66E-06 | 0.003413205 | embryo development ending in birth or egg hatching |
| Module 002 | GO:0035239 | 2.29207399 | 1.81E-05 | 0.008429367 | tube morphogenesis |
| Module 002 | GO:0007275 | 1.75089802 | 4.43E-05 | 0.016543813 | multicellular organism development |
| Module 002 | GO:0048869 | 1.62369645 | 5.30E-05 | 0.022256817 | cellular developmental process |
| Module 003 | GO:1990809 | 254 | 9.82E-05 | 0.033398564 | endoplasmic reticulum tubular network membrane organization |
| Module 006 | GO:1902969 | Inf | 0.000375611 | 0.021261718 | mitotic DNA replication |
| Module 006 | GO:0017126 | Inf | 0.000378878 | 0.021261718 | nucleologenesis |
| Module 006 | GO:1902298 | Inf | 0.000378878 | 0.021261718 | cell cycle DNA replication maintenance of fidelity |
| Module 006 | GO:1990426 | Inf | 0.000378878 | 0.021261718 | mitotic recombination-dependent replication fork processing |
| Module 006 | GO:0035350 | Inf | 0.001098281 | 0.040376283 | FAD transmembrane transport |
| Module 006 | GO:0051131 | 86.7777778 | 0.000376915 | 0.043345255 | chaperone-mediated protein complex assembly |
| Module 006 | GO:0006391 | 83.3385214 | 0.001640242 | 0.042636597 | transcription initiation from mitochondrial promoter |
| Module 006 | GO:0034501 | 34.0042105 | 0.000281637 | 0.01764081 | protein localization to kinetochore |
| Module 006 | GO:0006362 | 29.9395604 | 0.00021882 | 0.014121956 | transcription elongation from RNA polymerase I promoter |
| Module 006 | GO:0006363 | 28.8280423 | 0.000242437 | 0.014121956 | termination of RNA polymerase I transcription |

| | | | | | |
|---|---|---|---|---|---|
| Module 006 | GO:0006361 | 25.0990783 | 0.000353252 | 0.016461541 | transcription initiation from RNA polymerase I promoter |
| Module 006 | GO:0030490 | 21.1393979 | 0.000809484 | 0.036386303 | maturation of SSU-rRNA |
| Module 006 | GO:0070125 | 11.7633638 | 0.000118015 | 0.01189004 | mitochondrial translational elongation |
| Module 006 | GO:0036297 | 11.3851351 | 0.000150364 | 0.024337566 | interstrand cross-link repair |
| Module 006 | GO:0006521 | 10.0745649 | 1.58E-05 | 0.003461745 | regulation of cellular amino acid metabolic process |
| Module 006 | GO:0090305 | 9.92792912 | 1.84E-05 | 0.004294068 | nucleic acid phosphodiester bond hydrolysis |
| Module 006 | GO:0140053 | 8.8497537 | 0.000114682 | 0.01189004 | mitochondrial gene expression |
| Module 006 | GO:0031145 | 8.65781923 | 3.15E-06 | 0.001322222 | anaphase-promoting complex-dependent catabolic process |
| Module 006 | GO:0044106 | 8.57449971 | 4.03E-05 | 0.005636848 | cellular amine metabolic process |
| Module 006 | GO:0061418 | 8.13030303 | 1.65E-05 | 0.003461745 | regulation of transcription from RNA polymerase II promoter in response to hypoxia |
| Module 006 | GO:1902036 | 7.67398407 | 7.65E-05 | 0.007858873 | regulation of hematopoietic stem cell differentiation |
| Module 006 | GO:0031146 | 7.54538153 | 8.43E-05 | 0.007858873 | SCF-dependent proteasomal ubiquitin-dependent protein catabolic process |
| Module 006 | GO:0006120 | 7.35412335 | 4.04E-05 | 0.003958817 | mitochondrial electron transport, NADH to ubiquinone |
| Module 006 | GO:0010972 | 7.11202652 | 0.000118199 | 0.009015385 | negative regulation of G2/M transition of mitotic cell cycle |
| Module 006 | GO:0002479 | 6.96172382 | 0.000133672 | 0.009345882 | antigen processing and presentation of exogenous peptide antigen via MHC class I, TAP-dependent |
| Module 006 | GO:0032981 | 6.80064103 | 6.55E-05 | 0.005112341 | mitochondrial respiratory chain complex I assembly |
| Module 006 | GO:0006139 | 6.70151199 | 7.34E-09 | 3.95E-06 | nucleobase-containing compound metabolic process |
| Module 006 | GO:0070126 | 6.51155183 | 1.51E-06 | 0.000224528 | mitochondrial translational termination |
| Module 006 | GO:0042773 | 5.72067149 | 1.34E-05 | 0.001663448 | ATP synthesis coupled electron transport |
| Module 006 | GO:0043620 | 5.3478288 | 0.000247424 | 0.015968367 | regulation of DNA-templated transcription in response to stress |
| Module 006 | GO:0002474 | 5.31374911 | 0.00061069 | 0.034157945 | antigen processing and presentation of peptide antigen via MHC class I |
| Module 006 | GO:0006303 | 5.21453117 | 0.000754203 | 0.027972068 | double-strand break repair via nonhomologous end joining |
| Module 006 | GO:0002223 | 5.19062457 | 0.000695002 | 0.035012047 | stimulatory C-type lectin receptor signaling pathway |
| Module 006 | GO:0070498 | 5.13116101 | 0.000740482 | 0.035012047 | interleukin-1-mediated signaling pathway |
| Module 006 | GO:0060071 | 4.80094848 | 0.001065163 | 0.047035346 | Wnt signaling pathway, planar cell polarity pathway |
| Module 006 | GO:0002478 | 4.73610624 | 0.000275419 | 0.011780807 | antigen processing and presentation of exogenous peptide antigen |
| Module 006 | GO:0038095 | 4.73473007 | 0.000534186 | 0.025275447 | Fc-epsilon receptor signaling pathway |
| Module 006 | GO:0006260 | 4.65831537 | 0.000160027 | 0.015252556 | DNA replication |
| Module 006 | GO:0050000 | 4.64737205 | 0.00141193 | 0.037607902 | chromosome localization |
| Module 006 | GO:0008033 | 4.61065705 | 8.64E-05 | 0.008284649 | tRNA processing |
| Module 006 | GO:0046034 | 4.43909774 | 0.000408717 | 0.020030956 | ATP metabolic process |
| Module 006 | GO:0006396 | 4.39078283 | 2.67E-05 | 0.004473871 | RNA processing |
| Module 006 | GO:0000398 | 4.20848633 | 2.94E-07 | 5.20E-05 | mRNA splicing, via spliceosome |
| Module 006 | GO:0071453 | 3.68063499 | 0.000751152 | 0.035012047 | cellular response to oxygen levels |
| Module 006 | GO:0046483 | 3.58175274 | 3.57E-18 | 3.17E-15 | heterocycle metabolic process |
| Module 006 | GO:0000375 | 3.58078603 | 2.98E-08 | 2.27E-05 | RNA splicing, via transesterification reactions |
| Module 006 | GO:0043624 | 3.53047819 | 1.59E-05 | 0.00184619 | cellular protein complex disassembly |
| Module 006 | GO:0006725 | 3.50027859 | 3.98E-18 | 3.17E-15 | cellular aromatic compound metabolic process |
| Module 006 | GO:0034641 | 3.48586898 | 3.40E-16 | 1.35E-13 | cellular nitrogen compound metabolic process |

| | | | | | |
|---|---|---|---|---|---|
| Module 006 | GO:1901360 | 3.32849588 | 2.31E-16 | 1.22E-13 | organic cyclic compound metabolic process |
| Module 006 | GO:0009059 | 3.25633941 | 4.60E-08 | 2.46E-05 | macromolecule biosynthetic process |
| Module 006 | GO:1901796 | 3.20665446 | 0.00127444 | 0.037553499 | regulation of signal transduction by p53 class mediator |
| Module 006 | GO:1901990 | 3.20135566 | 0.000313109 | 0.012974471 | regulation of mitotic cell cycle phase transition |
| Module 006 | GO:0010608 | 3.18146599 | 0.000325286 | 0.016558611 | posttranscriptional regulation of gene expression |
| Module 006 | GO:0043170 | 2.98749218 | 0.000264346 | 0.015814667 | macromolecule metabolic process |
| Module 006 | GO:1902749 | 2.98039216 | 0.000602707 | 0.022833967 | regulation of cell cycle G2/M phase transition |
| Module 006 | GO:1901988 | 2.9769559 | 0.000264252 | 0.011679945 | negative regulation of cell cycle phase transition |
| Module 006 | GO:0043933 | 2.97264891 | 5.48E-05 | 0.004674004 | protein-containing complex subunit organization |
| Module 006 | GO:0009161 | 2.84134838 | 0.000909299 | 0.036153714 | ribonucleoside monophosphate metabolic process |
| Module 006 | GO:0009126 | 2.72467533 | 0.0013191 | 0.038111296 | purine nucleoside monophosphate metabolic process |
| Module 006 | GO:0009141 | 2.49629807 | 0.002043375 | 0.049541408 | nucleoside triphosphate metabolic process |
| Module 006 | GO:0006402 | 2.46534889 | 0.000380368 | 0.020683661 | mRNA catabolic process |
| Module 006 | GO:0043604 | 2.35206756 | 4.22E-06 | 0.000576306 | amide biosynthetic process |
| Module 006 | GO:0009058 | 2.2879163 | 6.22E-05 | 0.007452714 | biosynthetic process |
| Module 006 | GO:0044249 | 2.26904265 | 8.66E-06 | 0.002315103 | cellular biosynthetic process |
| Module 006 | GO:0006518 | 2.26841957 | 3.58E-06 | 0.0009092 | peptide metabolic process |
| Module 006 | GO:0071704 | 2.21096453 | 4.25E-05 | 0.0063484 | organic substance metabolic process |
| Module 006 | GO:0044238 | 2.18066263 | 5.69E-07 | 9.66E-05 | primary metabolic process |
| Module 006 | GO:0006807 | 2.04770226 | 1.56E-07 | 5.97E-05 | nitrogen compound metabolic process |
| Module 006 | GO:0010564 | 1.97120054 | 0.000477673 | 0.031579183 | regulation of cell cycle process |
| Module 006 | GO:0007049 | 1.87169733 | 0.000196231 | 0.010408069 | cell cycle |
| Module 006 | GO:0044260 | 1.46446026 | 0.001954576 | 0.048610641 | cellular macromolecule metabolic process |

**Supplementary Table 3: GO Terms Enriched in Non-Exponentiated Angiogenic Ovarian Cancer Networks Using ALPACA**

| Modules | GOID | OddsRatio | pval | adjPval | GO Terms |
|---|---|---|---|---|---|
| Module 001 | GO:0035520 | 148.829932 | 0.000150064 | 0.023560037 | monoubiquitinated protein deubiquitination |
| Module 002 | GO:0030505 | Inf | 0.000190777 | 0.031319227 | inorganic diphosphate transport |
| Module 002 | GO:0097105 | 364.8 | 3.83E-05 | 0.006580988 | presynaptic membrane assembly |
| Module 002 | GO:0098698 | 156.2857143 | 0.000137073 | 0.011788298 | postsynaptic specialization assembly |
| Module 002 | GO:1904861 | 84.10769231 | 0.000396892 | 0.022755127 | excitatory synapse assembly |
| Module 002 | GO:0042264 | 74.4 | 0.000229165 | 0.046130897 | peptidyl-aspartic acid hydroxylation |
| Module 002 | GO:0036342 | 39.3 | 0.000125745 | 0.037001894 | post-anal tail morphogenesis |
| Module 002 | GO:0010952 | 35.38436214 | 1.24E-05 | 0.007998892 | positive regulation of peptidase activity |
| Module 002 | GO:0051968 | 32.7439759 | 0.000197792 | 0.037001894 | positive regulation of synaptic transmission, glutamatergic |
| Module 002 | GO:0071481 | 31.54678363 | 0.0003072 | 0.047659922 | cellular response to X-ray |
| Module 002 | GO:0046328 | 27.61810345 | 0.000380247 | 0.04681797 | regulation of JNK cascade |
| Module 002 | GO:0061333 | 9.220034247 | 9.57E-05 | 0.02281345 | renal tubule morphogenesis |
| Module 002 | GO:0048598 | 7.701076321 | 0.000230183 | 0.037001894 | embryonic morphogenesis |
| Module 002 | GO:0022414 | 6.092839806 | 1.63E-05 | 0.005997852 | reproductive process |
| Module 002 | GO:0001503 | 5.823660714 | 0.000159525 | 0.034537943 | ossification |
| Module 002 | GO:0030900 | 5.309182332 | 0.000146125 | 0.042895153 | forebrain development |
| Module 002 | GO:0000003 | 3.465940054 | 5.68E-05 | 0.018632994 | reproduction |
| Module 002 | GO:0048736 | 3.444089808 | 0.000102724 | 0.042895153 | appendage development |
| Module 002 | GO:0072358 | 2.712482497 | 3.11E-07 | 0.000517099 | cardiovascular system development |
| Module 002 | GO:0032502 | 2.675913478 | 3.85E-07 | 0.0003266 | developmental process |
| Module 002 | GO:0048880 | 2.57801185 | 0.000164718 | 0.042895153 | sensory system development |
| Module 002 | GO:0001568 | 2.456512679 | 4.12E-05 | 0.021013765 | blood vessel development |
| Module 002 | GO:0007399 | 2.293627213 | 0.000161225 | 0.042895153 | nervous system development |
| Module 002 | GO:0071310 | 2.095974757 | 0.000285106 | 0.040118534 | cellular response to organic substance |
| Module 002 | GO:0009792 | 2.073996553 | 0.000170728 | 0.042895153 | embryo development ending in birth or egg hatching |
| Module 002 | GO:2000026 | 1.970215776 | 8.90E-05 | 0.030966082 | regulation of multicellular organismal development |
| Module 002 | GO:0051716 | 1.962804552 | 3.72E-05 | 0.018632994 | cellular response to stimulus |
| Module 002 | GO:0065007 | 1.703593371 | 0.000146511 | 0.034537943 | biological regulation |
| Module 002 | GO:0051173 | 1.535595462 | 0.000174283 | 0.042895153 | positive regulation of nitrogen compound metabolic process |
| Module 003 | GO:0030042 | 246.4545455 | 0.000199903 | 0.024577633 | actin filament depolymerization |
| Module 003 | GO:0071502 | 244.4269663 | 0.000203121 | 0.024577633 | cellular response to temperature stimulus |
| Module 003 | GO:2001046 | 161.762963 | 0.000178235 | 0.012369518 | positive regulation of integrin-mediated signaling pathway |
| Module 003 | GO:0002024 | 81.46067416 | 0.00066977 | 0.045023439 | diet induced thermogenesis |
| Module 003 | GO:2000463 | 16.20767263 | 1.27E-05 | 0.01843025 | positive regulation of excitatory postsynaptic potential |
| Module 003 | GO:0035023 | 13.56766917 | 6.45E-05 | 0.007458711 | regulation of Rho protein signal transduction |
| Module 003 | GO:0007186 | 12.1372549 | 0.000146448 | 0.02636065 | G protein-coupled receptor signaling pathway |
| Module 003 | GO:0051056 | 9.561602871 | 1.91E-06 | 0.000663048 | regulation of small GTPase mediated signal transduction |

| | | | | | |
|---|---|---|---|---|---|
| Module 003 | GO:0007265 | 8.240937224 | 5.66E-05 | 0.007458711 | Ras protein signal transduction |
| Module 003 | GO:0009966 | 4.227337552 | 0.000334536 | 0.019347329 | regulation of signal transduction |
| Module 003 | GO:0008360 | 4.21388443 | 4.45E-05 | 0.02128438 | regulation of cell shape |
| Module 003 | GO:0072511 | 4.200449062 | 0.000592631 | 0.045023439 | divalent inorganic cation transport |
| Module 003 | GO:0023052 | 2.988370424 | 0.000383111 | 0.038630354 | signaling |
| Module 003 | GO:0015849 | 2.953188136 | 0.00020423 | 0.041615822 | organic acid transport |
| Module 003 | GO:0051272 | 2.333798593 | 4.33E-05 | 0.020096527 | positive regulation of cellular component movement |
| Module 003 | GO:0040017 | 2.328043448 | 0.000208338 | 0.041615822 | positive regulation of locomotion |
| Module 003 | GO:0050801 | 2.088077558 | 0.000184649 | 0.041615822 | ion homeostasis |
| Module 003 | GO:0048870 | 2.069284235 | 3.46E-06 | 0.005393648 | cell motility |
| Module 003 | GO:0010562 | 1.879941435 | 8.62E-05 | 0.028205188 | positive regulation of phosphorus metabolic process |
| Module 003 | GO:0050896 | 1.811069153 | 8.39E-06 | 0.006899509 | response to stimulus |
| Module 003 | GO:0051179 | 1.567938 | 1.18E-05 | 0.010004258 | localization |
| Module 006 | GO:0006777 | 317.1304348 | 4.97E-05 | 0.003747249 | Mo-molybdopterin cofactor biosynthetic process |
| Module 006 | GO:0032324 | 237.826087 | 7.44E-05 | 0.003747249 | molybdopterin cofactor biosynthetic process |
| Module 006 | GO:0051189 | 237.826087 | 7.44E-05 | 0.003747249 | prosthetic group metabolic process |
| Module 006 | GO:0072423 | 123.3028571 | 0.000768821 | 0.017375365 | response to DNA damage checkpoint signaling |
| Module 006 | GO:0072396 | 92.47714286 | 4.10E-05 | 0.001352179 | response to cell cycle checkpoint signaling |
| Module 006 | GO:0002949 | 88.25514403 | 0.001468654 | 0.032091661 | tRNA threonylcarbamoyladenosine modification |
| Module 006 | GO:0035269 | 48.49777778 | 0.001154047 | 0.027600964 | protein O-linked mannosylation |
| Module 006 | GO:0046128 | 28.56293706 | 0.000253745 | 0.008228579 | purine ribonucleoside metabolic process |
| Module 006 | GO:1904874 | 24.50913242 | 8.63E-06 | 0.000349207 | positive regulation of telomerase RNA localization to Cajal body |
| Module 006 | GO:0000387 | 21.29688117 | 1.11E-05 | 0.000431477 | spliceosomal snRNP assembly |
| Module 006 | GO:0009116 | 20.64 | 7.44E-05 | 0.010125751 | nucleoside metabolic process |
| Module 006 | GO:0006369 | 20.59428571 | 1.56E-06 | 8.53E-05 | termination of RNA polymerase II transcription |
| Module 006 | GO:0043101 | 20.58143939 | 0.005682242 | 0.041575069 | purine-containing compound salvage |
| Module 006 | GO:0090670 | 19.92704826 | 2.50E-05 | 0.001022928 | RNA localization to Cajal body |
| Module 006 | GO:1904871 | 17.76666667 | 0.00154908 | 0.035301148 | positive regulation of protein localization to Cajal body |
| Module 006 | GO:0090672 | 17.5 | 3.13E-05 | 0.001038797 | telomerase RNA localization |
| Module 006 | GO:0000245 | 17.1952381 | 0.000157349 | 0.00472095 | spliceosomal complex assembly |
| Module 006 | GO:0008334 | 15.99606493 | 0.001315242 | 0.021807565 | histone mRNA metabolic process |
| Module 006 | GO:1903405 | 15.22714286 | 0.002167931 | 0.045766393 | protein localization to nuclear body |
| Module 006 | GO:1904851 | 15.22714286 | 0.002167931 | 0.045766393 | positive regulation of establishment of protein localization to telomere |
| Module 006 | GO:0006521 | 15.12940631 | 0.001406517 | 0.029233407 | regulation of cellular amino acid metabolic process |
| Module 006 | GO:0006614 | 14.99861111 | 4.17E-08 | 1.83E-05 | SRP-dependent cotranslational protein targeting to membrane |
| Module 006 | GO:0030490 | 14.24866548 | 0.002428458 | 0.047522405 | maturation of SSU-rRNA |
| Module 006 | GO:0001682 | 13.99069767 | 0.000639 | 0.013470803 | tRNA 5'-leader removal |
| Module 006 | GO:0000398 | 13.59649123 | 8.25E-05 | 0.003747249 | mRNA splicing, via spliceosome |
| Module 006 | GO:0072599 | 13.47066167 | 9.72E-08 | 2.13E-05 | establishment of protein localization to endoplasmic reticulum |

| | | | | | |
|---|---|---|---|---|---|
| Module 006 | GO:0000375 | 13.34428958 | 1.99E-05 | 0.003747249 | RNA splicing, via transesterification reactions |
| Module 006 | GO:0006413 | 12.56591781 | 5.52E-22 | 8.33E-19 | translational initiation |
| Module 006 | GO:1902036 | 11.7521645 | 0.002792869 | 0.044614114 | regulation of hematopoietic stem cell differentiation |
| Module 006 | GO:0031146 | 11.56747159 | 0.00291506 | 0.044614114 | SCF-dependent proteasomal ubiquitin-dependent protein catabolic process |
| Module 006 | GO:0016071 | 11.546819 | 0.000189854 | 0.007182806 | mRNA metabolic process |
| Module 006 | GO:0051131 | 10.95233161 | 0.00118618 | 0.02239878 | chaperone-mediated protein complex assembly |
| Module 006 | GO:0070125 | 10.93945504 | 2.57E-13 | 4.06E-11 | mitochondrial translational elongation |
| Module 006 | GO:0006335 | 10.87958656 | 0.001299816 | 0.025667249 | DNA replication-dependent nucleosome assembly |
| Module 006 | GO:0002479 | 10.7243083 | 0.003575347 | 0.044614114 | antigen processing and presentation of exogenous peptide antigen via MHC class I, TAP-dependent |
| Module 006 | GO:0061418 | 10.7243083 | 0.003575347 | 0.044614114 | regulation of transcription from RNA polymerase II promoter in response to hypoxia |
| Module 006 | GO:0002181 | 10.59654258 | 8.80E-08 | 4.57E-06 | cytoplasmic translation |
| Module 006 | GO:0010972 | 10.27462121 | 0.004011952 | 0.047976258 | negative regulation of G2/M transition of mitotic cell cycle |
| Module 006 | GO:0000966 | 10.22047397 | 0.000395032 | 0.008559032 | RNA 5'-end processing |
| Module 006 | GO:0016070 | 10.01196013 | 2.31E-08 | 2.37E-06 | RNA metabolic process |
| Module 006 | GO:0031145 | 9.995085995 | 0.004320295 | 0.049596988 | anaphase-promoting complex-dependent catabolic process |
| Module 006 | GO:0042773 | 9.956801471 | 0.001012277 | 0.014716947 | ATP synthesis coupled electron transport |
| Module 006 | GO:0070126 | 9.885084612 | 4.37E-12 | 6.15E-10 | mitochondrial translational termination |
| Module 006 | GO:0042273 | 9.707720588 | 0.000117302 | 0.003218051 | ribosomal large subunit biogenesis |
| Module 006 | GO:0006139 | 9.611678005 | 0.000409499 | 0.014690788 | nucleobase-containing compound metabolic process |
| Module 006 | GO:0006612 | 9.406824147 | 1.60E-06 | 0.000140746 | protein targeting to membrane |
| Module 006 | GO:0000184 | 9.339871362 | 3.04E-16 | 6.54E-14 | nuclear-transcribed mRNA catabolic process, nonsense-mediated decay |
| Module 006 | GO:1903706 | 8.453781513 | 0.00185123 | 0.033206447 | regulation of hemopoiesis |
| Module 006 | GO:0019083 | 8.331266534 | 1.10E-15 | 2.07E-13 | viral transcription |
| Module 006 | GO:0090090 | 8.259650471 | 1.42E-05 | 0.00103874 | negative regulation of canonical Wnt signaling pathway |
| Module 006 | GO:0002474 | 7.6796875 | 0.002505995 | 0.026312948 | antigen processing and presentation of peptide antigen via MHC class I |
| Module 006 | GO:0044106 | 7.6796875 | 0.002505995 | 0.026312948 | cellular amine metabolic process |
| Module 006 | GO:0140053 | 7.614950839 | 6.90E-11 | 7.28E-09 | mitochondrial gene expression |
| Module 006 | GO:0002223 | 7.507638889 | 0.002709821 | 0.02694897 | stimulatory C-type lectin receptor signaling pathway |
| Module 006 | GO:0070498 | 7.424450549 | 0.002815881 | 0.02694897 | interleukin-1-mediated signaling pathway |
| Module 006 | GO:0006753 | 7.320688444 | 0.003086521 | 0.044614114 | nucleoside phosphate metabolic process |
| Module 006 | GO:0001736 | 7.190921228 | 0.000322514 | 0.007865747 | establishment of planar polarity |
| Module 006 | GO:0006605 | 7.009206074 | 3.94E-09 | 2.77E-07 | protein targeting |
| Module 006 | GO:0060071 | 6.961340206 | 0.003512608 | 0.032384531 | Wnt signaling pathway, planar cell polarity pathway |
| Module 006 | GO:0022613 | 6.859351852 | 0.00014028 | 0.004719767 | ribonucleoprotein complex biogenesis |
| Module 006 | GO:0009144 | 6.743061396 | 0.000499533 | 0.015929564 | purine nucleoside triphosphate metabolic process |
| Module 006 | GO:1901135 | 6.737583893 | 0.000196824 | 0.0109315 | carbohydrate derivative metabolic process |
| Module 006 | GO:0038095 | 6.683168317 | 0.004037225 | 0.034683437 | Fc-epsilon receptor signaling pathway |
| Module 006 | GO:0043620 | 6.683168317 | 0.004037225 | 0.034683437 | regulation of DNA-templated transcription in response to stress |
| Module 006 | GO:0006402 | 6.152999131 | 0.005375832 | 0.043235412 | mRNA catabolic process |

| | | | | | |
|---|---|---|---|---|---|
| Module 006 | GO:0006396 | 6.128722137 | 0.000511777 | 0.01452167 | RNA processing |
| Module 006 | GO:0090263 | 5.966814159 | 0.005924153 | 0.046652702 | positive regulation of canonical Wnt signaling pathway |
| Module 006 | GO:1903311 | 5.873015873 | 0.002320048 | 0.021921282 | regulation of mRNA metabolic process |
| Module 006 | GO:0034440 | 5.805199008 | 0.002446532 | 0.037764415 | lipid oxidation |
| Module 006 | GO:0071826 | 5.795267072 | 0.000146919 | 0.004719767 | ribonucleoprotein complex subunit organization |
| Module 006 | GO:0009199 | 5.534161491 | 0.003187622 | 0.044614114 | ribonucleoside triphosphate metabolic process |
| Module 006 | GO:0008033 | 5.153510896 | 0.000329853 | 0.009393044 | tRNA processing |
| Module 006 | GO:0009451 | 5.152313758 | 0.000764488 | 0.019149495 | RNA modification |
| Module 006 | GO:0090150 | 5.042380952 | 1.46E-06 | 7.25E-05 | establishment of protein localization to membrane |
| Module 006 | GO:0033209 | 4.863800905 | 0.004995597 | 0.039873949 | tumor necrosis factor-mediated signaling pathway |
| Module 006 | GO:0002478 | 4.827471195 | 0.002346998 | 0.048753558 | antigen processing and presentation of exogenous peptide antigen |
| Module 006 | GO:0038061 | 4.826223619 | 0.005153323 | 0.040398374 | NIK/NF-kappaB signaling |
| Module 006 | GO:0042180 | 4.759239517 | 0.005453433 | 0.040769932 | cellular ketone metabolic process |
| Module 006 | GO:0002244 | 4.752764264 | 0.00547933 | 0.040769932 | hematopoietic progenitor cell differentiation |
| Module 006 | GO:0034641 | 4.733167712 | 4.93E-12 | 1.20E-09 | cellular nitrogen compound metabolic process |
| Module 006 | GO:0006163 | 4.700892857 | 0.001428914 | 0.029233407 | purine nucleotide metabolic process |
| Module 006 | GO:0010608 | 4.653571429 | 0.001288109 | 0.015707769 | posttranscriptional regulation of gene expression |
| Module 006 | GO:0043488 | 4.646626298 | 0.005995345 | 0.042450911 | regulation of mRNA stability |
| Module 006 | GO:0043170 | 4.59653741 | 1.02E-06 | 6.17E-05 | macromolecule metabolic process |
| Module 006 | GO:0006303 | 4.594677138 | 0.000756641 | 0.015944293 | double-strand break repair via nonhomologous end joining |
| Module 006 | GO:0044271 | 4.547656308 | 1.37E-08 | 1.37E-06 | cellular nitrogen compound biosynthetic process |
| Module 006 | GO:0071704 | 4.495316159 | 9.57E-07 | 5.56E-05 | organic substance metabolic process |
| Module 006 | GO:0044281 | 4.493404635 | 0.001012316 | 0.026412253 | small molecule metabolic process |
| Module 006 | GO:0006457 | 4.44416499 | 0.000190836 | 0.005294342 | protein folding |
| Module 006 | GO:0043043 | 4.388010333 | 4.00E-09 | 4.64E-07 | peptide biosynthetic process |
| Module 006 | GO:0045333 | 4.340475354 | 0.001902463 | 0.039874081 | cellular respiration |
| Module 006 | GO:0006364 | 4.334660261 | 6.34E-05 | 0.002264548 | rRNA processing |
| Module 006 | GO:0044260 | 4.26435715 | 0.001188396 | 0.024737399 | cellular macromolecule metabolic process |
| Module 006 | GO:0006283 | 4.249193548 | 0.000590089 | 0.014956079 | transcription-coupled nucleotide-excision repair |
| Module 006 | GO:0043624 | 4.229076479 | 8.50E-07 | 4.31E-05 | cellular protein complex disassembly |
| Module 006 | GO:0009059 | 4.223594695 | 9.59E-19 | 3.62E-16 | macromolecule biosynthetic process |
| Module 006 | GO:0034655 | 4.135621977 | 0.000217398 | 0.0091307 | nucleobase-containing compound catabolic process |
| Module 006 | GO:0000209 | 4.082948802 | 0.002598455 | 0.022814432 | protein polyubiquitination |
| Module 006 | GO:0030111 | 3.938050635 | 0.001685826 | 0.017211107 | regulation of Wnt signaling pathway |
| Module 006 | GO:0050000 | 3.938028169 | 0.001898899 | 0.034445153 | chromosome localization |
| Module 006 | GO:0010498 | 3.933300149 | 0.005991015 | 0.042450911 | proteasomal protein catabolic process |
| Module 006 | GO:0006518 | 3.928126374 | 2.60E-19 | 8.11E-17 | peptide metabolic process |
| Module 006 | GO:0043604 | 3.906458018 | 2.10E-19 | 8.11E-17 | amide biosynthetic process |
| Module 006 | GO:0044265 | 3.863422141 | 0.001931156 | 0.019267671 | cellular macromolecule catabolic process |

| | | | | | |
|---|---|---|---|---|---|
| Module 006 | GO:0034660 | 3.842887501 | 1.18E-08 | 8.76E-07 | ncRNA metabolic process |
| Module 006 | GO:0034645 | 3.813146998 | 2.36E-05 | 0.000812877 | cellular macromolecule biosynthetic process |
| Module 006 | GO:0044085 | 3.792461742 | 4.18E-06 | 0.000171667 | cellular component biogenesis |
| Module 006 | GO:0009259 | 3.641904762 | 0.002851743 | 0.02694897 | ribonucleotide metabolic process |
| Module 006 | GO:0043933 | 3.454616994 | 4.70E-06 | 0.000212686 | protein-containing complex subunit organization |
| Module 006 | GO:0046483 | 3.345488758 | 4.17E-14 | 6.51E-12 | heterocycle metabolic process |
| Module 006 | GO:0043603 | 3.320406349 | 1.11E-06 | 5.95E-05 | cellular amide metabolic process |
| Module 006 | GO:0072657 | 3.264798748 | 0.00121789 | 0.015275815 | protein localization to membrane |
| Module 006 | GO:0045930 | 3.2345157 | 0.00077819 | 0.015250508 | negative regulation of mitotic cell cycle |
| Module 006 | GO:0006725 | 3.230881177 | 2.46E-12 | 3.25E-10 | cellular aromatic compound metabolic process |
| Module 006 | GO:0009057 | 3.199257779 | 3.31E-08 | 1.92E-06 | macromolecule catabolic process |
| Module 006 | GO:1901360 | 3.052720813 | 2.02E-11 | 1.95E-09 | organic cyclic compound metabolic process |
| Module 006 | GO:0046700 | 3.048961021 | 2.92E-10 | 2.32E-08 | heterocycle catabolic process |
| Module 006 | GO:0044270 | 3.03518894 | 3.35E-10 | 2.53E-08 | cellular nitrogen compound catabolic process |
| Module 006 | GO:0019439 | 3.021649448 | 2.66E-10 | 2.23E-08 | aromatic compound catabolic process |
| Module 006 | GO:0050852 | 2.99712313 | 0.000394346 | 0.008500981 | T cell receptor signaling pathway |
| Module 006 | GO:0016579 | 2.949002849 | 0.000135469 | 0.00351137 | protein deubiquitination |
| Module 006 | GO:1901361 | 2.878747147 | 1.64E-09 | 1.12E-07 | organic cyclic compound catabolic process |
| Module 006 | GO:1901988 | 2.763287733 | 0.000194291 | 0.004375899 | negative regulation of cell cycle phase transition |
| Module 006 | GO:0006807 | 2.65258642 | 1.06E-05 | 0.000431477 | nitrogen compound metabolic process |
| Module 006 | GO:0033365 | 2.577486118 | 0.006693658 | 0.046643109 | protein localization to organelle |
| Module 006 | GO:1901566 | 2.573481912 | 0.002629215 | 0.026860628 | organonitrogen compound biosynthetic process |
| Module 006 | GO:0044237 | 2.422126489 | 7.36E-05 | 0.003005349 | cellular metabolic process |
| Module 006 | GO:0072594 | 2.419878328 | 0.002703942 | 0.049611464 | establishment of protein localization to organelle |
| Module 006 | GO:0044419 | 2.411947431 | 0.001932988 | 0.027270449 | interspecies interaction between organisms |
| Module 006 | GO:0046907 | 2.371722887 | 0.000429067 | 0.010606002 | intracellular transport |
| Module 006 | GO:0009058 | 2.357243807 | 5.25E-10 | 4.24E-08 | biosynthetic process |
| Module 006 | GO:0044403 | 2.333707097 | 3.64E-07 | 1.83E-05 | symbiont process |
| Module 006 | GO:0090304 | 2.324332559 | 5.60E-09 | 3.66E-07 | nucleic acid metabolic process |
| Module 006 | GO:0044249 | 2.299011319 | 1.90E-09 | 1.35E-07 | cellular biosynthetic process |
| Module 006 | GO:0044238 | 2.187560707 | 2.27E-06 | 0.000103665 | primary metabolic process |
| Module 006 | GO:0070647 | 1.990473752 | 3.96E-05 | 0.001220754 | protein modification by small protein conjugation or removal |
| Module 006 | GO:1901575 | 1.915862615 | 0.001741281 | 0.040259628 | organic substance catabolic process |
| Module 006 | GO:1901576 | 1.815227008 | 0.000168195 | 0.004689269 | organic substance biosynthetic process |
| Module 006 | GO:0065003 | 1.802072757 | 0.000141436 | 0.00351137 | protein-containing complex assembly |
| Module 006 | GO:0051649 | 1.757684158 | 5.34E-06 | 0.000213624 | establishment of localization in cell |
| Module 006 | GO:0070727 | 1.750374119 | 1.82E-05 | 0.000617384 | cellular macromolecule localization |
| Module 006 | GO:0034613 | 1.738062762 | 3.79E-05 | 0.001192892 | cellular protein localization |
| Module 006 | GO:0015833 | 1.616728761 | 0.000192135 | 0.004995519 | peptide transport |

| | | | | | |
|---|---|---|---|---|---|
| Module 006 | GO:0015031 | 1.596304707 | 0.0012014 | 0.02238164 | protein transport |
| Module 006 | GO:0042886 | 1.56351538 | 0.00066048 | 0.013114009 | amide transport |
| Module 006 | GO:0019538 | 1.496161678 | 0.000667871 | 0.018277846 | protein metabolic process |
| Module 006 | GO:0009056 | 1.436215974 | 0.002306825 | 0.040011487 | catabolic process |

**Supplementary Table 4: GO Terms Enriched in Non-Exponentiated Angiogenic Ovarian Cancer Networks Using Permutation Method**

| Modules | GOID | OddsRatio | pval | adjPval | GO Terms |
|---|---|---|---|---|---|
| Module 001 | GO:0002480 | 57.89071038 | 1.13E-07 | 0.00018801 | antigen processing and presentation of exogenous peptide antigen via MHC class I, TAP-independent |
| Module 001 | GO:0060333 | 5.607287449 | 3.94E-05 | 0.0328965 | interferon-gamma-mediated signaling pathway |
| Module 001 | GO:0006955 | 2.247965999 | 1.44E-05 | 0.01247989 | immune response |
| Module 002 | GO:0030505 | Inf | 3.79E-05 | 0.01810161 | inorganic diphosphate transport |
| Module 003 | GO:0044282 | 3.837882043 | 2.94E-05 | 0.03250644 | small molecule catabolic process |
| Module 005 | GO:0060315 | 34.13227092 | 3.28E-05 | 0.02961967 | negative regulation of ryanodine-sensitive calcium-release channel activity |
| Module 005 | GO:0051284 | 24.3756403 | 8.28E-05 | 0.02961967 | positive regulation of sequestering of calcium ion |
| Module 005 | GO:0010523 | 21.32669323 | 0.00012 | 0.02961967 | negative regulation of calcium ion transport into cytosol |
| Module 005 | GO:0009260 | 6.593596059 | 0.00019 | 0.03903004 | ribonucleotide biosynthetic process |
| Module 005 | GO:0009126 | 5.090838809 | 6.76E-05 | 0.02762919 | purine nucleoside monophosphate metabolic process |
| Module 005 | GO:0009161 | 5.085040588 | 6.82E-05 | 0.02762919 | ribonucleoside monophosphate metabolic process |
| Module 005 | GO:0019693 | 4.799451804 | 0.00013 | 0.03903004 | ribose phosphate metabolic process |
| Module 005 | GO:0009259 | 2.884607371 | 7.90E-05 | 0.02762919 | ribonucleotide metabolic process |
| Module 006 | GO:0002949 | 108.2110553 | 0.00099 | 0.03097251 | tRNA threonylcarbamoyladenosine modification |
| Module 006 | GO:0021796 | 89.00414938 | 0.00144 | 0.03507176 | cerebral cortex regionalization |
| Module 006 | GO:0051256 | 83.34615385 | 0.00085 | 0.03847969 | mitotic spindle midzone assembly |
| Module 006 | GO:0001302 | 68.28930818 | 0.00094 | 0.04091405 | replicative cell aging |
| Module 006 | GO:0019370 | 43.23809524 | 0.00151 | 0.03037754 | leukotriene biosynthetic process |
| Module 006 | GO:0009208 | 34.58412698 | 0.0022 | 0.03798541 | pyrimidine ribonucleoside triphosphate metabolic process |
| Module 006 | GO:0006123 | 30.5 | 3.91E-06 | 0.00029772 | mitochondrial electron transport, cytochrome c to oxygen |
| Module 006 | GO:0009148 | 28.81481481 | 0.00301 | 0.04216539 | pyrimidine nucleoside triphosphate biosynthetic process |
| Module 006 | GO:1901800 | 26.44878049 | 0.00334 | 0.04381335 | positive regulation of proteasomal protein catabolic process |
| Module 006 | GO:0031145 | 24.13787991 | 5.61E-07 | 0.00018898 | anaphase-promoting complex-dependent catabolic process |
| Module 006 | GO:0001682 | 20.40310078 | 3.63E-05 | 0.00143409 | tRNA 5'-leader removal |
| Module 006 | GO:0070125 | 17.8287311 | 1.89E-05 | 0.00238711 | mitochondrial translational elongation |
| Module 006 | GO:0070126 | 17.37980769 | 2.13E-05 | 0.00238711 | mitochondrial translational termination |
| Module 006 | GO:0006521 | 16.24091588 | 0.00116 | 0.03258673 | regulation of cellular amino acid metabolic process |
| Module 006 | GO:0061418 | 15.96617647 | 0.00019 | 0.01280381 | regulation of transcription from RNA polymerase II promoter in response to hypoxia |
| Module 006 | GO:0038061 | 15.27311321 | 0.00137 | 0.0356291 | NIK/NF-kappaB signaling |
| Module 006 | GO:0006120 | 14.65469229 | 1.36E-11 | 2.21E-09 | mitochondrial electron transport, NADH to ubiquinone |
| Module 006 | GO:1902036 | 12.6155633 | 0.00231 | 0.04053341 | regulation of hematopoietic stem cell differentiation |
| Module 006 | GO:0031146 | 12.41730183 | 0.00241 | 0.04053341 | SCF-dependent proteasomal ubiquitin-dependent protein catabolic process |
| Module 006 | GO:0032981 | 11.95568182 | 1.26E-10 | 1.86E-08 | mitochondrial respiratory chain complex I assembly |
| Module 006 | GO:0042773 | 11.94546265 | 2.10E-15 | 1.16E-12 | ATP synthesis coupled electron transport |
| Module 006 | GO:0070498 | 11.90549451 | 0.00055 | 0.02494549 | interleukin-1-mediated signaling pathway |
| Module 006 | GO:0002479 | 11.51219512 | 0.00296 | 0.04381335 | antigen processing and presentation of exogenous peptide antigen via MHC class I, TAP-dependent |

| | | | | | |
|---|---|---|---|---|---|
| Module 006 | GO:0010972 | 11.02947154 | 0.00333 | 0.04381335 | negative regulation of G2/M transition of mitotic cell cycle |
| Module 006 | GO:0140053 | 10.90046437 | 0.00017 | 0.01280381 | mitochondrial gene expression |
| Module 006 | GO:0009060 | 10.7177082 | 4.28E-06 | 0.00021808 | aerobic respiration |
| Module 006 | GO:0043620 | 10.71683168 | 0.0008 | 0.03002489 | regulation of DNA-templated transcription in response to stress |
| Module 006 | GO:1903311 | 10.0691602 | 0.00101 | 0.03080755 | regulation of mRNA metabolic process |
| Module 006 | GO:0002758 | 9.123076923 | 0.00138 | 0.03037754 | innate immune response-activating signal transduction |
| Module 006 | GO:0000715 | 9.098339719 | 0.00173 | 0.04505085 | nucleotide-excision repair, DNA damage recognition |
| Module 006 | GO:0001736 | 8.330246914 | 0.00054 | 0.02068518 | establishment of planar polarity |
| Module 006 | GO:0043624 | 8.258866882 | 0.00059 | 0.02494549 | cellular protein complex disassembly |
| Module 006 | GO:0002474 | 8.059612519 | 0.00212 | 0.03798541 | antigen processing and presentation of peptide antigen via MHC class I |
| Module 006 | GO:0044106 | 8.059612519 | 0.00212 | 0.03798541 | cellular amine metabolic process |
| Module 006 | GO:0002223 | 7.879052823 | 0.0023 | 0.03798541 | stimulatory C-type lectin receptor signaling pathway |
| Module 006 | GO:0043488 | 7.874452555 | 0.00238 | 0.04053341 | regulation of mRNA stability |
| Module 006 | GO:1903706 | 7.771554901 | 0.00242 | 0.03859812 | regulation of hemopoiesis |
| Module 006 | GO:0016579 | 7.612375533 | 0.00028 | 0.01546336 | protein deubiquitination |
| Module 006 | GO:0000209 | 7.466041171 | 0.00093 | 0.03080755 | protein polyubiquitination |
| Module 006 | GO:0006614 | 7.309236948 | 1.90E-10 | 2.62E-08 | SRP-dependent cotranslational protein targeting to membrane |
| Module 006 | GO:0060071 | 7.305729255 | 0.00298 | 0.04216539 | Wnt signaling pathway, planar cell polarity pathway |
| Module 006 | GO:0006413 | 7.143501913 | 1.32E-11 | 2.42E-09 | translational initiation |
| Module 006 | GO:0070555 | 7.0875 | 0.00343 | 0.04381335 | response to interleukin-1 |
| Module 006 | GO:0038095 | 7.013796462 | 0.00343 | 0.04559965 | Fc-epsilon receptor signaling pathway |
| Module 006 | GO:0090090 | 6.906410256 | 0.0012 | 0.02774349 | negative regulation of canonical Wnt signaling pathway |
| Module 006 | GO:0072599 | 6.460302749 | 1.18E-09 | 1.30E-07 | establishment of protein localization to endoplasmic reticulum |
| Module 006 | GO:0090502 | 6.365498652 | 4.45E-05 | 0.00167657 | RNA phosphodiester bond hydrolysis, endonucleolytic |
| Module 006 | GO:0000184 | 6.239181287 | 5.21E-06 | 0.00059056 | nuclear-transcribed mRNA catabolic process, nonsense-mediated decay |
| Module 006 | GO:0046034 | 5.635246996 | 5.37E-06 | 0.0003322 | ATP metabolic process |
| Module 006 | GO:0006612 | 5.591236587 | 4.26E-09 | 3.36E-07 | protein targeting to membrane |
| Module 006 | GO:0006406 | 5.583842144 | 0.00047 | 0.02456702 | mRNA export from nucleus |
| Module 006 | GO:0006412 | 5.4408849 | 0.00351 | 0.04381335 | translation |
| Module 006 | GO:1902600 | 5.351935646 | 0.00026 | 0.0087045 | proton transmembrane transport |
| Module 006 | GO:0006511 | 5.349311576 | 0.00162 | 0.03908765 | ubiquitin-dependent protein catabolic process |
| Module 006 | GO:0036294 | 5.325396825 | 0.00355 | 0.04559965 | cellular response to decreased oxygen levels |
| Module 006 | GO:0022900 | 5.272592957 | 0.00027 | 0.01298507 | electron transport chain |
| Module 006 | GO:1903047 | 5.030882353 | 0.0022 | 0.04053341 | mitotic cell cycle process |
| Module 006 | GO:0071426 | 5.00855615 | 0.00085 | 0.03723295 | ribonucleoprotein complex export from nucleus |
| Module 006 | GO:0015980 | 4.935951379 | 1.94E-07 | 3.19E-05 | energy derivation by oxidation of organic compounds |
| Module 006 | GO:0019083 | 4.88336784 | 9.22E-05 | 0.00522555 | viral transcription |
| Module 006 | GO:1901990 | 4.871259782 | 0.00253 | 0.04053341 | regulation of mitotic cell cycle phase transition |

| | | | | | |
|---|---|---|---|---|---|
| Module 006 | GO:0006163 | 4.746446518 | 1.52E-09 | 1.58E-07 | purine nucleotide metabolic process |
| Module 006 | GO:0009259 | 4.658260234 | 4.09E-09 | 4.74E-07 | ribonucleotide metabolic process |
| Module 006 | GO:0043632 | 4.465972729 | 0.00194 | 0.04053341 | modification-dependent macromolecule catabolic process |
| Module 006 | GO:0006402 | 4.432989691 | 0.0002 | 0.01513294 | mRNA catabolic process |
| Module 006 | GO:0043933 | 4.416492329 | 6.76E-08 | 1.26E-05 | protein-containing complex subunit organization |
| Module 006 | GO:0043043 | 4.36240367 | 5.71E-08 | 7.76E-06 | peptide biosynthetic process |
| Module 006 | GO:0043603 | 4.272457176 | 4.29E-11 | 4.58E-08 | cellular amide metabolic process |
| Module 006 | GO:0006283 | 4.170758158 | 0.00126 | 0.04039623 | transcription-coupled nucleotide-excision repair |
| Module 006 | GO:0034641 | 4.012248539 | 6.10E-18 | 1.01E-14 | cellular nitrogen compound metabolic process |
| Module 006 | GO:0044271 | 3.918293998 | 4.02E-10 | 1.43E-07 | cellular nitrogen compound biosynthetic process |
| Module 006 | GO:0002478 | 3.841571195 | 0.00114 | 0.03222915 | antigen processing and presentation of exogenous peptide antigen |
| Module 006 | GO:0009161 | 3.838368223 | 9.30E-06 | 0.00051123 | ribonucleoside monophosphate metabolic process |
| Module 006 | GO:0009126 | 3.783968368 | 1.93E-05 | 0.00091203 | purine nucleoside monophosphate metabolic process |
| Module 006 | GO:0006401 | 3.702773498 | 1.13E-09 | 1.98E-07 | RNA catabolic process |
| Module 006 | GO:0009141 | 3.654930442 | 1.67E-05 | 0.00085307 | nucleoside triphosphate metabolic process |
| Module 006 | GO:0044770 | 3.62671875 | 0.00148 | 0.04759399 | cell cycle phase transition |
| Module 006 | GO:0034655 | 3.576300176 | 1.02E-06 | 0.00012562 | nucleobase-containing compound catabolic process |
| Module 006 | GO:0032446 | 3.559821429 | 0.00241 | 0.04053341 | protein modification by small protein conjugation |
| Module 006 | GO:0006139 | 3.496170767 | 6.20E-05 | 0.00401797 | nucleobase-containing compound metabolic process |
| Module 006 | GO:0022411 | 3.389159364 | 0.00012 | 0.00807209 | cellular component disassembly |
| Module 006 | GO:0090305 | 3.321408518 | 2.08E-06 | 9.87E-05 | nucleic acid phosphodiester bond hydrolysis |
| Module 006 | GO:0044265 | 3.318516043 | 5.27E-05 | 0.00217196 | cellular macromolecule catabolic process |
| Module 006 | GO:0051168 | 3.310518053 | 0.0006 | 0.01930356 | nuclear export |
| Module 006 | GO:0034660 | 3.274904561 | 0.0001 | 0.00362479 | ncRNA metabolic process |
| Module 006 | GO:0010498 | 3.263134058 | 0.00109 | 0.04512611 | proteasomal protein catabolic process |
| Module 006 | GO:0010608 | 3.231187386 | 0.00116 | 0.03195479 | posttranscriptional regulation of gene expression |
| Module 006 | GO:1901988 | 3.168223867 | 0.00053 | 0.01621078 | negative regulation of cell cycle phase transition |
| Module 006 | GO:0046483 | 3.159825654 | 1.94E-14 | 5.35E-12 | heterocycle metabolic process |
| Module 006 | GO:1901360 | 3.034303115 | 1.34E-13 | 2.77E-11 | organic cyclic compound metabolic process |
| Module 006 | GO:0006725 | 3.012802418 | 1.18E-13 | 2.77E-11 | cellular aromatic compound metabolic process |
| Module 006 | GO:0006091 | 2.969000023 | 5.24E-08 | 3.10E-06 | generation of precursor metabolites and energy |
| Module 006 | GO:0046700 | 2.959532531 | 2.13E-09 | 3.35E-07 | heterocycle catabolic process |
| Module 006 | GO:0044270 | 2.946355685 | 2.41E-09 | 3.46E-07 | cellular nitrogen compound catabolic process |
| Module 006 | GO:0055086 | 2.90358785 | 7.88E-07 | 4.26E-05 | nucleobase-containing small molecule metabolic process |
| Module 006 | GO:0044237 | 2.852865014 | 2.38E-05 | 0.00107072 | cellular metabolic process |
| Module 006 | GO:0044281 | 2.852820071 | 0.00042 | 0.01239024 | small molecule metabolic process |
| Module 006 | GO:0019439 | 2.834395898 | 1.39E-08 | 1.44E-06 | aromatic compound catabolic process |
| Module 006 | GO:1901361 | 2.811029015 | 1.25E-08 | 1.41E-06 | organic cyclic compound catabolic process |

| | | | | | |
|---|---|---|---|---|---|
| Module 006 | GO:0006396 | 2.659371092 | 2.13E-05 | 0.00088196 | RNA processing |
| Module 006 | GO:1901564 | 2.632011759 | 5.26E-06 | 0.00022942 | organonitrogen compound metabolic process |
| Module 006 | GO:0016071 | 2.578940119 | 1.63E-07 | 1.17E-05 | mRNA metabolic process |
| Module 006 | GO:0051169 | 2.534314544 | 0.0013 | 0.03671222 | nuclear transport |
| Module 006 | GO:0055114 | 2.155475377 | 7.61E-06 | 0.00032346 | oxidation-reduction process |
| Module 006 | GO:1901576 | 2.13187399 | 3.52E-06 | 0.00019309 | organic substance biosynthetic process |
| Module 006 | GO:0071704 | 2.084152726 | 0.00033 | 0.01195207 | organic substance metabolic process |
| Module 006 | GO:0019637 | 2.050038437 | 0.00174 | 0.04641214 | organophosphate metabolic process |
| Module 006 | GO:0009057 | 2.044633369 | 1.35E-06 | 7.63E-05 | macromolecule catabolic process |
| Module 006 | GO:0044238 | 1.829249636 | 1.09E-06 | 5.50E-05 | primary metabolic process |
| Module 006 | GO:1901575 | 1.786306471 | 0.0016 | 0.04029404 | organic substance catabolic process |
| Module 006 | GO:0006807 | 1.620654975 | 0.00121 | 0.03677931 | nitrogen compound metabolic process |
| Module 006 | GO:0065003 | 1.61657242 | 0.00162 | 0.04536901 | protein-containing complex assembly |
| Module 006 | GO:0009059 | 1.49368488 | 4.00E-05 | 0.00154133 | macromolecule biosynthetic process |
| Module 006 | GO:0009058 | 1.450005374 | 9.56E-05 | 0.00337404 | biosynthetic process |
| Module 006 | GO:0090304 | 1.444272997 | 0.00154 | 0.04775579 | nucleic acid metabolic process |

**Supplementary Table 5: GO Terms Enriched in Non-Exponentiated Angiogenic Ovarian Cancer Networks Using SBM**

| Modules | GOID | OddsRatio | pval | adjPval | GO Terms |
|---|---|---|---|---|---|
| Module 001 | GO:0016339 | 61.22689076 | 0.0007 | 0.043281604 | calcium-dependent cell-cell adhesion via plasma membrane cell adhesion molecules |
| Module 001 | GO:0002480 | 53.77008032 | 7.48E-07 | 0.00029684 | antigen processing and presentation of exogenous peptide antigen via MHC class I, TAP-independent |
| Module 001 | GO:0007156 | 35.78271605 | 1.19E-05 | 0.001465153 | homophilic cell adhesion via plasma membrane adhesion molecules |
| Module 001 | GO:0060337 | 8.228350266 | 1.83E-05 | 0.00525177 | type I interferon signaling pathway |
| Module 001 | GO:0034340 | 5.985356238 | 2.08E-05 | 0.00525177 | response to type I interferon |
| Module 001 | GO:0060333 | 5.915343915 | 0.00015 | 0.030658081 | interferon-gamma-mediated signaling pathway |
| Module 002 | GO:0030505 | Inf | 6.36E-05 | 0.023891388 | inorganic diphosphate transport |
| Module 002 | GO:0043116 | 121.5333333 | 0.00021 | 0.00919199 | negative regulation of vascular permeability |
| Module 002 | GO:1902667 | 91.125 | 0.00034 | 0.010823721 | regulation of axon guidance |
| Module 002 | GO:0007163 | 33.45510836 | 0.00216 | 0.029647009 | establishment or maintenance of cell polarity |
| Module 002 | GO:0001541 | 30.30833333 | 0.00258 | 0.031583596 | ovarian follicle development |
| Module 002 | GO:0050771 | 30.30833333 | 0.00258 | 0.031583596 | negative regulation of axonogenesis |
| Module 002 | GO:0072091 | 23.69782609 | 0.0041 | 0.037393673 | regulation of stem cell proliferation |
| Module 002 | GO:0050773 | 23.29108485 | 0.00425 | 0.037393673 | regulation of dendrite development |
| Module 002 | GO:0046660 | 21.44802632 | 0.00058 | 0.015947671 | female sex differentiation |
| Module 002 | GO:0043297 | 20.17222222 | 0.00554 | 0.045045783 | apical junction assembly |
| Module 002 | GO:0050772 | 19.80363636 | 0.00573 | 0.045045783 | positive regulation of axonogenesis |
| Module 002 | GO:0051149 | 18.145 | 0.00675 | 0.049513189 | positive regulation of muscle cell differentiation |
| Module 002 | GO:0022414 | 17.73459384 | 2.95E-05 | 0.004110558 | reproductive process |
| Module 002 | GO:0010769 | 16.38829625 | 3.74E-05 | 0.004110558 | regulation of cell morphogenesis involved in differentiation |
| Module 002 | GO:0008064 | 15.13833256 | 0.00153 | 0.024347629 | regulation of actin polymerization or depolymerization |
| Module 002 | GO:0040017 | 11.76031215 | 0.00018 | 0.00919199 | positive regulation of locomotion |
| Module 002 | GO:0045137 | 11.36526316 | 0.00337 | 0.034223632 | development of primary sexual characteristics |
| Module 002 | GO:0010720 | 10.9338843 | 6.57E-05 | 0.004818245 | positive regulation of cell development |
| Module 002 | GO:0007015 | 9.280381944 | 0.00161 | 0.024347629 | actin filament organization |
| Module 002 | GO:0051961 | 8.844572368 | 0.00664 | 0.049513189 | negative regulation of nervous system development |
| Module 002 | GO:0051272 | 8.00409339 | 0.00091 | 0.022366278 | positive regulation of cellular component movement |
| Module 002 | GO:0051962 | 7.814862622 | 0.003 | 0.034223632 | positive regulation of nervous system development |
| Module 002 | GO:2000145 | 7.71835206 | 0.00115 | 0.024347629 | regulation of cell motility |
| Module 002 | GO:0120035 | 7.525824964 | 0.00342 | 0.034223632 | regulation of plasma membrane bounded cell projection organization |
| Module 002 | GO:0120039 | 7.317480184 | 0.00134 | 0.024347629 | plasma membrane bounded cell projection morphogenesis |
| Module 002 | GO:0032990 | 6.957472178 | 0.00166 | 0.024347629 | cell part morphogenesis |
| Module 002 | GO:0048514 | 6.521806612 | 0.00558 | 0.045045783 | blood vessel morphogenesis |
| Module 002 | GO:0045595 | 6.333958724 | 0.00027 | 0.010045782 | regulation of cell differentiation |
| Module 002 | GO:0000902 | 5.816742081 | 0.00162 | 0.024347629 | cell morphogenesis |
| Module 002 | GO:0022603 | 5.019382392 | 0.00324 | 0.034223632 | regulation of anatomical structure morphogenesis |
| Module 002 | GO:2000026 | 4.438355254 | 0.0038 | 0.036338501 | regulation of multicellular organismal development |
| Module 003 | GO:0006103 | 41.30506329 | 0.00011 | 0.031751745 | 2-oxoglutarate metabolic process |
| Module 003 | GO:0045017 | 6.119262119 | 0.00011 | 0.031751745 | glycerolipid biosynthetic process |
| Module 006 | GO:0017126 | Inf | 0.00019 | 0.015088912 | nucleologenesis |
| Module 006 | GO:0000447 | 335.4153846 | 0.00011 | 0.037865555 | endonucleolytic cleavage in ITS1 to separate SSU-rRNA from 5.8S rRNA and LSU-rRNA from tricistronic rRNA transcript (SSU-rRNA, 5.8S rRNA, LSU-rRNA) |
| Module 006 | GO:0000480 | 167.6923077 | 0.00022 | 0.037865555 | endonucleolytic cleavage in 5'-ETS of tricistronic rRNA transcript (SSU-rRNA, 5.8S rRNA, LSU-rRNA) |
| Module 006 | GO:0000478 | 111.7846154 | 0.00036 | 0.037865555 | endonucleolytic cleavage involved in rRNA processing |
| Module 006 | GO:0000460 | 31.24137931 | 0.00281 | 0.04689458 | maturation of 5.8S rRNA |
| Module 006 | GO:0019370 | 31.24137931 | 0.00281 | 0.04689458 | leukotriene biosynthetic process |
| Module 006 | GO:0030490 | 27.76756066 | 0.00341 | 0.049675384 | maturation of SSU-rRNA |
| Module 006 | GO:0034501 | 27.76756066 | 0.00341 | 0.049675384 | protein localization to kinetochore |
| Module 006 | GO:0070125 | 22.64166667 | 0.0005 | 0.026640119 | mitochondrial translational elongation |
| Module 006 | GO:0070126 | 22.08536585 | 0.00054 | 0.026640119 | mitochondrial translational termination |

| | | | | | |
|---|---|---|---|---|---|
| Module 006 | GO:0006414 | 19 | 0.00011 | 0.014608562 | translational elongation |
| Module 006 | GO:0019827 | 15.03888889 | 0.00158 | 0.046736419 | stem cell population maintenance |
| Module 006 | GO:0140053 | 13.97803618 | 0.00193 | 0.047374527 | mitochondrial gene expression |
| Module 006 | GO:0042276 | 13.84485447 | 9.61E-05 | 0.004720425 | error-prone translesion synthesis |
| Module 006 | GO:0022900 | 13.25 | 0.00224 | 0.047374527 | electron transport chain |
| Module 006 | GO:0031145 | 13.18292683 | 2.77E-06 | 0.000990068 | anaphase-promoting complex-dependent catabolic process |
| Module 006 | GO:0070987 | 11.03859104 | 0.001 | 0.031342843 | error-free translesion synthesis |
| Module 006 | GO:0006521 | 10.61960784 | 0.00081 | 0.025504021 | regulation of cellular amino acid metabolic process |
| Module 006 | GO:0031146 | 10.3859447 | 0.0002 | 0.014608562 | SCF-dependent proteasomal ubiquitin-dependent protein catabolic process |
| Module 006 | GO:0038061 | 9.843406593 | 0.00106 | 0.029553331 | NIK/NF-kappaB signaling |
| Module 006 | GO:0061418 | 9.606432125 | 0.00029 | 0.017939254 | regulation of transcription from RNA polymerase II promoter in response to hypoxia |
| Module 006 | GO:0032201 | 9.498661909 | 0.00041 | 0.016381129 | telomere maintenance via semi-conservative replication |
| Module 006 | GO:0010972 | 9.192176871 | 0.00035 | 0.019298904 | negative regulation of G2/M transition of mitotic cell cycle |
| Module 006 | GO:0044106 | 9.177721088 | 0.00135 | 0.033908066 | cellular amine metabolic process |
| Module 006 | GO:0006120 | 8.531731732 | 1.47E-05 | 0.001467027 | mitochondrial electron transport, NADH to ubiquinone |
| Module 006 | GO:1902036 | 8.211005693 | 0.00199 | 0.038333235 | regulation of hematopoietic stem cell differentiation |
| Module 006 | GO:0043043 | 8.07040619 | 0.00034 | 0.026640119 | peptide biosynthetic process |
| Module 006 | GO:0042773 | 7.993233083 | 2.20E-07 | 5.26E-05 | ATP synthesis coupled electron transport |
| Module 006 | GO:0043603 | 7.661995178 | 0.00123 | 0.04565283 | cellular amide metabolic process |
| Module 006 | GO:0002479 | 7.482352941 | 0.00274 | 0.04689458 | antigen processing and presentation of exogenous peptide antigen via MHC class I, TAP-dependent |
| Module 006 | GO:0002478 | 7.295238095 | 0.00299 | 0.048261849 | antigen processing and presentation of exogenous peptide antigen |
| Module 006 | GO:0006296 | 7.238230519 | 0.0011 | 0.040019941 | nucleotide-excision repair, DNA incision, 5'-to lesion |
| Module 006 | GO:0070498 | 7.136243386 | 0.00103 | 0.029553331 | interleukin-1-mediated signaling pathway |
| Module 006 | GO:0044786 | 7.100567992 | 4.85E-05 | 0.003098007 | cell cycle DNA replication |
| Module 006 | GO:0006614 | 7.062950648 | 2.05E-09 | 5.97E-07 | SRP-dependent cotranslational protein targeting to membrane |
| Module 006 | GO:0032981 | 6.709567096 | 0.00019 | 0.009399158 | mitochondrial respiratory chain complex I assembly |
| Module 006 | GO:0006406 | 6.615368852 | 0.00049 | 0.027403542 | mRNA export from nucleus |
| Module 006 | GO:0043620 | 6.416666667 | 0.0016 | 0.03493105 | regulation of DNA-templated transcription in response to stress |
| Module 006 | GO:0006301 | 6.382953181 | 0.0007 | 0.024764793 | postreplication repair |
| Module 006 | GO:0072599 | 6.262997347 | 1.03E-08 | 1.98E-06 | establishment of protein localization to endoplasmic reticulum |
| Module 006 | GO:0000184 | 6.108276644 | 0.00197 | 0.038333235 | nuclear-transcribed mRNA catabolic process, nonsense-mediated decay |
| Module 006 | GO:1903364 | 6.108276644 | 0.00197 | 0.038333235 | positive regulation of cellular protein catabolic process |
| Module 006 | GO:0006413 | 6.088578089 | 1.42E-08 | 5.27E-06 | translational initiation |
| Module 006 | GO:0071426 | 5.941425677 | 0.00084 | 0.039872436 | ribonucleoprotein complex export from nucleus |
| Module 006 | GO:0090263 | 5.722789116 | 0.00258 | 0.04689458 | positive regulation of canonical Wnt signaling pathway |
| Module 006 | GO:0034641 | 5.473090278 | 0.00017 | 0.043715314 | cellular nitrogen compound metabolic process |
| Module 006 | GO:0002181 | 5.459521687 | 0.00067 | 0.029054597 | cytoplasmic translation |
| Module 006 | GO:0045930 | 5.447837672 | 0.00316 | 0.049521026 | negative regulation of mitotic cell cycle |
| Module 006 | GO:0006612 | 5.352242424 | 3.73E-08 | 5.22E-06 | protein targeting to membrane |
| Module 006 | GO:0006283 | 5.095728712 | 0.00081 | 0.03472094 | transcription-coupled nucleotide-excision repair |
| Module 006 | GO:0006402 | 5.056029685 | 0.00017 | 0.014608562 | mRNA catabolic process |
| Module 006 | GO:0043624 | 4.743191408 | 5.93E-06 | 0.001195608 | cellular protein complex disassembly |
| Module 006 | GO:1901988 | 4.702162434 | 0.00121 | 0.031917412 | negative regulation of cell cycle phase transition |
| Module 006 | GO:0000082 | 4.631313131 | 0.00141 | 0.040098295 | G1/S transition of mitotic cell cycle |
| Module 006 | GO:0046034 | 4.347107438 | 0.00012 | 0.008667992 | ATP metabolic process |
| Module 006 | GO:0009199 | 4.309415049 | 3.33E-05 | 0.002504807 | ribonucleoside triphosphate metabolic process |
| Module 006 | GO:0019083 | 4.239340306 | 7.38E-05 | 0.004652145 | viral transcription |
| Module 006 | GO:1903311 | 4.081278027 | 0.00069 | 0.031459541 | regulation of mRNA metabolic process |
| Module 006 | GO:1901360 | 4.039254824 | 0.00032 | 0.043715314 | organic cyclic compound metabolic process |
| Module 006 | GO:0009144 | 4.038001784 | 6.13E-05 | 0.003406068 | purine nucleoside triphosphate metabolic process |
| Module 006 | GO:0044265 | 4.027868277 | 0.00152 | 0.034576419 | cellular macromolecule catabolic process |

| | | | | | |
|---|---|---|---|---|---|
| Module 006 | GO:0043488 | 3.945827475 | 0.00086 | 0.034912633 | regulation of mRNA stability |
| Module 006 | GO:1901990 | 3.935999403 | 0.00051 | 0.021594484 | regulation of mitotic cell cycle phase transition |
| Module 006 | GO:0006725 | 3.892177115 | 0.0004 | 0.043715314 | cellular aromatic compound metabolic process |
| Module 006 | GO:0009896 | 3.676973684 | 0.00145 | 0.034516281 | positive regulation of catabolic process |
| Module 006 | GO:0015980 | 3.480489418 | 8.22E-05 | 0.006614041 | energy derivation by oxidation of organic compounds |
| Module 006 | GO:0016579 | 3.352240138 | 0.00012 | 0.011928713 | protein deubiquitination |
| Module 006 | GO:0034655 | 3.340206186 | 0.00065 | 0.023131951 | nucleobase-containing compound catabolic process |
| Module 006 | GO:0043933 | 3.323280423 | 5.24E-06 | 0.000447795 | protein-containing complex subunit organization |
| Module 006 | GO:0090305 | 3.183624155 | 0.00051 | 0.02640395 | nucleic acid phosphodiester bond hydrolysis |
| Module 006 | GO:0009259 | 3.041962175 | 0.00077 | 0.025214901 | ribonucleotide metabolic process |
| Module 006 | GO:0043632 | 2.99635466 | 0.00347 | 0.049675384 | modification-dependent macromolecule catabolic process |
| Module 006 | GO:0000375 | 2.917717528 | 0.00048 | 0.02640395 | RNA splicing, via transesterification reactions |
| Module 006 | GO:0046483 | 2.898711871 | 4.18E-10 | 2.14E-07 | heterocycle metabolic process |
| Module 006 | GO:0009161 | 2.89864155 | 0.00051 | 0.024964093 | ribonucleoside monophosphate metabolic process |
| Module 006 | GO:0046700 | 2.870868238 | 2.44E-07 | 6.84E-05 | heterocycle catabolic process |
| Module 006 | GO:0044270 | 2.858443786 | 2.68E-07 | 6.84E-05 | cellular nitrogen compound catabolic process |
| Module 006 | GO:0090304 | 2.839481555 | 0.00023 | 0.01093126 | nucleic acid metabolic process |
| Module 006 | GO:0009126 | 2.83429784 | 0.00093 | 0.034944906 | purine nucleoside monophosphate metabolic process |
| Module 006 | GO:0009141 | 2.780869104 | 0.0011 | 0.040019941 | nucleoside triphosphate metabolic process |
| Module 006 | GO:0006396 | 2.750068587 | 0.0002 | 0.00947933 | RNA processing |
| Module 006 | GO:0010608 | 2.71472189 | 0.00138 | 0.039959699 | posttranscriptional regulation of gene expression |
| Module 006 | GO:1901361 | 2.710882409 | 1.15E-06 | 0.0001628 | organic cyclic compound catabolic process |
| Module 006 | GO:0019439 | 2.709508145 | 1.59E-06 | 0.000202592 | aromatic compound catabolic process |
| Module 006 | GO:0034660 | 2.379935841 | 0.00078 | 0.031824809 | ncRNA metabolic process |
| Module 006 | GO:1901575 | 2.353960869 | 0.00011 | 0.006247523 | organic substance catabolic process |
| Module 006 | GO:0055086 | 2.197876388 | 0.00146 | 0.04065016 | nucleobase-containing small molecule metabolic process |
| Module 006 | GO:0022411 | 2.160348306 | 0.00091 | 0.036179006 | cellular component disassembly |
| Module 006 | GO:0009057 | 1.927229347 | 9.37E-05 | 0.004720425 | macromolecule catabolic process |
| Module 006 | GO:0044238 | 1.726605932 | 1.95E-05 | 0.001420545 | primary metabolic process |
| Module 006 | GO:0044271 | 1.53913389 | 1.17E-05 | 0.001026151 | cellular nitrogen compound biosynthetic process |
| Module 006 | GO:0009059 | 1.44675648 | 0.00014 | 0.008670751 | macromolecule biosynthetic process |
| Module 006 | GO:0009058 | 1.422536043 | 0.00019 | 0.011414552 | biosynthetic process |

**Supplementary Table 6: GO Terms Enriched in Non-Exponentiated ER+ Breast Cancer Networks Using Consensus Method**

| Modules | GOID | OddsRatio | pval | adjPval | GO Terms |
|---|---|---|---|---|---|
| Module 002 | GO:0050911 | 2.831981601 | 2.92E-05 | 0.04288418 | detection of chemical stimulus involved in sensory perception of smell |
| Module 002 | GO:0010826 | Inf | 0.000606504 | 0.02929365 | negative regulation of centrosome duplication |
| Module 002 | GO:2001162 | Inf | 0.00069458 | 0.02461417 | positive regulation of histone H3-K79 methylation |
| Module 002 | GO:0033128 | 184.3513514 | 0.000349941 | 0.03499512 | negative regulation of histone phosphorylation |
| Module 002 | GO:0010847 | 92.16891892 | 0.000694878 | 0.04019346 | regulation of chromatin assembly |
| Module 002 | GO:0006353 | 87.37179487 | 0.000580802 | 0.03806379 | DNA-templated transcription, termination |
| Module 002 | GO:2001168 | 80.11176471 | 0.000912485 | 0.04358204 | positive regulation of histone H2B ubiquitination |
| Module 002 | GO:0007076 | 58.44460227 | 3.84E-05 | 0.00552536 | mitotic chromosome condensation |
| Module 002 | GO:0150093 | 39.95669291 | 0.000211722 | 0.02573387 | amyloid-beta clearance by transcytosis |
| Module 002 | GO:0085020 | 36.71195652 | 0.000269823 | 0.02016572 | protein K6-linked ubiquitination |
| Module 002 | GO:0032402 | 32.21176471 | 0.000191063 | 0.01817344 | melanosome transport |
| Module 002 | GO:0051905 | 28.41799308 | 0.000264474 | 0.02002446 | establishment of pigment granule localization |
| Module 002 | GO:0031167 | 27.89127424 | 0.000589553 | 0.0227916 | rRNA methylation |
| Module 002 | GO:0031440 | 24.2875 | 0.000489847 | 0.0377936 | regulation of mRNA 3'-end processing |
| Module 002 | GO:0045132 | 15.49092971 | 0.000205737 | 0.01817344 | meiotic chromosome segregation |
| Module 002 | GO:0032465 | 13.0928382 | 0.000403178 | 0.03499512 | regulation of cytokinesis |
| Module 002 | GO:0045740 | 12.45092593 | 5.75E-06 | 0.00099963 | positive regulation of DNA replication |
| Module 002 | GO:1901673 | 11.46666667 | 0.000864562 | 0.0294124 | regulation of mitotic spindle assembly |
| Module 002 | GO:0010824 | 9.936990363 | 0.000333522 | 0.01575892 | regulation of centrosome duplication |
| Module 002 | GO:0030433 | 9.724939467 | 1.73E-06 | 0.00024522 | ubiquitin-dependent ERAD pathway |
| Module 002 | GO:1904886 | 9.314583333 | 0.001653278 | 0.04848666 | beta-catenin destruction complex disassembly |
| Module 002 | GO:0000398 | 8.419510452 | 4.28E-05 | 0.00552536 | mRNA splicing, via spliceosome |
| Module 002 | GO:1903312 | 7.904988124 | 0.000884675 | 0.03766726 | negative regulation of mRNA metabolic process |
| Module 002 | GO:0000375 | 7.85195748 | 6.79E-06 | 0.00262851 | RNA splicing, via transesterification reactions |
| Module 002 | GO:1902275 | 7.610094851 | 0.00078657 | 0.04152264 | regulation of chromatin organization |
| Module 002 | GO:0006323 | 7.362528147 | 9.03E-05 | 0.01195992 | DNA packaging |
| Module 002 | GO:0090503 | 6.907051282 | 0.001459845 | 0.04917516 | RNA phosphodiester bond hydrolysis, exonucleolytic |
| Module 002 | GO:0042795 | 6.717131474 | 0.000481954 | 0.04956715 | snRNA transcription by RNA polymerase II |
| Module 002 | GO:0098781 | 6.289077213 | 9.97E-05 | 0.00628223 | ncRNA transcription |
| Module 002 | GO:0060964 | 6.274537037 | 0.00062064 | 0.04019346 | regulation of gene silencing by miRNA |
| Module 002 | GO:0040029 | 5.970666667 | 0.000791504 | 0.04122416 | regulation of gene expression, epigenetic |
| Module 002 | GO:0060968 | 5.967617968 | 0.000301072 | 0.03447273 | regulation of gene silencing |
| Module 002 | GO:0016073 | 5.89469046 | 0.000149111 | 0.00874613 | snRNA metabolic process |
| Module 002 | GO:0000280 | 5.652983496 | 0.000164549 | 0.02153242 | nuclear division |
| Module 002 | GO:1902850 | 5.61489899 | 2.44E-06 | 0.00031881 | microtubule cytoskeleton organization involved in mitosis |
| Module 002 | GO:0072395 | 5.495675044 | 0.00052828 | 0.03190812 | signal transduction involved in cell cycle checkpoint |

| | | | | | |
|---|---|---|---|---|---|
| Module 002 | GO:0051276 | 5.481524459 | 6.69E-06 | 0.00112245 | chromosome organization |
| Module 002 | GO:0051865 | 5.464351379 | 0.000237632 | 0.01224884 | protein autoubiquitination |
| Module 002 | GO:0006368 | 5.397094431 | 0.001484213 | 0.0467527 | transcription elongation from RNA polymerase II promoter |
| Module 002 | GO:0034502 | 5.300909091 | 0.000702316 | 0.03041697 | protein localization to chromosome |
| Module 002 | GO:0007098 | 4.989159689 | 8.27E-05 | 0.00541322 | centrosome cycle |
| Module 002 | GO:1901796 | 4.788340661 | 4.89E-05 | 0.00582726 | regulation of signal transduction by p53 class mediator |
| Module 002 | GO:0070507 | 4.777658046 | 1.27E-06 | 0.00019644 | regulation of microtubule cytoskeleton organization |
| Module 002 | GO:0051303 | 4.682422969 | 0.001306935 | 0.04194521 | establishment of chromosome localization |
| Module 002 | GO:0098813 | 4.511390683 | 1.01E-05 | 0.0010357 | nuclear chromosome segregation |
| Module 002 | GO:0071426 | 4.113276451 | 0.000644638 | 0.03743862 | ribonucleoprotein complex export from nucleus |
| Module 002 | GO:1903052 | 4.104115479 | 0.000670033 | 0.03122354 | positive regulation of proteolysis involved in cellular protein catabolic process |
| Module 002 | GO:0006403 | 4.02335319 | 3.11E-05 | 0.00426835 | RNA localization |
| Module 002 | GO:0016055 | 4.01241412 | 6.11E-05 | 0.00524377 | Wnt signaling pathway |
| Module 002 | GO:0006807 | 4.004926486 | 9.36E-08 | 0.00010729 | nitrogen compound metabolic process |
| Module 002 | GO:0001736 | 3.965752612 | 0.000439036 | 0.02987354 | establishment of planar polarity |
| Module 002 | GO:0071704 | 3.870481928 | 5.31E-06 | 0.002431 | organic substance metabolic process |
| Module 002 | GO:0007051 | 3.782102273 | 0.000348301 | 0.01601244 | spindle organization |
| Module 002 | GO:0090304 | 3.43951112 | 2.21E-05 | 0.00334352 | nucleic acid metabolic process |
| Module 002 | GO:0006139 | 3.389423077 | 0.000342266 | 0.03311423 | nucleobase-containing compound metabolic process |
| Module 002 | GO:0045787 | 3.343847926 | 0.000872559 | 0.0477968 | positive regulation of cell cycle |
| Module 002 | GO:0031329 | 3.208105911 | 0.000468833 | 0.01993711 | regulation of cellular catabolic process |
| Module 002 | GO:1903320 | 3.10128655 | 0.000399701 | 0.0174331 | regulation of protein modification by small protein conjugation or removal |
| Module 002 | GO:0043632 | 3.088023306 | 1.31E-07 | 4.47E-05 | modification-dependent macromolecule catabolic process |
| Module 002 | GO:0044419 | 2.983101648 | 0.000117474 | 0.01472825 | interspecies interaction between organisms |
| Module 002 | GO:0050658 | 2.929799134 | 0.001045734 | 0.04016473 | RNA transport |
| Module 002 | GO:0016579 | 2.918952224 | 0.000700544 | 0.0270341 | protein deubiquitination |
| Module 002 | GO:0006725 | 2.911228972 | 4.78E-09 | 7.22E-06 | cellular aromatic compound metabolic process |
| Module 002 | GO:0070647 | 2.863240183 | 9.82E-05 | 0.01498627 | protein modification by small protein conjugation or removal |
| Module 002 | GO:0046483 | 2.843836491 | 1.22E-08 | 9.24E-06 | heterocycle metabolic process |
| Module 002 | GO:1901360 | 2.612914572 | 1.57E-07 | 6.40E-05 | organic cyclic compound metabolic process |
| Module 002 | GO:0044238 | 2.585441541 | 9.43E-05 | 0.01965511 | primary metabolic process |
| Module 002 | GO:0030163 | 2.542841629 | 0.000179952 | 0.01020327 | protein catabolic process |
| Module 002 | GO:0010608 | 2.526099707 | 7.80E-05 | 0.00591141 | posttranscriptional regulation of gene expression |
| Module 002 | GO:0070727 | 2.51394905 | 5.05E-06 | 0.00168873 | cellular macromolecule localization |
| Module 002 | GO:0051301 | 2.512596899 | 0.000209188 | 0.01147835 | cell division |
| Module 002 | GO:0034641 | 2.49869723 | 3.85E-06 | 0.00109455 | cellular nitrogen compound metabolic process |
| Module 002 | GO:0051603 | 2.465111851 | 0.000148647 | 0.00874613 | proteolysis involved in cellular protein catabolic process |
| Module 002 | GO:0051640 | 2.445587728 | 0.000170719 | 0.01731577 | organelle localization |

| | | | | | |
|---|---|---|---|---|---|
| Module 002 | GO:0044237 | 2.350177305 | 0.000326052 | 0.01753226 | cellular metabolic process |
| Module 002 | GO:0030031 | 2.339175018 | 0.000886298 | 0.0477968 | cell projection assembly |
| Module 002 | GO:0009057 | 2.276372934 | 0.000936344 | 0.0340642 | macromolecule catabolic process |
| Module 002 | GO:0042886 | 2.205253333 | 2.70E-05 | 0.00584635 | amide transport |
| Module 002 | GO:0051171 | 2.113711002 | 8.12E-05 | 0.00694481 | regulation of nitrogen compound metabolic process |
| Module 002 | GO:0008104 | 2.109911909 | 0.000724528 | 0.04037233 | protein localization |
| Module 002 | GO:0044260 | 1.962358731 | 8.99E-07 | 0.00018799 | cellular macromolecule metabolic process |
| Module 002 | GO:0051649 | 1.954846226 | 0.000644473 | 0.04809834 | establishment of localization in cell |
| Module 002 | GO:0080090 | 1.859346235 | 7.76E-05 | 0.00541322 | regulation of primary metabolic process |
| Module 002 | GO:0051173 | 1.821369176 | 0.000374574 | 0.02349734 | positive regulation of nitrogen compound metabolic process |
| Module 002 | GO:0071840 | 1.821092934 | 6.99E-05 | 0.01106943 | cellular component organization or biogenesis |
| Module 002 | GO:0043412 | 1.80230947 | 1.84E-07 | 5.78E-05 | macromolecule modification |
| Module 002 | GO:0036211 | 1.801698244 | 0.000570067 | 0.04019346 | protein modification process |
| Module 002 | GO:0044085 | 1.787743952 | 2.13E-05 | 0.00196117 | cellular component biogenesis |
| Module 002 | GO:0044267 | 1.754821852 | 0.000149207 | 0.01860613 | cellular protein metabolic process |
| Module 002 | GO:0009059 | 1.569623487 | 7.15E-05 | 0.00641202 | macromolecule biosynthetic process |
| Module 002 | GO:0009889 | 1.397056277 | 0.001493238 | 0.04938547 | regulation of biosynthetic process |
| Module 003 | GO:0010839 | 44.24350649 | 7.64E-06 | 0.00667631 | negative regulation of keratinocyte proliferation |
| Module 003 | GO:0016339 | 14.52702703 | 5.73E-05 | 0.03232065 | calcium-dependent cell-cell adhesion via plasma membrane cell adhesion molecules |
| Module 004 | GO:0031581 | 46.0832396 | 9.35E-05 | 0.03785165 | hemidesmosome assembly |
| Module 004 | GO:1901750 | 41.41538462 | 2.06E-05 | 0.02514476 | leukotriene D4 biosynthetic process |
| Module 004 | GO:0030220 | 24.77378243 | 5.01E-05 | 0.0476234 | platelet formation |
| Module 004 | GO:0034329 | 19.53188406 | 9.76E-05 | 0.03785165 | cell junction assembly |
| Module 004 | GO:0046486 | 7.372080089 | 9.39E-05 | 0.04740791 | glycerolipid metabolic process |
| Module 004 | GO:0060079 | 6.216649251 | 0.000102941 | 0.03798507 | excitatory postsynaptic potential |
| Module 004 | GO:1905114 | 5.338308663 | 4.80E-05 | 0.02514476 | cell surface receptor signaling pathway involved in cell-cell signaling |
| Module 004 | GO:0043547 | 3.172831581 | 5.11E-05 | 0.02514476 | positive regulation of GTPase activity |
| Module 004 | GO:0007308 | Inf | 6.54E-05 | 0.01354953 | oocyte construction |
| Module 004 | GO:0007314 | Inf | 6.54E-05 | 0.01354953 | oocyte anterior/posterior axis specification |
| Module 004 | GO:0030719 | Inf | 6.54E-05 | 0.01354953 | P granule organization |
| Module 004 | GO:0000478 | 218.1904762 | 9.89E-05 | 0.01087378 | endonucleolytic cleavage involved in rRNA processing |
| Module 004 | GO:0000460 | 81.79166667 | 0.000440343 | 0.03229185 | maturation of 5.8S rRNA |
| Module 004 | GO:0090305 | 9.939290086 | 6.71E-05 | 0.01087378 | nucleic acid phosphodiester bond hydrolysis |

**Supplementary Table 7: GO Terms Enriched in Non-Exponentiated ER+ Breast Cancer Networks Using ALPACA**

| Modules | GOID | OddsRatio | pval | adjPval | GO Terms |
|---|---|---|---|---|---|
| Module 001 | GO:0042340 | 138.989899 | 0.00016 | 0.02071 | keratan sulfate catabolic process |
| Module 001 | GO:0050911 | 3.977447101 | 2.77E-09 | 4.14E-06 | detection of chemical stimulus involved in sensory perception of smell |
| Module 001 | GO:0007606 | 3.295628939 | 5.00E-06 | 0.00317 | sensory perception of chemical stimulus |
| Module 001 | GO:0009593 | 3.142882642 | 0.00015 | 0.04924 | detection of chemical stimulus |
| Module 001 | GO:0050906 | 3.027194492 | 0.00022 | 0.04924 | detection of stimulus involved in sensory perception |
| Module 001 | GO:0051606 | 2.584152236 | 2.59E-06 | 0.00202 | detection of stimulus |
| Module 001 | GO:0007600 | 2.054040896 | 7.77E-05 | 0.04044 | sensory perception |
| Module 002 | GO:0030033 | 156.4090909 | 0.00012 | 0.01754 | microvillus assembly |
| Module 002 | GO:0051651 | 10.61472425 | 4.24E-05 | 0.01149 | maintenance of location in cell |
| Module 002 | GO:0045185 | 9.007954043 | 9.88E-05 | 0.02011 | maintenance of protein location |
| Module 002 | GO:1901796 | 7.570808236 | 2.39E-05 | 0.01149 | regulation of signal transduction by p53 class mediator |
| Module 002 | GO:0007098 | 6.24715975 | 9.06E-05 | 0.02802 | centrosome cycle |
| Module 002 | GO:0070507 | 4.432373531 | 0.00018 | 0.04558 | regulation of microtubule cytoskeleton organization |
| Module 002 | GO:0000819 | 4.343035772 | 0.00011 | 0.03902 | sister chromatid segregation |
| Module 002 | GO:0044770 | 3.864411765 | 0.00016 | 0.03534 | cell cycle phase transition |
| Module 002 | GO:0051726 | 2.902549383 | 2.82E-08 | 3.49E-05 | regulation of cell cycle |
| Module 002 | GO:0051301 | 2.823160173 | 0.0002 | 0.04749 | cell division |
| Module 002 | GO:0000278 | 2.57511658 | 1.71E-05 | 0.00996 | mitotic cell cycle |
| Module 002 | GO:0043632 | 2.490979107 | 5.30E-05 | 0.025 | modification-dependent macromolecule catabolic process |
| Module 002 | GO:0043412 | 2.434441706 | 8.59E-05 | 0.04699 | macromolecule modification |
| Module 002 | GO:0070647 | 2.159709619 | 8.29E-05 | 0.0438 | protein modification by small protein conjugation or removal |
| Module 002 | GO:0036211 | 1.884809411 | 3.17E-05 | 0.0339 | protein modification process |
| Module 002 | GO:0006793 | 1.66577988 | 0.00017 | 0.04735 | phosphorus metabolic process |
| Module 002 | GO:0044260 | 1.52037098 | 6.58E-05 | 0.02786 | cellular macromolecule metabolic process |
| Module 003 | GO:0007308 | Inf | 1.04E-05 | 0.00091 | oocyte construction |
| Module 003 | GO:0007314 | Inf | 1.04E-05 | 0.00091 | oocyte anterior/posterior axis specification |
| Module 003 | GO:0030719 | Inf | 1.04E-05 | 0.00091 | P granule organization |
| Module 003 | GO:0000478 | 138.6060606 | 0.00024 | 0.02289 | endonucleolytic cleavage involved in rRNA processing |
| Module 003 | GO:0007028 | 91.30232558 | 0.00037 | 0.0241 | cytoplasm organization |
| Module 003 | GO:0008595 | 58.0845666 | 0.00079 | 0.04142 | anterior/posterior axis specification, embryo |
| Module 003 | GO:0032012 | 39.80232558 | 0.00011 | 0.01318 | regulation of ARF protein signal transduction |
| Module 003 | GO:0090305 | 4.911436255 | 0.00041 | 0.04902 | nucleic acid phosphodiester bond hydrolysis |
| Module 004 | GO:1901750 | 65.51169591 | 5.15E-07 | 0.0008 | leukotriene D4 biosynthetic process |

| | | | | | |
|---|---|---|---|---|---|
| Module 004 | GO:0031581 | 52.78764479 | 6.33E-05 | 0.02129 | hemidesmosome assembly |
| Module 004 | GO:0034329 | 22.13453651 | 6.12E-05 | 0.02129 | cell junction assembly |
| Module 004 | GO:0014068 | 6.936839789 | 0.00013 | 0.03803 | positive regulation of phosphatidylinositol 3-kinase signaling |
| Module 004 | GO:0035023 | 6.167124542 | 1.39E-05 | 0.01586 | regulation of Rho protein signal transduction |
| Module 004 | GO:0043547 | 3.7064403 | 6.98E-05 | 0.03803 | positive regulation of GTPase activity |
| Module 004 | GO:0045937 | 2.151531222 | 0.00014 | 0.04744 | positive regulation of phosphate metabolic process |
| Module 004 | GO:0043085 | 2.095913871 | 4.20E-05 | 0.04744 | positive regulation of catalytic activity |
| Module 005 | GO:1903241 | Inf | 1.85E-05 | 0.00142 | U2-type prespliceosome assembly |
| Module 005 | GO:2001162 | Inf | 0.00097 | 0.03164 | positive regulation of histone H3-K79 methylation |
| Module 005 | GO:1904903 | 218.4761905 | 7.40E-05 | 0.00353 | ESCRT III complex disassembly |
| Module 005 | GO:2000286 | 215.1653543 | 0.00026 | 0.01445 | receptor internalization involved in canonical Wnt signaling pathway |
| Module 005 | GO:1901534 | 110.0803213 | 0.00037 | 0.0223 | positive regulation of hematopoietic progenitor cell differentiation |
| Module 005 | GO:0061952 | 93.57823129 | 0.00031 | 0.01134 | midbody abscission |
| Module 005 | GO:1901673 | 87.33333333 | 0.00036 | 0.01134 | regulation of mitotic spindle assembly |
| Module 005 | GO:0000389 | 71.71128609 | 0.00085 | 0.03358 | mRNA 3'-splice site recognition |
| Module 005 | GO:0150093 | 66.03855422 | 0.00077 | 0.02858 | amyloid-beta clearance by transcytosis |
| Module 005 | GO:0036258 | 59.51515152 | 0.00072 | 0.01959 | multivesicular body assembly |
| Module 005 | GO:0007080 | 58.86428571 | 3.29E-05 | 0.00209 | mitotic metaphase plate congression |
| Module 005 | GO:0021860 | 51.62264151 | 0.00124 | 0.03587 | pyramidal neuron development |
| Module 005 | GO:0050000 | 48.92417484 | 3.28E-06 | 0.00063 | chromosome localization |
| Module 005 | GO:1904874 | 43.54920635 | 0.0014 | 0.0455 | positive regulation of telomerase RNA localization to Cajal body |
| Module 005 | GO:0006403 | 43.44771242 | 0.00022 | 0.01164 | RNA localization |
| Module 005 | GO:0010824 | 42.20890937 | 0.00136 | 0.03066 | regulation of centrosome duplication |
| Module 005 | GO:0032506 | 40.88690476 | 0.00144 | 0.03066 | cytokinetic process |
| Module 005 | GO:0090224 | 37.37414966 | 0.00171 | 0.0313 | regulation of spindle organization |
| Module 005 | GO:0030199 | 36.33333333 | 0.0018 | 0.0313 | collagen fibril organization |
| Module 005 | GO:0045653 | 22.48418248 | 1.17E-05 | 0.00177 | negative regulation of megakaryocyte differentiation |
| Module 005 | GO:0000281 | 21.76190476 | 0.00473 | 0.04938 | mitotic cytokinesis |
| Module 005 | GO:0007032 | 21.40359094 | 0.00488 | 0.04938 | endosome organization |
| Module 005 | GO:0009451 | 21.3828125 | 0.00491 | 0.04938 | RNA modification |
| Module 005 | GO:0080009 | 19.73183813 | 0.00105 | 0.03575 | mRNA methylation |
| Module 005 | GO:0006369 | 19.25234522 | 0.00074 | 0.02858 | termination of RNA polymerase II transcription |
| Module 005 | GO:0007059 | 17.43055556 | 2.68E-05 | 0.00209 | chromosome segregation |
| Module 005 | GO:0000244 | 16.37090909 | 0.00156 | 0.04991 | spliceosomal tri-snRNP complex assembly |
| Module 005 | GO:0006413 | 12.36818182 | 0.00263 | 0.04183 | translational initiation |

| | | | | | |
|---|---|---|---|---|---|
| Module 005 | GO:0070125 | 10.96857373 | 3.50E-05 | 0.00253 | mitochondrial translational elongation |
| Module 005 | GO:0006335 | 10.743749 | 0.00022 | 0.01211 | DNA replication-dependent nucleosome assembly |
| Module 005 | GO:0070126 | 10.6745098 | 4.04E-05 | 0.00259 | mitochondrial translational termination |
| Module 005 | GO:0034080 | 9.957446809 | 0.00109 | 0.04685 | CENP-A containing nucleosome assembly |
| Module 005 | GO:0000398 | 9.794181818 | 6.61E-05 | 0.00516 | mRNA splicing, via spliceosome |
| Module 005 | GO:0006303 | 9.334713596 | 9.07E-05 | 0.00715 | double-strand break repair via nonhomologous end joining |
| Module 005 | GO:0000375 | 9.237049743 | 5.74E-07 | 0.00011 | RNA splicing, via transesterification reactions |
| Module 005 | GO:0140053 | 7.968534989 | 5.54E-05 | 0.00299 | mitochondrial gene expression |
| Module 005 | GO:0006614 | 7.766990291 | 0.0007 | 0.02506 | SRP-dependent cotranslational protein targeting to membrane |
| Module 005 | GO:0022613 | 7.626984127 | 2.57E-05 | 0.00335 | ribonucleoprotein complex biogenesis |
| Module 005 | GO:0016233 | 7.506684492 | 0.00092 | 0.03574 | telomere capping |
| Module 005 | GO:0072599 | 7.172330097 | 0.00098 | 0.03129 | establishment of protein localization to endoplasmic reticulum |
| Module 005 | GO:0051656 | 7.128580946 | 0.00395 | 0.04938 | establishment of organelle localization |
| Module 005 | GO:0006396 | 7.098951953 | 1.34E-06 | 0.00039 | RNA processing |
| Module 005 | GO:0019083 | 6.889922481 | 1.03E-07 | 3.28E-05 | viral transcription |
| Module 005 | GO:0000245 | 6.274906015 | 0.00077 | 0.0284 | spliceosomal complex assembly |
| Module 005 | GO:0071840 | 6.265051606 | 0.00183 | 0.04403 | cellular component organization or biogenesis |
| Module 005 | GO:0000184 | 6.180533394 | 0.00066 | 0.03181 | nuclear-transcribed mRNA catabolic process, nonsense-mediated decay |
| Module 005 | GO:0034660 | 5.715447154 | 0.00042 | 0.02068 | ncRNA metabolic process |
| Module 005 | GO:0044419 | 5.680284036 | 0.00382 | 0.04938 | interspecies interaction between organisms |
| Module 005 | GO:0016071 | 5.624009914 | 0.00019 | 0.01495 | mRNA metabolic process |
| Module 005 | GO:0046483 | 5.210132412 | 0.00434 | 0.04938 | heterocycle metabolic process |
| Module 005 | GO:0006807 | 5.110838947 | 1.20E-16 | 1.79E-13 | nitrogen compound metabolic process |
| Module 005 | GO:0006725 | 5.102433503 | 0.00476 | 0.04938 | cellular aromatic compound metabolic process |
| Module 005 | GO:0071826 | 5.06493994 | 0.00033 | 0.01497 | ribonucleoprotein complex subunit organization |
| Module 005 | GO:0060964 | 4.833593656 | 0.00022 | 0.01263 | regulation of gene silencing by miRNA |
| Module 005 | GO:0006518 | 4.760090703 | 0.00069 | 0.0225 | peptide metabolic process |
| Module 005 | GO:0042773 | 4.740643201 | 0.00123 | 0.04358 | ATP synthesis coupled electron transport |
| Module 005 | GO:0043604 | 4.682142857 | 0.00077 | 0.0225 | amide biosynthetic process |
| Module 005 | GO:0071704 | 4.674545455 | 6.59E-07 | 0.00012 | organic substance metabolic process |
| Module 005 | GO:0060968 | 4.630023822 | 0.00029 | 0.01465 | regulation of gene silencing |
| Module 005 | GO:0061418 | 4.586200291 | 0.00034 | 0.0134 | regulation of transcription from RNA polymerase II promoter in response to hypoxia |
| Module 005 | GO:0043043 | 4.581540597 | 0.00018 | 0.01189 | peptide biosynthetic process |
| Module 005 | GO:0006139 | 4.463951271 | 1.48E-15 | 1.87E-12 | nucleobase-containing compound metabolic process |
| Module 005 | GO:0006457 | 4.198887423 | 0.00015 | 0.0085 | protein folding |

| | | | | | |
|---|---|---|---|---|---|
| Module 005 | GO:0006612 | 4.061992695 | 0.0001 | 0.00614 | protein targeting to membrane |
| Module 005 | GO:1903706 | 4.023565262 | 0.00072 | 0.03646 | regulation of hemopoiesis |
| Module 005 | GO:0044265 | 3.834012497 | 0.00033 | 0.01357 | cellular macromolecule catabolic process |
| Module 005 | GO:0070498 | 3.80952381 | 0.00063 | 0.02277 | interleukin-1-mediated signaling pathway |
| Module 005 | GO:0032200 | 3.736308162 | 0.00037 | 0.01562 | telomere organization |
| Module 005 | GO:0006364 | 3.719687093 | 0.00129 | 0.04459 | rRNA processing |
| Module 005 | GO:0006402 | 3.709469325 | 0.00013 | 0.00574 | mRNA catabolic process |
| Module 005 | GO:0034641 | 3.65893559 | 0.00055 | 0.02345 | cellular nitrogen compound metabolic process |
| Module 005 | GO:0044260 | 3.61475074 | 0.00054 | 0.02215 | cellular macromolecule metabolic process |
| Module 005 | GO:0010608 | 3.472089314 | 0.00022 | 0.01222 | posttranscriptional regulation of gene expression |
| Module 005 | GO:0035194 | 3.466582198 | 0.00195 | 0.04985 | posttranscriptional gene silencing by RNA |
| Module 005 | GO:0009144 | 3.463735457 | 0.00116 | 0.03954 | purine nucleoside triphosphate metabolic process |
| Module 005 | GO:0043603 | 3.421206484 | 0.00054 | 0.02595 | cellular amide metabolic process |
| Module 005 | GO:1903311 | 3.363651182 | 0.00089 | 0.03012 | regulation of mRNA metabolic process |
| Module 005 | GO:0009059 | 3.323554748 | 5.63E-07 | 0.00012 | macromolecule biosynthetic process |
| Module 005 | GO:0043488 | 3.270196801 | 0.00109 | 0.03362 | regulation of mRNA stability |
| Module 005 | GO:0016579 | 3.098810037 | 0.00062 | 0.0239 | protein deubiquitination |
| Module 005 | GO:0006259 | 2.829314421 | 0.00042 | 0.01835 | DNA metabolic process |
| Module 005 | GO:0043624 | 2.816294479 | 0.00143 | 0.04739 | cellular protein complex disassembly |
| Module 005 | GO:0072594 | 2.809789069 | 4.80E-05 | 0.00265 | establishment of protein localization to organelle |
| Module 005 | GO:0044237 | 2.615214846 | 9.11E-08 | 3.60E-05 | cellular metabolic process |
| Module 005 | GO:1901360 | 2.600073506 | 7.99E-06 | 0.00103 | organic cyclic compound metabolic process |
| Module 005 | GO:0034645 | 2.433374844 | 0.00197 | 0.04714 | cellular macromolecule biosynthetic process |
| Module 005 | GO:0044238 | 2.384882787 | 1.12E-05 | 0.00125 | primary metabolic process |
| Module 005 | GO:0090304 | 2.354765025 | 3.07E-17 | 2.46E-14 | nucleic acid metabolic process |
| Module 005 | GO:1901566 | 2.353419483 | 0.00079 | 0.02692 | organonitrogen compound biosynthetic process |
| Module 005 | GO:0070647 | 2.322818668 | 1.73E-05 | 0.00129 | protein modification by small protein conjugation or removal |
| Module 005 | GO:0043170 | 2.241541881 | 0.00014 | 0.00993 | macromolecule metabolic process |
| Module 005 | GO:0006886 | 2.159714834 | 0.00018 | 0.00984 | intracellular protein transport |
| Module 005 | GO:0044270 | 2.153936517 | 8.10E-05 | 0.00418 | cellular nitrogen compound catabolic process |
| Module 005 | GO:0046700 | 2.153936517 | 8.10E-05 | 0.00418 | heterocycle catabolic process |
| Module 005 | GO:0044403 | 2.139898002 | 6.26E-05 | 0.0038 | symbiont process |
| Module 005 | GO:0019439 | 2.112253541 | 0.00011 | 0.00539 | aromatic compound catabolic process |
| Module 005 | GO:1901361 | 2.018266589 | 0.00025 | 0.01061 | organic cyclic compound catabolic process |
| Module 005 | GO:1901576 | 1.896218447 | 5.62E-06 | 0.00069 | organic substance biosynthetic process |

| | | | | | |
|---|---|---|---|---|---|
| Module 005 | GO:0019219 | 1.834447755 | 0.00034 | 0.01645 | regulation of nucleobase-containing compound metabolic process |
| Module 005 | GO:0070727 | 1.803914944 | 0.00078 | 0.0318 | cellular macromolecule localization |
| Module 005 | GO:0051649 | 1.794834219 | 0.00055 | 0.02532 | establishment of localization in cell |
| Module 005 | GO:0044271 | 1.766210338 | 9.61E-05 | 0.0076 | cellular nitrogen compound biosynthetic process |
| Module 005 | GO:0042886 | 1.743599236 | 3.04E-05 | 0.0018 | amide transport |
| Module 005 | GO:0016070 | 1.647229543 | 0.00101 | 0.0375 | RNA metabolic process |
| Module 005 | GO:0034654 | 1.600568408 | 0.00022 | 0.01211 | nucleobase-containing compound biosynthetic process |
| Module 005 | GO:0097659 | 1.591550563 | 0.0003 | 0.01465 | nucleic acid-templated transcription |
| Module 005 | GO:2001141 | 1.536489904 | 0.00079 | 0.02737 | regulation of RNA biosynthetic process |
| Module 005 | GO:0018130 | 1.455985944 | 0.00076 | 0.03166 | heterocycle biosynthetic process |
| Module 005 | GO:0009889 | 1.447992663 | 0.00123 | 0.03969 | regulation of biosynthetic process |
| Module 005 | GO:0019438 | 1.436536115 | 0.00081 | 0.03214 | aromatic compound biosynthetic process |
| Module 005 | GO:1901362 | 1.422875544 | 0.00138 | 0.04555 | organic cyclic compound biosynthetic process |
| Module 006 | GO:0050911 | 5.085688878 | 1.42E-05 | 0.00422 | detection of chemical stimulus involved in sensory perception of smell |
| Module 006 | GO:0009593 | 4.702233115 | 1.46E-05 | 0.00422 | detection of chemical stimulus |
| Module 006 | GO:0007606 | 4.207734573 | 8.13E-05 | 0.01395 | sensory perception of chemical stimulus |
| Module 006 | GO:0050906 | 4.125855277 | 9.69E-05 | 0.01395 | detection of stimulus involved in sensory perception |
| Module 007 | GO:0050911 | 3.77143993 | 3.44E-05 | 0.02204 | detection of chemical stimulus involved in sensory perception of smell |
| Module 007 | GO:0051606 | 3.354871795 | 0.00012 | 0.046 | detection of stimulus |

**Supplementary Table 8: GO Terms Enriched in Non-Exponentiated ER+ Breast Cancer Networks Using CRANE**

| Modules | GOID | OddsRatio | pval | adjPval | GO Terms |
|---|---|---|---|---|---|
| Module 001 | GO:0010909 | Inf | 0.00031 | 0.048385314 | positive regulation of heparan sulfate proteoglycan biosynthetic process |
| Module 001 | GO:0044334 | Inf | 0.00031 | 0.048385314 | canonical Wnt signaling pathway involved in positive regulation of epithelial to mesenchymal transition |
| Module 001 | GO:1902524 | Inf | 0.00031 | 0.048385314 | positive regulation of protein K48-linked ubiquitination |
| Module 001 | GO:0048227 | 218.4761905 | 7.40E-05 | 0.011029675 | plasma membrane to endosome transport |
| Module 001 | GO:0090596 | 4.533688748 | 0.00032 | 0.048385314 | sensory organ morphogenesis |
| Module 001 | GO:0032774 | 2.473535654 | 3.19E-05 | 0.008079325 | RNA biosynthetic process |
| Module 001 | GO:0097659 | 2.427487124 | 6.99E-06 | 0.00405537 | nucleic acid-templated transcription |
| Module 001 | GO:2001141 | 2.33627398 | 8.33E-06 | 0.00405537 | regulation of RNA biosynthetic process |
| Module 001 | GO:0016070 | 2.30920863 | 5.46E-05 | 0.02406678 | RNA metabolic process |
| Module 001 | GO:0051252 | 2.171179189 | 0.00022 | 0.044143668 | regulation of RNA metabolic process |
| Module 001 | GO:0034654 | 1.982653729 | 1.32E-05 | 0.004291135 | nucleobase-containing compound biosynthetic process |
| Module 001 | GO:0009889 | 1.891924211 | 9.18E-05 | 0.034376527 | regulation of biosynthetic process |
| Module 001 | GO:0034645 | 1.8910942 | 4.26E-05 | 0.010365101 | cellular macromolecule biosynthetic process |
| Module 001 | GO:1901362 | 1.863811707 | 2.07E-05 | 0.008079325 | organic cyclic compound biosynthetic process |
| Module 001 | GO:0019438 | 1.843733712 | 3.39E-05 | 0.008079325 | aromatic compound biosynthetic process |
| Module 001 | GO:0018130 | 1.835027354 | 3.78E-05 | 0.008079325 | heterocycle biosynthetic process |
| Module 001 | GO:0031326 | 1.822917 | 6.17E-05 | 0.011009981 | regulation of cellular biosynthetic process |
| Module 001 | GO:0010556 | 1.812738341 | 8.62E-05 | 0.011528416 | regulation of macromolecule biosynthetic process |
| Module 001 | GO:0044271 | 1.756184896 | 8.50E-05 | 0.011528416 | cellular nitrogen compound biosynthetic process |
| Module 001 | GO:0090304 | 1.651242107 | 0.0004 | 0.047717527 | nucleic acid metabolic process |
| Module 002 | GO:0021589 | Inf | 5.74E-05 | 0.018569959 | cerebellum structural organization |
| Module 002 | GO:0046854 | 51.62030075 | 4.72E-05 | 0.014363415 | phosphatidylinositol phosphorylation |
| Module 002 | GO:0019883 | 33.86528926 | 0.00019 | 0.022884064 | antigen processing and presentation of endogenous antigen |
| Module 002 | GO:0036507 | 26.80980392 | 0.00032 | 0.043183993 | protein demannosylation |
| Module 002 | GO:0030433 | 14.82826087 | 4.13E-05 | 0.018569959 | ubiquitin-dependent ERAD pathway |
| Module 002 | GO:0035967 | 7.090909091 | 0.00033 | 0.043183993 | cellular response to topologically incorrect protein |
| Module 002 | GO:0006986 | 7.065172054 | 0.00011 | 0.024437152 | response to unfolded protein |
| Module 002 | GO:0080135 | 6.3165511 | 8.07E-05 | 0.016493582 | regulation of cellular response to stress |
| Module 002 | GO:0033036 | 1.929972006 | 1.34E-06 | 0.00185829 | macromolecule localization |
| Module 003 | GO:0071391 | 26.06362217 | 1.16E-05 | 0.021384179 | cellular response to estrogen stimulus |
| Module 005 | GO:0006475 | 91.28619529 | 0.00053 | 0.045729584 | internal protein amino acid acetylation |
| Module 005 | GO:0033211 | 83.58282209 | 3.33E-05 | 0.01428695 | adiponectin-activated signaling pathway |
| Module 005 | GO:0033540 | 37.68595041 | 0.00013 | 0.024602759 | fatty acid beta-oxidation using acyl-CoA oxidase |
| Module 005 | GO:0006388 | 32.24586288 | 0.00023 | 0.045706792 | tRNA splicing, via endonucleolytic cleavage and ligation |
| Module 005 | GO:0045653 | 31.88344988 | 0.00021 | 0.024602759 | negative regulation of megakaryocyte differentiation |
| Module 005 | GO:0051131 | 15.85054428 | 0.00029 | 0.022861463 | chaperone-mediated protein complex assembly |
| Module 005 | GO:0006890 | 14.49465812 | 1.04E-08 | 8.87E-06 | retrograde vesicle-mediated transport, Golgi to ER |

| | | | | | |
|---|---|---|---|---|---|
| Module 005 | GO:0006302 | 12.3071244 | 0.00013 | 0.01598865 | double-strand break repair |
| Module 005 | GO:0006625 | 12.11606825 | 0.0001 | 0.024602759 | protein targeting to peroxisome |
| Module 005 | GO:0072662 | 12.11606825 | 0.0001 | 0.024602759 | protein localization to peroxisome |
| Module 005 | GO:0019395 | 8.996167063 | 0.00037 | 0.036847227 | fatty acid oxidation |
| Module 005 | GO:0070125 | 8.244412879 | 4.71E-08 | 6.47E-05 | mitochondrial translational elongation |
| Module 005 | GO:0070126 | 7.255045375 | 5.11E-07 | 0.000233622 | mitochondrial translational termination |
| Module 005 | GO:0006521 | 6.144444444 | 0.00029 | 0.022861463 | regulation of cellular amino acid metabolic process |
| Module 005 | GO:0140053 | 5.724699828 | 3.31E-07 | 0.000227164 | mitochondrial gene expression |
| Module 005 | GO:0061418 | 5.586052563 | 0.0002 | 0.019244353 | regulation of transcription from RNA polymerase II promoter in response to hypoxia |
| Module 005 | GO:0006303 | 5.561929751 | 0.00055 | 0.034721246 | double-strand break repair via nonhomologous end joining |
| Module 005 | GO:0031146 | 5.203666777 | 0.00072 | 0.041535525 | SCF-dependent proteasomal ubiquitin-dependent protein catabolic process |
| Module 005 | GO:0044106 | 5.117650273 | 0.00078 | 0.041854283 | cellular amine metabolic process |
| Module 005 | GO:0002479 | 5.08425467 | 0.00017 | 0.020248534 | antigen processing and presentation of exogenous peptide antigen via MHC class I, TAP-dependent |
| Module 005 | GO:1902036 | 4.950769555 | 0.00094 | 0.046991108 | regulation of hematopoietic stem cell differentiation |
| Module 005 | GO:0006402 | 4.470871594 | 0.00037 | 0.027457171 | mRNA catabolic process |
| Module 005 | GO:0006412 | 4.361890127 | 1.39E-05 | 0.002733393 | translation |
| Module 005 | GO:1903311 | 4.283334681 | 0.00099 | 0.046991108 | regulation of mRNA metabolic process |
| Module 005 | GO:0009199 | 4.26125 | 1.91E-05 | 0.004443627 | ribonucleoside triphosphate metabolic process |
| Module 005 | GO:0043624 | 4.195583596 | 5.66E-06 | 0.001474625 | cellular protein complex disassembly |
| Module 005 | GO:0009144 | 4.10875 | 2.75E-05 | 0.004978546 | purine nucleoside triphosphate metabolic process |
| Module 005 | GO:0038095 | 4.042628894 | 0.00073 | 0.041535525 | Fc-epsilon receptor signaling pathway |
| Module 005 | GO:0034660 | 3.829899809 | 9.48E-05 | 0.012678446 | ncRNA metabolic process |
| Module 005 | GO:0009167 | 3.773189161 | 0.00036 | 0.031379823 | purine ribonucleoside monophosphate metabolic process |
| Module 005 | GO:0010608 | 3.520584554 | 4.10E-05 | 0.013111518 | posttranscriptional regulation of gene expression |
| Module 005 | GO:0009259 | 3.147288952 | 0.00023 | 0.021736241 | ribonucleotide metabolic process |
| Module 005 | GO:0000398 | 3.11443292 | 0.00039 | 0.032737181 | mRNA splicing, via spliceosome |
| Module 005 | GO:0062012 | 3.095198773 | 0.00099 | 0.046991108 | regulation of small molecule metabolic process |
| Module 005 | GO:0072594 | 3.091155482 | 4.52E-05 | 0.007747196 | establishment of protein localization to organelle |
| Module 005 | GO:0000375 | 2.881422925 | 3.39E-05 | 0.005409063 | RNA splicing, via transesterification reactions |
| Module 005 | GO:0043687 | 2.871473901 | 7.54E-05 | 0.013111518 | post-translational protein modification |
| Module 005 | GO:0034641 | 2.57741134 | 3.99E-05 | 0.007854393 | cellular nitrogen compound metabolic process |
| Module 005 | GO:0022411 | 2.386008599 | 0.00041 | 0.027853926 | cellular component disassembly |
| Module 005 | GO:0034655 | 2.244148406 | 0.00042 | 0.027853926 | nucleobase-containing compound catabolic process |
| Module 005 | GO:0070647 | 2.1349663 | 2.25E-06 | 0.001222269 | protein modification by small protein conjugation or removal |
| Module 005 | GO:0006886 | 1.88949453 | 0.00077 | 0.04849709 | intracellular protein transport |
| Module 005 | GO:0009058 | 1.877207682 | 5.67E-05 | 0.008341906 | biosynthetic process |
| Module 005 | GO:0042886 | 1.746151773 | 3.65E-05 | 0.005409063 | amide transport |
| Module 005 | GO:0051649 | 1.717806064 | 0.00021 | 0.022378026 | establishment of localization in cell |
| Module 005 | GO:0006807 | 1.658208408 | 0.00015 | 0.017803306 | nitrogen compound metabolic process |

| | | | | | |
|---|---|---|---|---|---|
| Module 005 | GO:0070727 | 1.630297244 | 0.00055 | 0.034721246 | cellular macromolecule localization |
| Module 005 | GO:0044237 | 1.61653282 | 4.64E-05 | 0.00758514 | cellular metabolic process |
| Module 005 | GO:1901576 | 1.505068936 | 0.0004 | 0.032737181 | organic substance biosynthetic process |
| Module 007 | GO:0090179 | 178.7272727 | 0.0001 | 0.008312652 | planar cell polarity pathway involved in neural tube closure |
| Module 007 | GO:0090177 | 138.989899 | 0.00016 | 0.008312652 | establishment of planar polarity involved in neural tube closure |
| Module 007 | GO:1905276 | 104.219697 | 0.00026 | 0.010688958 | regulation of epithelial tube formation |
| Module 007 | GO:0009311 | 33.73955774 | 0.00207 | 0.04237906 | oligosaccharide metabolic process |
| Module 007 | GO:0050732 | 30.43902439 | 0.00251 | 0.046750406 | negative regulation of peptidyl-tyrosine phosphorylation |
| Module 007 | GO:0045739 | 28.35743802 | 0.00287 | 0.047155297 | positive regulation of DNA repair |
| Module 007 | GO:0048546 | 27.72525253 | 0.00299 | 0.047155297 | digestive tract morphogenesis |
| Module 007 | GO:0001841 | 21.95826645 | 0.00053 | 0.015378805 | neural tube formation |
| Module 007 | GO:0030308 | 18.92361111 | 0.00011 | 0.008312652 | negative regulation of cell growth |
| Module 007 | GO:0050866 | 17.22911392 | 0.00016 | 0.008312652 | negative regulation of cell activation |
| Module 007 | GO:0001736 | 16.96149068 | 0.00108 | 0.027765599 | establishment of planar polarity |
| Module 007 | GO:0022408 | 13.61238761 | 0.002 | 0.04237906 | negative regulation of cell-cell adhesion |
| Module 007 | GO:0030155 | 7.621028307 | 0.00039 | 0.013311106 | regulation of cell adhesion |
